# Cost-effectiveness of Precision Guided SIT for Control of *Anopheles gambiae* in the Upper River Region, The Gambia

**DOI:** 10.1101/2023.07.20.549762

**Authors:** William A.C. Gendron, Robyn Raban, Agastya Mondal, Héctor M. Sánchez C., Andrea Smidler, David Zilberman, Patrick G. Ilboudo, Umberto D’Alessandro, John M. Marshall, Omar S. Akbari

**Author notes:** To whom correspondence should be addressed: Omar S. Akbari, Ph.D., School of Biological Sciences, Department of Cell and Developmental Biology, University of California, San Diego, La Jolla, CA 92093, USA.

## Abstract

Precision-guided sterile insect technique (pgSIT) is an extremely promising vector control intervention that can reduce and potentially eliminate the unacceptable malaria burden, particularly in sub-Saharan Africa. The deployment of pgSIT shows the greatest promise and does not have a near peer competitor in the fight to eradicate malaria. Here we explore the cost effectiveness of using this approach in Africa using mathematical modeling and economical analysis. Overall, we find that pgSIT represents a cost-effective and promising approach to *A. gambiae* control in The Gambia, with the potential to deliver significant economic and social benefits.

**Summary:** Precision-guided sterile insect technique (pgSIT) is an extremely promising vector control intervention that can reduce and potentially eliminate the unacceptable malaria burden, particularly in sub-Saharan Africa. pgSIT is a safe, innovative, and highly targeted approach to mosquito control that combines the principles of the sterile insect technique (SIT) with advanced state-of-the-art technologies of genetic engineering (Akbari et al. 2023; M. Li et al. 2021; Kandul, Liu, and Akbari 2021; Kandul et al. 2019, 2022). The technique involves inundative releases of genetically sterile male mosquitoes into the environment to mate with their wild counterparts, sterilizing them in the process. After multiple releases, this method can suppress, and even eradicate pest populations without the use of chemical pesticides or other less specific agents. The use of pgSIT has the potential to be a lasting, safe, cost-effective, sustainable, and environmentally friendly method to suppress a target species, acting as a chemical-free, species-specific insecticide.

Before this work is considered for field application, however, we need to have a robust, data-driven modeling framework that will accurately predict the outcome of pgSIT release scenarios for malaria control, and we need an assessment of the costs and health benefits of implementing this technology in a region in Africa. This cost assessment evaluates the economic feasibility of both capacity building and establishing the infrastructure for a pgSIT facility in The Gambia to control the deadly malaria mosquito vector *Anopheles gambiae*. We focused on the Upper River region (URR) for three key reasons: (1) this area is known to have the highest per capita malaria rates in The Gambia (2) this region has more comprehensive historical and current data on malaria incidence and prevalence, malaria-associated healthcare and prevention costs, and human demographic data and (3) its location could feasibly demonstrate that a single pgSIT production facility can support phased suppression on a regional scale and then upon local extinction be repurposed to support vector suppression on a country or continent-wide scale.

The pgSIT treatment of the URR (∼2069 km^2^) is predicted to prevent approximately **230 deaths and about 48,000 sick days per yea**r. This estimate is based on a model for localized extinction of *A. gambiae*, reaching full epidemiological impact by the third year. There are multiple ways to calculate the value of this intervention monetarily, from the **value of statistical life (VSL) to quality adjusted life years (QALY), which ranges from 367 million to 880 million USD** saved in the first ten years of the facility being active. Other metrics such as willingness-to-pay (WTP), estimates the willingness of locals to contribute to malaria prevention financially, and estimates based on gross domestic product (**GDP) growth predict this model to save either 53 million or 551 million USD**, respectively, in the first ten years of intervention. This model assumes localized extinction of *A. gambiae* by the second year of intervention with repeated releases to maintain extinction despite seasonal mosquito migration from beyond the treatment area. Localized extinction, however, is expected to have a year-to-year suppression effect making it easier to suppress mosquito populations with reduced sterile male releases in subsequent years. It is, therefore, likely an underestimate of the costs and benefits of pgSIT sterile male releases. In later years, the release of sterile males from this facility could be redirected to new areas to expand the suppression region. Additionally, this facility would have a significant off-season where the facility is not producing sterile male *A. gambiae*. This off-season could, therefore, produce sterile males to suppress *A. gambiae* populations in other regions with seasonal malaria during the off-season in The Gambiae. It could even be used to mass produce pgSIT sterile males for other mosquito species, such as the dengue vector, *Aedes aegypti*, which have eggs that can be stored for many months and consequently can be stockpiled to aid in the suppression of dengue outbreaks, which are also common in the area. Initially, however, the off-season can be used to build local capacity for genetically engineered (GE) mosquitos, which would consist of training staff, optimizing procedures, and troubleshooting any issues that arose during the higher production phases.

The costs for this approach will vary as there are some unknowns and variability in the expected efficiencies of the facility, equipment, and procedures. In particular, mosquito survival rates and fecundity may vary more widely at scale and selection of the rearing equipment and protocols, and the mosquito sorting technology will dictate production levels and procedures. This variability is factored into the cost, and, therefore, a range of technologies and costs are considered. The **expected start-up costs range from 6 to 11.5 million USD**, which includes all necessary development, field trials, equipment, facility construction, staffing, and other establishment costs. The major upfront costs vary by facility size and capital equipment, much of which is dependent on mosquito rearing efficiencies. The annual estimated **operational costs range from approximately 315,000 USD,** depending on equipment selection and the size of the facility. Annually, this intervention costs less than **1 USD per person** to suppress mosquitoes in the URR and prevent malaria transmission, which is about **6% of a malaria prevention WTP** based on current interventions or **0.3-11%** when using the VSL or QALY metrics, with most variation from the VSL calculation method. This intervention **costs 15 to 124 USD (2022) to save one life-year and prevents malaria infections at 13 to 113 USD per case prevented,** making this method competitive with many current interventions. WTP demonstrates that this facility could be made locally sustainable long term by local funding.

Overall, the estimates of the capital and operational costs associated with the production of pgSIT sterile males, and the construction and management of the production facility indicate the cost savings associated with the annual decrease in morbidity and mortality (value of life) resulting from the use of pgSIT are significantly higher than the implementation costs. These estimates suggest that pgSIT represents a cost-effective and promising approach to *A. gambiae* control in The Gambia, with the potential to deliver significant economic and social benefits.

## Introduction

### Background

Mosquito-borne diseases account for more than 700 million human infections annually (Caraballo and King 2014)). These infections cause sickness and a significant number of deaths requiring investment in prevention and treatment. Malaria kills approximately 700,000 people a year, with African children under the age of five accounting for most malaria-related deaths (“Vector-Borne Diseases” n.d.). Progress has been made in disease prevention and treatment, which has reduced the numbers significantly, but further efforts are required to eliminate these mosquito-borne diseases.

Mosquito control is the cornerstone of mosquito-borne disease prevention. Long lasting insecticidal nets (LLINs) and indoor residual spraying (IRS) are the primary current tools to prevent indoor mosquito biting and resting (Hamel et al. 2011; Musoke et al. 2023; Pryce, Medley, and Choi 2022). These current strategies are insufficient to eradicate the mosquito population and could benefit from the synergistic use of genetic control technologies.

### Precision guided SIT

Precision guided sterile insect technique (pgSIT) combines the benefits of the established sterile insect technique (SIT) with the benefits of the precision and adaptability of genetic engineering. Despite success in other species, traditional SIT methods of using irradiation to sterilize fragile (by randomly fragmenting DNA) adult males have resulted in enormous fitness costs, which has precluded the use of SIT for malaria control. PgSIT has the advantage of using creative genetic engineering tools to precisely disrupt genes important for female viability and male fertility to produce highly fit and competitive sterile males. Moreover, pgSIT eggs can be hatched directly into the environment (e.g., dropped via drones) or alternatively released into water containers - or even small “just add water” containers-days later, only sterile males will emerge. These distribution procedures would require limited resources and training, which simplifies application at any site. Given that we can distribute eggs (as opposed to very fragile adult males, which is the requirement for other SIT, *Wolbachia* mediated suppression, and female killing technologies except for Ifegenia (Smidler, Pai, et al. 2023)), this also enables us to reach remote locations around the globe. Our technology allows the release of any mosquito life stage, which can benefit the entomological or epidemiological impact of the technology. Therefore, in contrast to current technologies, the pgSIT technology is:

1. **Adaptable:** The CRISPR/Cas9 system used to develop pgSIT has been successfully applied to a large number of insect species, including disease vectors (Kistler, Vosshall, and Matthews 2015; Gantz et al. 2015; Reid and O’Brochta 2016; Hammond et al. 2015), agricultural pests (Y. Li et al. 2016; Awata et al. 2015; Huang et al. 2016), beneficial insects (Ma et al. 2014; X. Li et al. 2015) and valuable research species (Bassett et al. 2013; Gratz et al. 2013) as well as a large number of non-insect species. Additionally, this binary approach only requires two components: Cas9 and gRNAs, which greatly simplifies its application to vector control in comparison to the current Release of Insects carrying a Dominant Lethal (RIDL) (Thomas et al. 2000) and *Wolbachia* systems (which remain mechanistically elusive), both of which are not available technologies for Malaria mosquitoes. The modular design of our system also permits easy modification of the components. To target different genes, the 23 bp gRNA target for CRISPR/Cas9 can be trivially replaced with a new gRNA targeting an alternate gene. The genes targeted for knockout can be interspecific, targeting a gene highly conserved between vector species or species specific. Likewise, this technology does not require drugs or chemicals, such as in the RIDL (Thomas et al. 2000), to rear insects in the laboratory, which are known to have a negative effect on mitochondria (Moullan et al. 2015; Chatzispyrou et al. 2015) and insect gut microflora (Kalghatgi et al. 2013) likely reducing released male fitness in the environment.
2. **Efficient and reproducible:** The efficiency of pgSIT relies on the exceedingly robust and precise cleavage rate of *wt* alleles by CRISPR Cas9/gRNA and the inability of insects to rescue mosaic loss-of-function phenotypes at the organismic level. Any CRISPR-based gene-drive system requires two actions: (1) cleavage of a target locus by Cas9/gRNA, and (2) homology-directed repair (HDR) of the induced cut locus to integrate a transgene carrying the Cas9/gRNA system. Our observations of multiple pgSIT systems engineered in multiple species suggest that Cas9/gRNA knockouts in somatic and germ tissues early during development ensure reproducible and complete penetrance of loss-of-function phenotypes in an adult insect (Kandul et al. 2018). In the target species, *A. gambiae*, we have been able to achieve 100% sterile male production using the pgSIT system (**Fig. 1**)(Smidler, Apte, et al. 2023). Moreover, given that pgSIT sterile males are developed using only laboratory strains with known sequences at targeted loci in a factory, then released, the genetic diversity in the wild will not affect the pgSIT technology, i.e., the males are already sterile. Therefore, the significant advantage of the proposed technology over any gene drive system is that pgSIT is not creating resistance alleles, not sampling naturally occurring alleles, and not relying on error-prone and unpredictable HDR pathways.
3. **Scalable:** A remarkable benefit of the pgSIT technology is that the homozygous Cas9 and homozygous gRNA lines can be healthily maintained separately and then crossed in a central rearing facility. The resulting progeny which are “dead-end” F_1_ trans-hemizygous CRISPR/Cas9 sterile male eggs can be easily distributed to any location for immediate hatching and release. pgSIT would eliminate the high cost (and error) associated with rearing and sexing mosquitoes near release locations as well as have the benefit of improved quality control and supply chain management from centralized manufacturing, especially for applications in resource-limited locations. It also eliminates the need to handle the fragile adult sterile males - which is a major limitation of related approaches (e.g. SIT, Wolbachia).
4. **Production efficient:** The pgSIT technology will provide a simple procedure to mass-produce sterile mosquito males. Aside from a recent alternate system we developed in *A. gambiae* termed inherited female elimination by genetically encoded nucleases to interrupt allele (Ifegenia)(Smidler, Pai, et al. 2023), currently, there are no other highly efficient, non-labor-intensive genetic methods to kill all female mosquitoes early in development (Papathanos et al. 2009). Sexing at the pupal or adult stages is highly labor-intensive, imprecise, or not feasible for large-scale production, or they are prohibitively expensive for most target disease-endemic countries and require infrastructure that is simply unreliable in resource-limited countries. For example, the Debug program at Verily Life Sciences has generated a state-of-the-art automated mosquito rearing and sexing facility for *Ae. aegypti* that achieves essentially 99.99% sexing accuracy for their population suppression *Wolbachia* strains. However, this approach is very rate limiting as each male sorted is a single male that is released (1x). For pgSIT, each female sorted results in 150X males produced as eggs making this approach far more scalable and efficient (150x). Moreover, these technologies require large and expensive facilities, and since mosquitoes are sexed at the adult stage, they cannot be shipped long distances. Many areas with the highest need for vector-borne disease interventions lack reliable electricity, water, supply chains, and other resources that are required to reliably produce mosquitoes with this technology and maintain these facilities. Therefore, this technology is unfortunately only feasible in developed countries, which have less epidemiological need for these technologies, and has not been adapted for malaria vectors. In our approach, the simple pgSIT system either kills or masculinizes all females, rendering all viable progeny to be sterile males that can be released as eggs (Kandul et al. 2018).
5. **Environmentally safe: insecticide-free and genetically non-invasive:** The pgSIT technology is an environment-friendly SIT, which is species-specific. Most malaria prevention technologies are insecticide-based, which are at high risk for failure due to insecticide resistance and can adversely impact beneficial insects, such as bees. pgSIT males are sterile, so their transgenes are not passed to the next generation and therefore do not persist in wild mosquito populations, which removes the long-term risk associated with synthetic genes in wild populations. While we would initially develop and test pgSIT in *A. gambiae*, there is evidence that other related malaria transmitting vectors can mate and produce hybrids with this species. That said, **inundative releases of sterile pgSIT *A. gambiae* males may result in suppression of other malaria transmitting species** - which would be a massive benefit to using this technology. This benefit would be unique to pgSIT, as suppression gene drives would require fertile hybrids to be produced, AND the gene drive target site would need 100% conservation (assuming resistant alleles are not generated, which is not possible) in that species for the drive to function - which would be very unlikely.
6. **Faster to market**: There is a regulatory precedent for GE SIT based technologies with established regulatory pipelines, and approval pathways and procedures. Other suppression technologies, particularly gene drive-based approaches, have highly uncertain regulatory requirements and pathways to market.
7. **Ecological evaluation of localized extinction and future purpose facility:** The application of pgSIT on a large scale provides the opportunity to evaluate downstream ecological effects of localized extinction of *A. gambiae* in a controlled manner. This would provide valuable data that could inform future decisions on the use of more invasive genetic suppression strategies. Additionally, the facility infrastructure could also be used and adapted to produce the next generations of genetically engineered mosquito suppression technologies. Scalable and affordable genetic technologies, like pgSIT, have the potential to be implemented by malaria endemic nations, which would give these mosquito suppression systems resilience to global issues along with economic benefits to these regions. This project consists of two objectives: (1) model native and simulated release mosquito ecology in The Gambia and (2) assess the health benefits and the economics of utilizing pgSIT in The Gambia.
8. **Novel and a sole competitor for improving malaria vector control:** Precision guided sterile insect technique (pgSIT) is the most viable of emerging and historic approaches to eradicate malaria. While there are several potential competitors, these currently exceed the technological capabilities in these fields or shift the valuation of the associated risks of the approaches. We will discuss each of these alternatives and explain why these are not peer competitor technologies to pgSIT.

a. **Current approaches**: While long lasting insecticidal nets (LLINs), indoor residual sprays (IRS) and prophylactic treatment with anti-malaria drugs have saved millions of lives since their implementation, these methods cannot completely eliminate malaria globally. These interventions still result in a significant number of malaria cases and increasing the scale of these interventions will not likely achieve elimination (Shretta et al. 2017). Achieving complete compliance for all of these techniques is nearly impossible, so a subset of the population is unprotected. These approaches have become less effective as mosquito insecticide and parasite drug resistance has become more widespread. Novel insecticides and drugs are in development, but resistance to these new chemicals will also occur. Mosquitoes are also changing their behavior to avoid these interventions, so more bites are now occurring outdoors or at times when people are not protected by LLINs. Therefore, increasing the coverage or intensity of current interventions is unlikely to eliminate malaria worldwide.
b. **Wide-scale insecticide use and environmental modification:** From 1946- 1951, the malaria endemic regions of the United States were saturated with DDT and water sources were either treated or removed on a massive scale to eliminate malaria mosquitoes. While successful at malaria elimination, when the WHO attempted a similar plan from 1955-1969, they failed to eliminate malaria. Mosquito resistance to DDT had developed rapidly and thwarted their efforts and a switch to the more integrated malaria control approach used today was adopted. Therefore, achieving malaria elimination with insecticides would likely require an environmentally devastating amount of insecticide and environmental modification, or novel insecticides will need new modes of action to be developed and their implementation closely managed. Insecticides would better serve malaria programs as one of many tools in an integrated management program as their abundant usage is toxic to humans and the environment and leads to their ineffectiveness.
c. **Malaria vaccines:** Malaria vaccines have long been in development to prevent severe disease and death in patients. Only two vaccines have been approved for use in preventing malaria, with the most recent showing 66% efficacy (“WHO Recommends R21/Matrix-M Vaccine for Malaria Prevention in Updated Advice on Immunization” n.d.). Vaccines have some fundamental limitations, however. High vaccine coverage is difficult in regions with limited infrastructure and census data. Current vaccines require cold chain transport and storage at 2-8 degrees Celsius, require boosters to maintain efficacy and specialized training for injection. Future vaccines may be thermostable and oral, but current vaccines have shortcomings in efficacy, storage and distribution that make them unable to achieve malaria elimination.
d. **Sterile Insect Technique (SIT):** Traditional SIT males are sterilized with radiation and then released. This approach is the closest competitor to pgSIT, but is inherently less efficient. Traditional SIT can only sterilize adult stage insects, so this technique can only support adult releases. Since the mode of sterilization in pgSIT occurs in early embryogenesis, any life stage can be released. Traditional SIT has to sex sort, irradiate, and transport the highly fragile adult mosquitoes, which decreases their survival and fitness. Additional cost is associated with rearing traditional SIT mosquitoes to adulthood in the laboratory, whereby pgSIT eggs save cost by being maintained in local water sources. The requirement to deliver adult male mosquitoes inherently incurs a 400-600 fold increase in the number of mosquitoes reared in the facility.
e. **Release of Insects carrying a Dominant Lethal gene (RIDL):** Another close competitor to pgSIT is the Release of Insects carrying a Dominant Lethal gene (RIDL), which relies on a gene that induces larval stage lethality in the absence of a drug. In the presence of the drug, the mosquitoes can be mass produced for release into the wild. These RIDL mosquitoes then mate with wild mosquitoes, and in the absence of the drug, produce inviable offspring. This approach has suppressed the dengue vector, *Ae. aegypti* in Brazil and other parts of the world(Carvalho et al. 2015; Garziera et al. 2017). While this approach is promising, a RIDL system has yet to be developed for any malaria vector. If an Anopheles RIDL system is developed, however, it would be years behind the development of pgSIT and likely have reduced efficacy (Kandul et al. 2019) and increased cost compared to pgSIT.
f. **Gene drive:** Gene drives are in development to make mosquito populations resistant to malaria or to suppress these populations. While this is a potentially powerful technology, there are risks and political and regulatory concerns that will make release of this technology more difficult than non-self propagating technologies, like pgSIT. There are also technical issues and uncertainties that make the performance and predictability of gene drives in the wild questionable. Many years of research will likely be needed to resolve these issues. Many gene drives are also not easily reversible like pgSIT systems, so they would be difficult to recall if unexpected effects occur. With pgSIT, halting releases will quickly remove the pgSIT individuals from the population.

**Fig 1.**
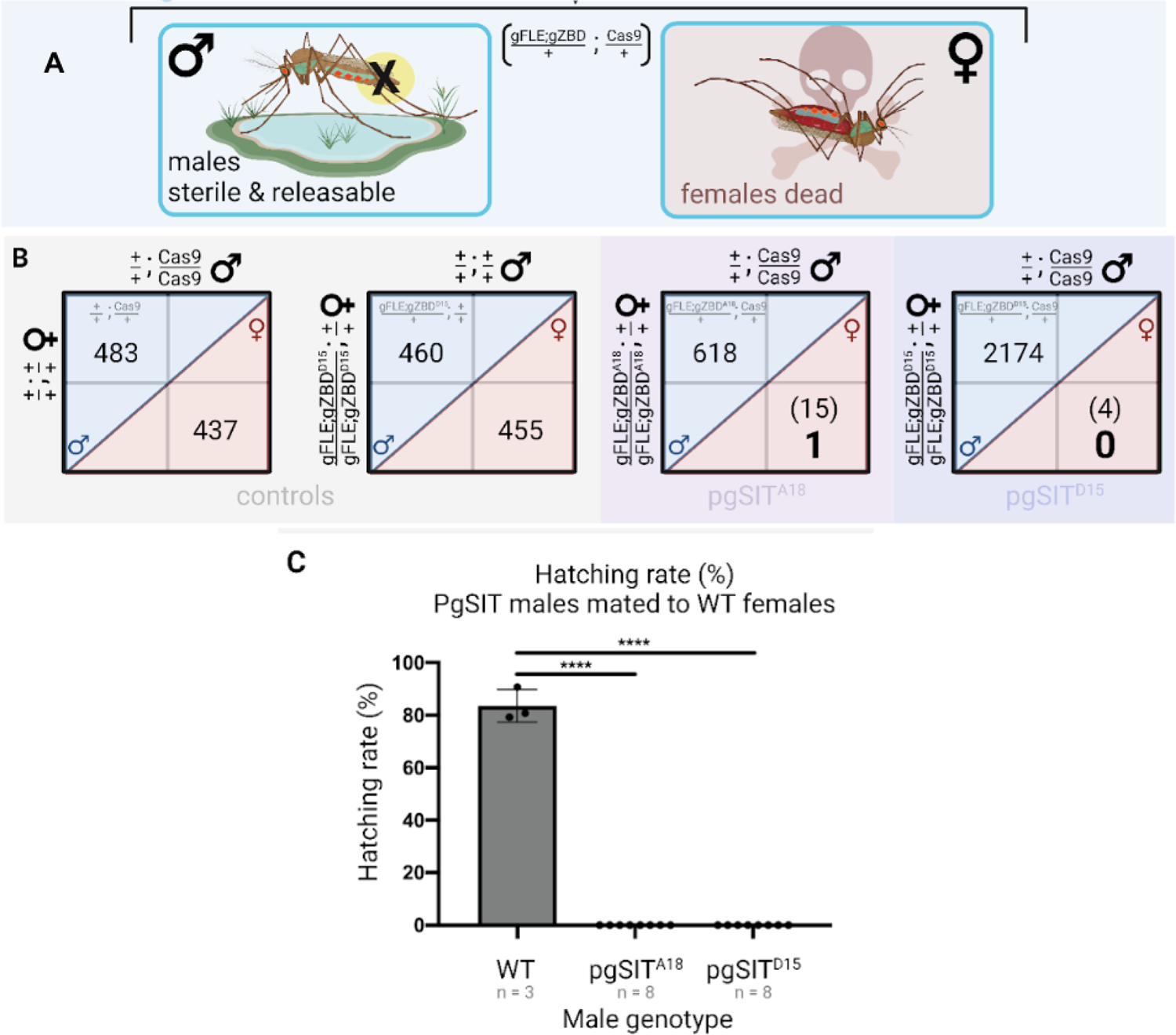
pgSIT production of sterile *Anopheles gambiae* males. **A)** Overview of expected phenotypes from gFLE;gZBD / Cas9 transheterozygotes. gFLE targets a female essential gene, *fle*, and gZBD targets the *zero population growth (zpg)*, *beta-2 tubulin (*β*2-tubulin),* and the female-specific transcript of the *doublesex* (*dsx*) genes. **B)** Testing two gFLE;gZBD families reveals complete female killing when crossed to Cas9 compared to controls. **C)** When mated to wild type females, pgSIT males sire no progeny (0% hatching rate, number larvae hatched/total number of eggs). Modified from (Smidler, Apte, et al. 2023).

### Pre-Large Scale URR Release Risk Management

This early stage benefit cost analysis is prudent for planning and risk management of the pgSIT technology. Much of the early stage funding will be sought from philanthropic organizations with a need for milestone-driven deliverables to minimize risk and optimize their funding portfolios. At this stage, we suggest the following pre-release milestones and “go/no go” criteria to provide the data needed for regulatory approvals, to evaluate the efficacy, scalability, risk and cost of the pgSIT technology, and to develop the facilities, and infrastructure needed to support large scale release of the technology. This milestone-driven approach will provide a stepwise structure to evaluate and de-risk the project.

### Milestone 1: Improved mass production efficiency estimates for pgSIT technology

The cost of pgSIT mass production is dependent on the efficiency of mass producing pgSIT mosquitoes. These mass production processes are detailed throughout this report, but their scale and costs rely primarily on the efficiencies of mass rearing and sex sorting. The International Atomic Energy Agency (IAEA) has produced guidelines for mass rearing Anopheles mosquitoes that was used as a basis for some of the rearing practices and survival and fecundity efficiency estimates that were key to this analysis. Based on this guidance, we used a 50 - 75% survival estimate for first instar/ first larval stage (L1) larvae to adulthood. If this estimate is significantly below 50%, additional resources will be required to optimize survival, or the benefit costs will need to be re-evaluated for pgSIT. The rearing process is inexpensive, however, so production to compensate for lower survival rates is likely not cost-prohibitive. We also used the literature derived fecundity (average number of eggs laid per female over their lifespan) estimate of 210 or 300 eggs (A. S. Yaro et al. 2006; Agyapong et al. 2014). If the average female fecundity cannot be optimized within this range, the benefit cost and cost effectiveness of this intervention should be revisited. Again, mosquito production is affordable, so there should be minimal added cost for production increases to compensate for lower fecundity.

Sex sorting technologies are a larger expense for a pgSIT facility and their efficiency and the mass rearing efficiency directly impacts the sorting throughput and requisite production output. There are multiple competitors in this field, but these technologies will have to be evaluated as mass sex sorting *Anopheles* gambiae mosquitoes is a novel use. Union Biometrica, has been transparent with pricing and expected capabilities so we have good estimations for the efficiency of their equipment. The pgSIT technology is also designed to support a high-throughput fluorescence marker based sex sorting procedure, which is supported by the COPAS FP 500. Using our mass rearing efficiency estimates and the expected sorting capability of a COPAS FP 500, only 15 - 31% capacity of a single device is needed to to support daily pgSIT releases in the Upper River Region. These estimates may change if mass rearing or COPAS FP 500 sex sorting efficiencies are lower than expected. This cost benefit analysis includes costs to address any reduced sorting capacity, such as a spare device and enrollment in a service program, but this technology has not been evaluated in long-term, large-scale use studies. Studies to evaluate the COPAS FP 500 capabilities are underway and will help determine if this technology can support mass production of pgSIT mosquitoes. These studies will allow us to more accurately estimate the cost of scaling pgSIT for mass field release. Alternative sorting technologies can also be explored, including technologies from Senecio Robotics (Tel Aviv, Israel), Verily Life Sciences (South San Francisco, CA, USA) and others.

### Milestone 2: Improved estimates of pgSIT efficacy, scalability and safety in preparation for larger field release

These trials consist of a series of efficacy, scalability, and safety studies that are confined geographically or in cages (**Fig. 4**). These experiments provide more accurate estimates of field performance and behavior as they are conducted under conditions similar to those expected in the field. Updated estimates from these confined trials could improve cost benefit and release rate estimates for future field trials and large scale release programs.

### Milestone 3: Improved estimates of pgSIT efficacy, scalability and safety from small scale field studies to prepare for large scale release in malaria public health programs

A series of field trials can provide the entomological mosquito suppression and epidemiological malaria reduction data needed to better understand how pgSIT will perform for mosquito and malaria control on a larger scale. PgSIT is expected to reduce and potentially eliminate *Anopheles gambiae* populations locally and consequently reduce malaria cases in the treated region. The design and scale of these field trials will be informed by data collected in the confined trials and in early field trials.

## Results

### Objective 1: Modeling release scenarios of pgSIT in The Gambia

We used the MGDrivE 3 framework (Mondal, C, and Marshall 2023) to simulate releases of pgSIT *A. gambiae* mosquitoes to control malaria in the Upper River region of The Gambia. MGDrivE 3 is a modular framework for simulating releases of genetic control systems in mosquito populations and includes modules for: i) inheritance (i.e., the dynamics of the pgSIT system), ii) life history (i.e., the development of mosquitoes from egg to larva to pupa to adult), and iii) epidemiology (i.e., the reciprocal transmission of malaria parasites between mosquitoes and humans) (**Fig. 2A-D**). To simulate epidemiological outcomes of relevance to the cost-effectiveness analysis, we linked the MGDrivE 3 framework (Mondal, C, and Marshall 2023) to the Imperial College London (ICL) malaria model (Griffin et al. 2016, 2010). The ICL malaria model provides a validated and parsimonious framework to describe malaria transmission in human populations, including important features such as acquired and maternal immunity, symptomatic and asymptomatic infection, superinfection, age structure, biting heterogeneity, and antimalarial drug therapy and prophylaxis (**Fig. 2E**). Full details of the model framework are described in the Methods section.

**Fig. 2.**
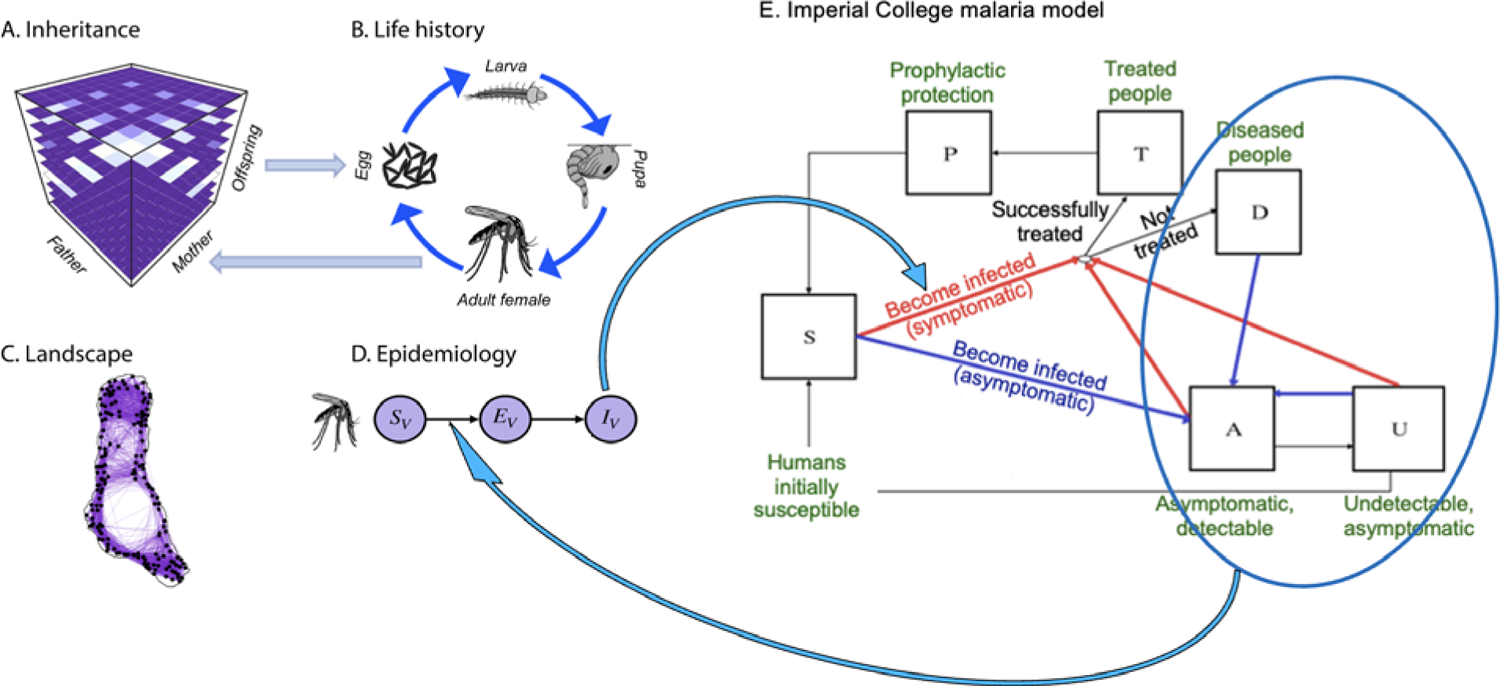
Modules of the MGDrivE 3 framework linked to the ICL malaria model. (A) Genetic inheritance is embodied by a three-dimensional tensor referred to as an “inheritance cube.” Maternal and paternal genotypes are depicted on the x and y-axes and offspring genotypes on the z-axis. (B) Mosquito life history is modeled according to an egg-larva-pupa-adult (female and male) life cycle in which density dependence occurs at the larval stage, and life cycle parameters may vary as a function of environmental variables over time. Genotypes are tracked across all life stages, and females obtain a composite genotype upon mating - their own and that of the male they mate with. Egg genotypes are determined by the inheritance cube. (C) The landscape represents a metapopulation in which mosquitoes are distributed across population nodes and move between them according to a dispersal kernel. (D) The epidemiology module describes reciprocal transmission of a vector-borne pathogen between mosquitoes and humans. Transmission in the human population is modeled according to the ICL malaria model, in which humans move between the following infectious states: S, susceptible; T, treated clinical disease; D, untreated clinical disease; P, prophylaxis; A, asymptomatic patent infection; and U, asymptomatic sub-patent infection.

The modeling framework was calibrated to malaria prevalence data from The Gambia (Dabira et al. 2022), and informed by local entomological data (Soumare et al. 2022) and rainfall data sourced from Climate Hazards Group InfraRed Precipitation with Station data (CHIRPS, https://www.chc.ucsb.edu/data/chirps). The model was also calibrated with existing coverage of artemisinin-based combination treatment (ACTs), LLINS, and IRS with insecticides specific to the Upper River region based on data from the Malaria Atlas Project (Pfeffer et al. 2018). The seasonal carrying capacity of the environment for *A. gambiae* larvae was determined from the CHIRPS rainfall data, entomological data suggesting that dry season larval carrying capacity was ∼10% that of the peak rainy season (Soumare et al. 2022), and for consistency with human malaria prevalence data from a recent randomized controlled trial conducted in the Upper River region (Dabira et al. 2022). Further bionomic parameters for the *A. gambiae* population were sourced from the literature and are summarized in **Table S1**. Finally, parameters describing the pgSIT system were based on laboratory data thus far generated in *D. melanogaster* (Kandul et al. 2018) and *Ae. aegypti* (M. Li et al. 2021), which suggests the pgSIT system in *A. gambiae* can induce complete male sterility and female inviability, with inviability being manifest at pupation (e.g., through an adult flightless phenotype). Based on laboratory data (Smidler, Apte, et al. 2023), we assumed a 50% reduction in pgSIT male mating competitiveness. To be conservative, we also assumed a 25% reduction in pgSIT male lifespan compared to wild-type males, despite no reductions in lifespan being observed in this work (M. Li et al. 2021; Kandul et al. 2018).

With the parameterized modeling framework in place, weekly releases of pgSIT *A. gambiae* eggs were simulated from the beginning of the rainy season (June 1st), for a variable number of weeks and release sizes. Previous pgSIT modeling studies (M. Li et al. 2021; Kandul et al. 2018) suggested that local mosquito populations could potentially be eliminated by ∼10-24 consecutive releases of ∼40-400 pgSIT eggs per wild adult. We explored release schemes within and slightly below this range, focusing on schemes that required smaller weekly release sizes, as the following cost analyses determined this to be the most cost-efficient approach. This exploratory analysis predicted two release schemes with relatively low release sizes would be effective at eliminating the local *A. gambiae* population - ≥12 weekly releases of 32 eggs per adult mosquito, or ≥10 weekly releases of 40 eggs per adult mosquito. For the purpose of the cost-effectiveness analysis, we opted for 12 weekly releases of 32 eggs per adult mosquito (**Fig. 3**). In addition to eliminating the local *A. gambiae* population, in the second year following the beginning of releases (June 1st - May 31st), this led to 10,673 fewer total cases (**Table 1**) and 184 fewer deaths (**Table 2**). This release will be done near the end of the year at the beginning of the rainy season so the case and death rates are not fully reflected until the following year and the year after that, at approximately 13,000 cases prevented and 230 deaths prevented across the Upper River Region. Age-stratified (0-5, 5-17, 17-40, 50-60, and 60 years) cases and deaths averted are shown in **Tables 1-2**.

**Fig. 3.**
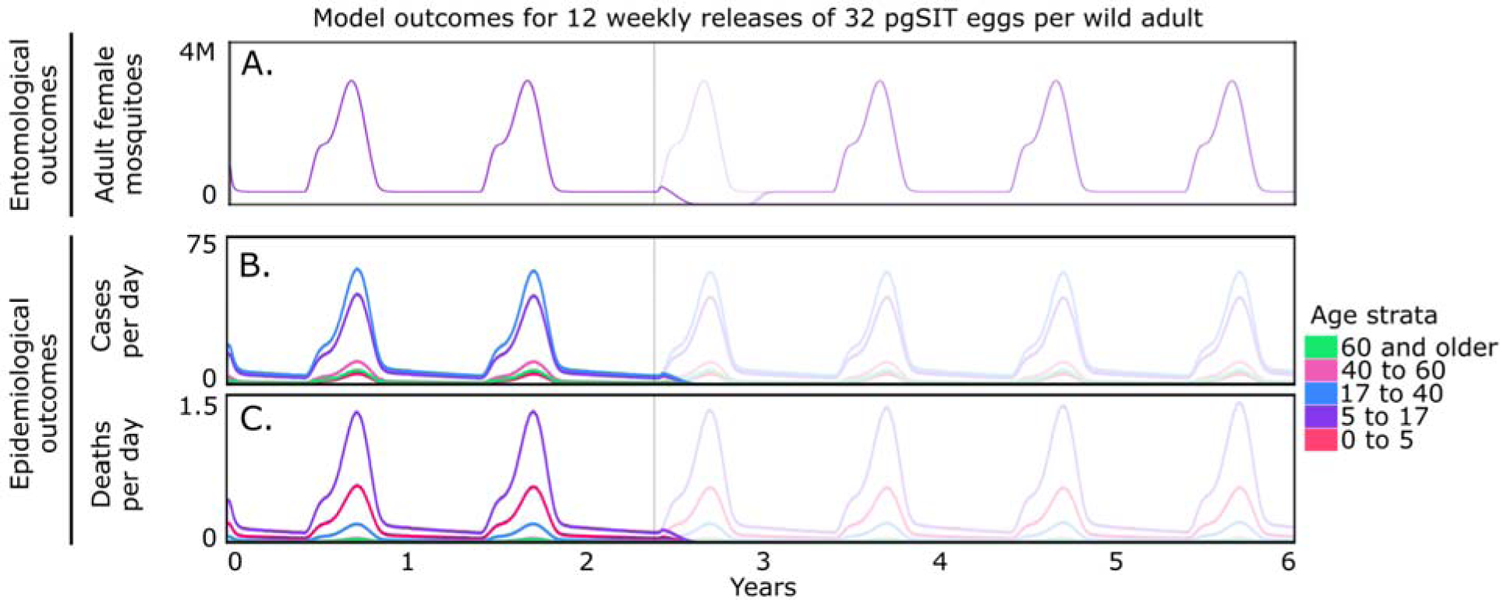
Predicted reduction in entomological and epidemiological outcomes following a cost-efficient pgSIT release campaign.

**Table 1.**
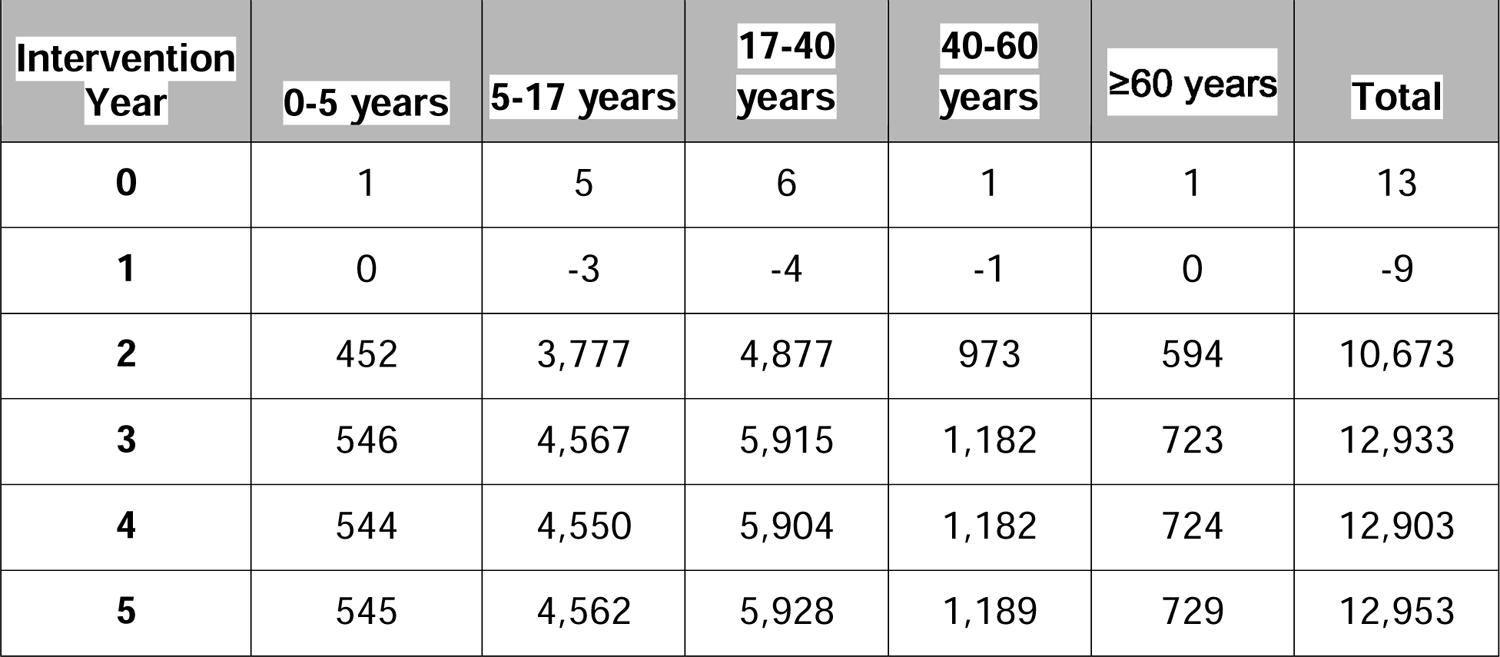
Model-predicted annual malaria cases averted for 12 releases of 32 pgSIT eggs per wild adult.

**Table 2.** Model-predicted annual deaths averted for 12 releases of 32 pgSIT eggs per wild adult.

| **Intervention**<br>**Year** | **0-5 years** | **5-17 years** | **17-40**<br>**years** | **40-60**<br>**years** | **≥60 years** | **Total** |
| --- | --- | --- | --- | --- | --- | --- |
| **0** | 0 | 0 | 0 | 0 | 0 | 0 |
| **1** | 0 | 0 | 0 | 0 | 0 | 0 |
| **2** | 48 | 114 | 16 | 3 | 2 | 184 |
| **3** | 57 | 141 | 20 | 4 | 3 | 225 |
| **4** | 57 | 143 | 21 | 4 | 3 | 228 |
| **5** | 57 | 146 | 21 | 4 | 3 | 232 |

An achievable 12 consecutive weekly releases of 32 pgSIT eggs per wild adult were simulated in a mosquito population resembling that of the Upper River region of The Gambia. Releases begin on June 1st of the second year and are denoted by a vertical line in each figure panel. Of the release schemes we explored, this was determined to be the most cost-efficient. Full model details are provided in the Methods section, and model parameters are provided in **Table S1**. Model outcomes depicted include: i) adult female mosquito density (purple), ii) daily malaria cases averted (colors represent age strata, depicted in key), and iii) daily deaths averted (age strata depicted in key). Solid lines represent predicted pre-intervention dynamics, and low-opacity lines represent dynamics during and post-intervention.

In summary, mathematical modeling predicts that weekly releases of 32 pgSIT eggs per wild adult would **result in the elimination of the local *A. gambiae* population** in the Upper River region of The Gambia when carried out over 12 consecutive weeks, and with it, locally acquired cases of malaria and deaths resulting from severe cases. This is predicted to reach 13,000 fewer malaria cases and 230 fewer deaths annually post intervention year 1, and comparable reductions in subsequent years. The highest number of averted cases are in the 5-17 and 17-40 year age groups, and the highest number of averted deaths are in the 0-5 and 5-17 year age groups. As with any modeling analysis, there are limitations. Importantly, the spatial structure has been ignored in this analysis, and the URR has been treated as a closed population without migration. The results of this analysis will apply most closely to the first year following the intervention, as mosquitoes from outside the URR are likely to recolonize between seasons, and would require subsequent pgSIT releases for maintenance of suppression, albeit at likely lower levels. The modeling study is also an overly simplified representation of mosquito and malaria dynamics in the URR, and subsequent modeling efforts would be strengthened by data generated from pgSIT and other mosquito suppression interventions.

### Objective 2: Assessment of the costs and health benefits of eliminating malaria from The Gambia

#### Task 2.1 Cost estimate of implementing pgSIT in The Gambia

##### 2.1.1a Pre-deployment technology development

The pgSIT technology has been validated in the laboratory for the small scale production of sterile males (**Fig. 1**). However, prior to wider scale use as part of a regional or national vector and malaria control program, additional technological development and field trials are necessary to improve pgSIT scalability and confirm its safety and performance in the field (Smidler, Apte, et al. 2023).

###### Development of pgSIT sex separator line (pgSIT 2.0)

Sex sorting technologies, which are needed to sort the Cas9 and gRNA parent lines prior to the generation of sterile males, is a limiting factor for the scaling of pgSIT technologies. Few scalable technologies exist, but their sorting strategy can greatly impact their production capabilities and costs (discussed in **Section 2.1.2.4**). One of the most cost-effective and scalable sex sorting technologies, the Complex Object Parametric Analyzer and Sorter (COPAS) FP 500 (Union Biometrica), requires the pgSIT Cas9 and gRNA lines to be engineered with sex-specific fluorescent markers. Fortunately, we have just developed a robust sex-sorting approach termed SEPARATOR (<u>Sexing Element Produced by Alternative RNA-splicing of A Transgenic</u> <u>Observable Reporter</u>) that exploits sex-specific alternative splicing of an innocuous reporter to ensure exclusive dominant male-specific expression (Weng et al. 2023). To use SEPARATOR to scale pgSIT production, we will, therefore, need to redesign our pgSIT lines to incorporate SEPARATOR, termed pgSIT 2.0, and evaluate these lines for sterile male production.

##### Field trials

Prior to large-scale use of the pgSIT technology, confined field trials, and open field trials are required to generate the safety and efficacy data needed to justify the scale up of the intervention (discussed in **Section 2.1.3.20**). This stepwise approach, outlined by the World Health Organization (WHO) (**Fig. 4**), begins with Phase I laboratory safety and efficacy studies, which we have completed for the pgSIT technology (**Fig. 1**). It then focuses on Phase II confinable studies, in which mosquitoes are tested in the local environment, against local mosquitoes, and often after introgression into the genetic background of the local wild population, but a cage, or another barrier prevents their full release into the environment. These confined trials limit the risk to the environment while providing important safety and efficacy data prior to open field release, but lack the necessary rigor needed to fully assess the safety and performance of the technology. If successful in Phase II, a series of increasingly larger and more complex Phase III open field releases are then necessary to first provide safety and assessments of entomological parameters before focusing on the epidemiological impact on malaria transmission. The Phase III open releases will then be used to determine whether pgSIT can be used for large scale vector and disease control (Phase IV).

**Fig. 4.**
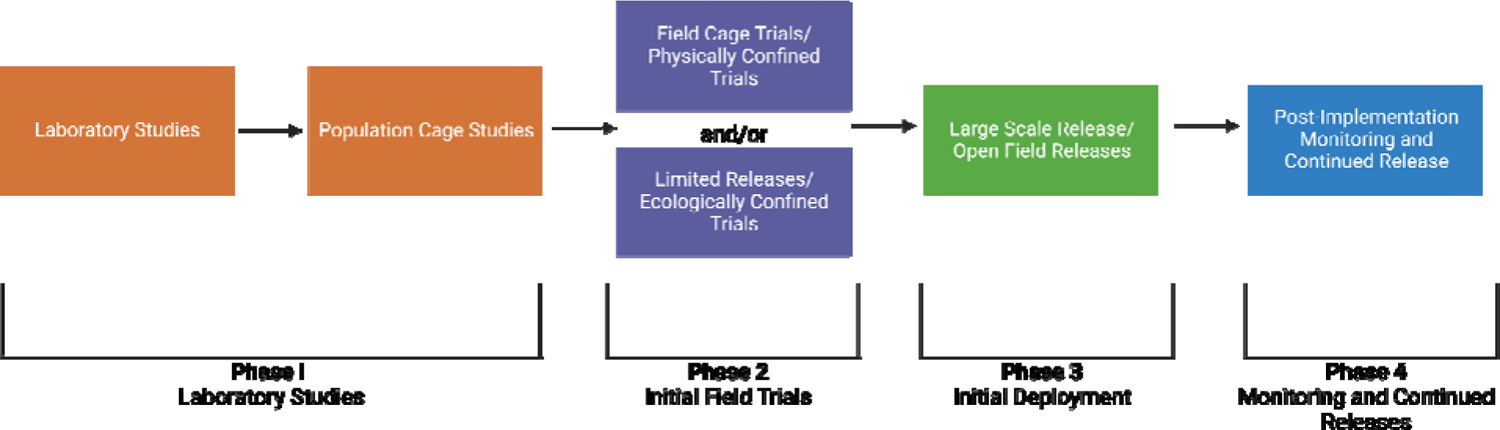
Phased testing pathway for genetically modified mosquitoes.

This figure was based on guidelines and a figure by WHO (Health Organization n.d.). Figure generated in BioRender.com.

##### 2.1.1b Technological development, infrastructure and training for regional deployment of pgSIT sterile males

###### Scaling technologies

*Anopheles* mosquito production at the scale required for large field releases is complex, but has been achieved for SIT programs, so there are technologies to scale *Anopheles* production (Fabrizio Balestrino et al. 2011; F. Balestrino, Benedict, and Gilles 2012; FAO/IAEA 2017). We plan to optimize and validate these existing technologies and some new ones, to support pgSIT sterile male production for population suppression. pgSIT has similar technical requirements for mass production as traditional SIT. Both approaches require mass rearing systems and techniques, and an efficient sex separation method, but pgSIT does not require the labor intensive steps of irradiation-based sterilization, or sex separation of the adults just prior to field release, which simplifies the scaling of pgSIT compared to traditional SIT (**Fig. 5**). Until we are able to test these technologies, however, there are uncertainties that need to be built into our production and costs estimates for scaling pgSIT (discussed in Sections **2.1.2.4 - 2.1.3.21**).

**Fig. 5.**
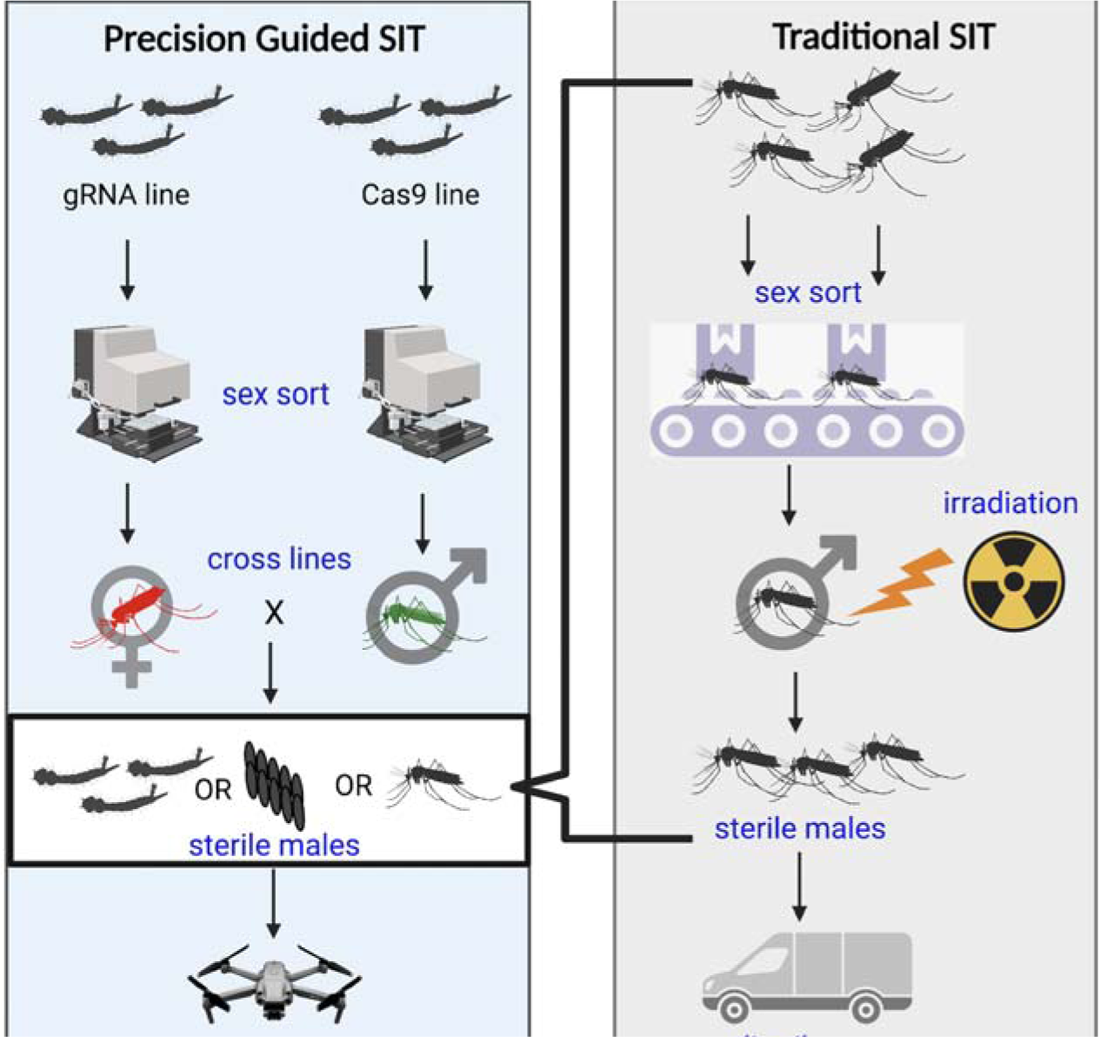
Overview of the production differences between pgSIT and traditional SIT.

When pgSIT is used to produce sterile males (left-blue box) only the parent Cas9 and gRNA lines need to be sex sorted prior to crossing to generate sterile males. No sorting or treatment of the final product (white box) is required whatsoever, so sterile males can rapidly be produced and distributed into the field. With traditional SIT (right-gray box), however, the final product needs to be sex sorted to isolate the males, the males need to then be irradiated to sterilize the males before they can be distributed to the field. The traditional SIT approach, therefore, has increased labor and costs associated with the extra processing of the sterile males and to compensate for fitness reductions, or mortality associated with the increased handling and irradiation of fragile males. The additional processing time and reduced fitness also limit the distance in which the traditional SIT males can be distributed. Figure generated in BioRender.com.

### Deployment technologies

Since pgSIT involves minimal handling of the sterile male product, the eggs can quickly be packaged and delivered to their field site (**Fig. 5 and 6**). We will focus on drone egg delivery to the field site, which can minimize the impacts of poor transportation infrastructure on egg delivery, making the technology accessible to all areas of need. However, testing and optimization of the current drone technologies will be necessary to ensure they can meet the delivery requirements of the project (discussed in Section **2.1.3.11**).

**Fig. 6:**
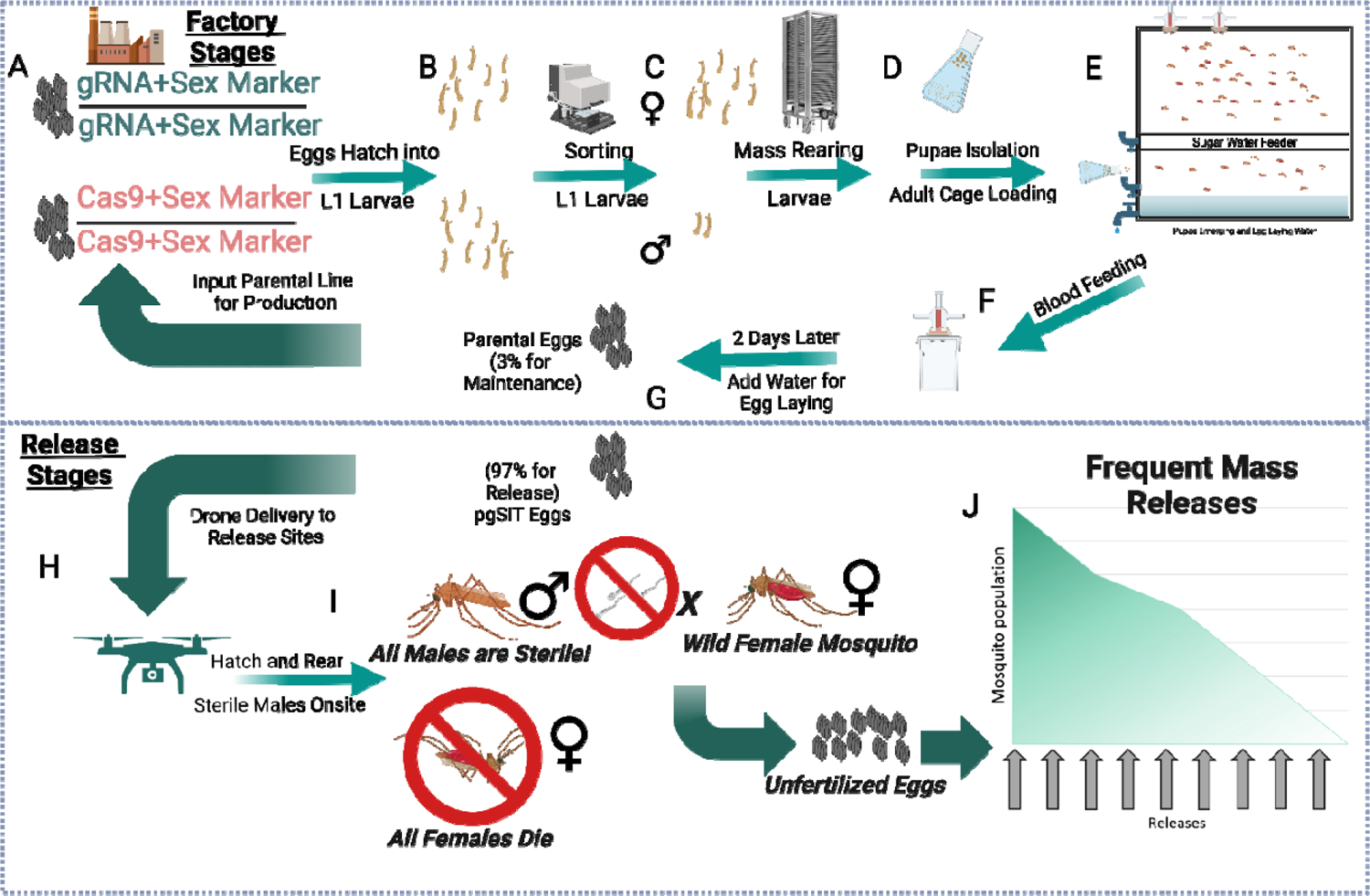
Mass rearing during the facility’s active phase.

### Facility design

A pgSIT production facility needs to support the equipment and activities required to generate sterile males at the scale needed for release. Precise estimates of the size and cost of the facility are reliant on the results of early field studies of the sex sorting capabilities, mosquito survival under mass rearing conditions, and the fecundity, or the number of eggs laid per female of the parent Cas9 and gRNA lines. However, using estimates from traditional SIT scaling programs, we provide a range of estimates for the design size and cost of these facilities (discussed in **Section 2.1.3.21**). Options to include easily expandable facilities or modular additions in the facility design may allow us to account for these uncertainties during the planning and scale-up of pgSIT production.

### Training and personnel

The goal is to generate a community-driven program led and executed by community leaders and operations and technical experts in The Gambia. However, the skills needed to plan and accomplish the scaling of pgSIT technology are specialized, so an initial one year training period will provide time for local staff to obtain the expertise needed for high-throughput sterile male production. Prior to this training, pre-deployment field trials will provide opportunities to exchange knowledge with the community and identify and train future leaders and managers of the facility. Upon completion of this training, operations will be locally managed in The Gambia. We provide multiple wage estimates for the different roles anticipated for the facility to generate a range of training and personnel costs (discussed in **Sections 2.1.3.18 - 2.1.3.20f**).

### Operations and distribution

Sterile male releases will be done for 12 weeks per year to enable complete suppression and predicted localized extinction. In order to maximize production efficiency, we anticipate three phases of the facility production cycle: Maintenance Phase, Ramping Phase, and Active Phase (**Section 2.1.4**). The Maintenance Phase will maintain a small population of the gRNA and Cas9 lines on three week cycles with minimal effort and resources (**Section 2.1.4.1**). The Ramping Phase increases population numbers to support the mass production of sterile males in the Active Phase (**Section 2.1.4.2**). The Active Phase facilitates the high production of mosquito eggs for 12 weeks for delivery via drone to the Upper River Region (**Section 2.1.4.3**). Drones will make a daily delivery of eggs and each release site will require one batch of eggs per week.

### Onsite rearing in the Upper River region

Eggs delivered to the URR need to be reared to adulthood. Multiple options to rear the mosquitoes are discussed, and their costs are estimated and range from standard rearing conditions in plastic trays (**Sections 2.1.3.13 - 2.1.3.14**) to synthetic ponds with netting to mass rear the sterile male mosquitoes. Evaluation of these methods during the field trials, however, will identify the best method for this intervention. In the meantime, we provide cost estimates for multiple onsite rearing approaches. This cost may be unnecessary as other interventions rely on volunteer labor for distribution. Onsite rearing could similarly rely on local resources as mosquitoes can survive in local water sources, so these could be used to rear the larvae with local materials.

#### 2.1.1c Overview of goals and assumptions

Understanding the process of producing pgSIT insects for releases is crucial for conducting a cost assessment of the technology. The costs of implementing pgSIT are directly related to the steps involved in the process, including the building of a rearing facility, insect production within the facility (rearing, blood feeding, mating, harvesting eggs), and releasing eggs into the target population. Without a thorough understanding of these steps, it is difficult to accurately assess the costs of implementing pgSIT and its cost compared to alternative mosquito control strategies.

This exercise will start with an overview of the pgSIT production process and then delve into the costs associated with each step. **Section 1.2** describes how we calculated the release scenario required to achieve local extinction of *A. gambiae* (32 eggs per wild adult per week for 12 weeks). Therefore, this section is built upon the goal of deploying this many mosquitoes per season. This release scenario is likely an overestimation, however, as there are likely year-to-year suppression effects that will reduce the size of the target population in later treatment years and therefore require a less intensive release scenario to achieve the same level of suppression in later treatment years. As rearing experience, mechanization, and sexing efficiency are increased in the facility over time, we expect improved production efficiency. Additional cost benefits to the project expected over time come from improvements in the supply chain and optimizing delivery and release methods. More than likely, as production processes and the delivery pipelines improve, this facility will have reduced production and delivery requirements to the target regions, and resources can be reallocated towards suppression in other regions or turned towards export. However, the actual impact and timeline of these gains are uncertain, so the facility is designed to fully suppress *A. gambiae* in the URR with the same release scenario each year again.

With this in mind, we provide both high and low estimates of the yearly and total costs based on three key production variables: female fecundity, hatch rate, larvae to adult survival, and sex sorting methods. These estimates are aimed at meeting the requisite production level for local extinction, but fecundity, hatch rate, percent survival to adulthood, and sex sorting at this scale have associated uncertainties until they can be further investigated. Rearing density, for example, impacts larval development, larval survival, and adult size (Muriu et al. 2013). To scale egg production, there is a trade-off between optimizing production and density-related increases in development time and mortality, which will reduce production efficiency.

The mosquito fecundity, the number of eggs produced per female in their lifetime, is impacted by their rearing conditions (C. D. Christiansen-Jucht et al. 2015; C. Christiansen-Jucht et al. 2014) and their blood-feeding procedures (Takken, Klowden, and Chambers 1998; Chikwendu, Onekutu, and Ogbonna 2019; de Swart et al. 2023). Mosquitoes should be uniformly blood fed, but uncertainties in optimal blood meal source, feeding frequency, and feeding methods make estimating the costs of this important factor imprecise. Optimal blood-feeding procedures will have to be established for the facility and will also be restricted by the blood meal sources reliably available in the region. To account for this variability, we estimated the costs for (1) high fecundity (300 eggs per female) or (2) moderate fecundity (210 eggs per female) production (A. S. Yaro et al. 2006; Agyapong et al. 2014). We expect that we will be able to get three oviposition cycles during the lifetime of a mosquito. The literature varies on the expected fecundity of *A. gambiae* over sequential oviposition cycles but suggests that 300 eggs per female are a conservative estimate under laboratory conditions, since up to 1000 eggs per female have been recorded (targetmalaria.org 2020; Clements 1992). However, if mass rearing unduly complicates blood feeding and oviposition, we can expect a concurrent decrease in fecundity. We estimated a 30% fecundity decrease (210 eggs per female) to address these possible complications. Fecundity will have a large impact on the production output of the facility (**Table 3**), potentially increasing production and upstart costs, augmenting equipment needs (e.g., the number of COPAS machines, rearing racks, and adult cages), and increasing production and maintenance costs overall. More equipment will also impact the size and cost of the facility, but including easily expandable facilities or modular additions in the facility design may reduce these costs.

**Table 3:**
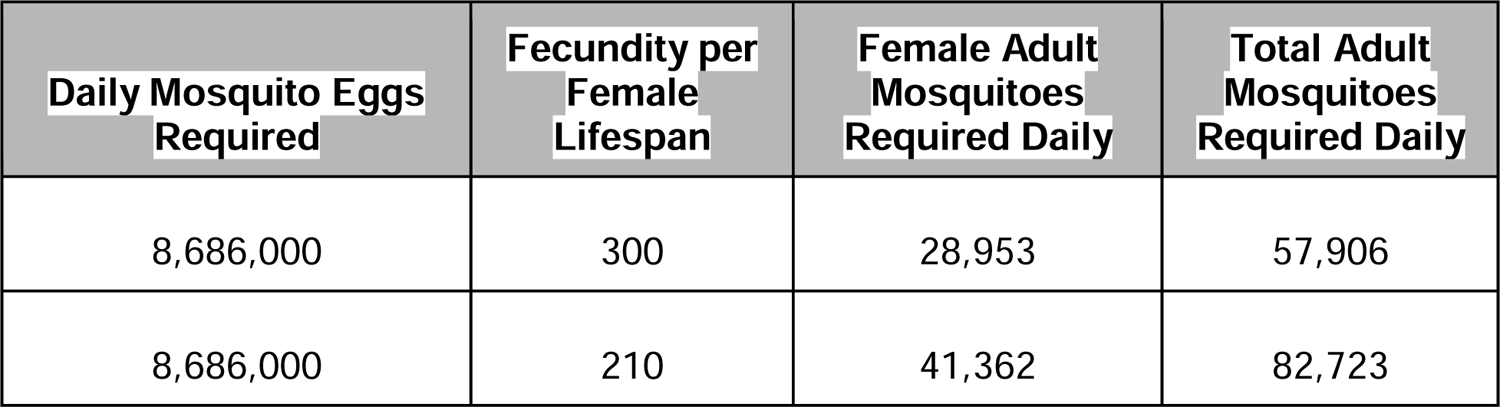
Total adult mosquitoes required for egg production.

| Daily Mosquito Eggs Required | Fecundity per Female Lifespan | Female Adult Mosquitoes Required Daily | Total Adult Mosquitoes Required Daily |
| --- | --- | --- | --- |
| 8,686,000 | 300 | 28,953 | 57,906 |
| 8,686,000 | 210 | 41,362 | 82,723 |

The second variable that has a large impact on cost is the larvae to adulthood survival. Not all *A. gambiae* larvae will reach adulthood, especially at high density production levels. The IAEA protocol for the mass rearing of *Anopheles* mosquitoes for SIT programs estimates a 50-75% expected larval to adult survival rate in their mass-rearing trays (FAO/IAEA 2017). To achieve the required number of adults, 33.3% - 50% extra larvae are needed to compensate for larval stage mortality. This increased larval production results in 33.3%-50% increases in equipment (rack and tray systems), equipment run times (COPAS sex sorting), and supplies to support larval development (food and water). There are also labor costs associated with the increased production. By building these considerations into the facility design and procedures, we can optimize the density and larval survival trade-offs to optimize adult production. The results demonstrated in the IAEA protocol are likely a conservative range of larval survival expectations in our facility, so we will use this to estimate larval mortality in our facility (**Tables 4 and 5**).

**Table 4:** COPAS FP 500 larvae daily rearing requirements.

| Conditions | Fecundity | Survival | Mosquito Larvae Required for Production | Total Mosquito Larvae Required |
| --- | --- | --- | --- | --- |
| **High Fecundity,**<br>**High Survival** | 300 | 75% | 77,208 | 81,840 |
| **Low Fecundity,**<br>**High Survival** | 210 | 75% | 110,297 | 116,915 |
| **High Fecundity,**<br>**Low Survival** | 300 | 50% | 115,812 | 122,761 |
| **Low Fecundity,**<br>**Low Survival** | 210 | 50% | 165,446 | 175,394 |

**Table 5:** Cost of COPAS FP 500 and annual service fee.

| **Conditions** | **Daily Number of Larvae Sorted** | **Number of COPAS Machines Needed** | **Total COPAS Needed (with spare)** | **Equipment Cost (Machines + Laser Spares) USD** | **Annual Service Fee\* USD** | **Total Upfront Cost# USD** |
| --- | --- | --- | --- | --- | --- | --- |
| **High Fecundity, High Survival** | 81,840 | 0.15 | 2 | 759,072 | 91,089 | 948,840 |
| **Low Fecundity, High Survival** | 116,915 | 0.21 | 2 | 759,072 | 91,089 | 948,840 |
| **High Fecundity, Low Survival** | 122,761 | 0.22 | 2 | 759,072 | 91,089 | 948,840 |
| **Low Fecundity, Low Survival** | 175,394 | 0.31 | 2 | 759,072 | 91,089 | 948,840 |

\* Annual maintenance/service fee is 12% of initial machine costs. Additional spare laser costs are excluded from the annual maintenance/service fee calculations.

A rate-limiting step of scaling is the sex sorting required for the parent strains (Cas9 and gRNA lines), from which females of one genotype and males from another are needed. Crossing these strains together generates the releasable pgSIT sterile male eggs. This approach saves significant time, labor, and cost compared to other SIT and female killing technologies since pgSIT does not require sex sorting of the final product (sterile males). Sorting the parental generation amplifies productivity because sorting the equivalent of one mosquito yields approximately 37.5 releasable males (1:37.5 sort: release ratio). To be conservative, we assume these crosses are not reciprocal, which means that the male and female of this cross must be particular genotypes to successfully produce sterile males. However, we will aim to engineer the novel Cas9 and gRNA strains that incorporate the sex-sorting genetic cassette, and these novel strains may enable bi-directional (reciprocal) genetic cross for pgSIT egg production. In the current iteration of pgSIT, to achieve 100% sterile male progeny the fathers must be from the Cas9 line, while the mothers carry the gRNAs genes. However, when applying updated capabilities to pgSIT by adding sex sorting cassettes, there will be the opportunity to update the expression cassettes for Cas9 and gRNAs to be functional bidirectionally. If pgSIT has bidirectional capabilities, this would halve the sex sorting requirement cost and reduce the annual and upfront costs by roughly half. However, this approach may not be technically successful, so this cost assessment assumes the current uni-directional nature of the crosses. The method of sex sorting will impact production and costs, nonetheless, so we evaluate multiple options for this system. The technologies for high-throughput sex sorting commercially available are Union Biometrica’s COPAS FP 500 (**Table 4**) and Senecio Robotics. These methods differ in their rearing requirements as these methods sex sort at different life stages. The COPAS FP 500 sorts larvae shortly after hatching (L1 stage). Senecio Robotics technologies sort mosquitoes as newly emerged adults.

Quality control procedures in the production facility and field monitoring will identify production and field issues. Deactivation of Cas9 expression, or mutations in multiple gRNA targets would cause the pgSIT system to fail and could occur from spontaneous base change or background mutation. The background mutation rate for related *Anopheles* mosquitoes is extremely low minimizing the likelihood of multiple spontaneous mutations inactivating these components. Negative selection is not expected for gRNA components as these constructs have low fitness costs. While there may be some fitness costs for the Cas9 construct, the fitness cost would be calculated between the functional Cas9 line and a mutant line, which the mutant line may still have significant fitness cost as it is producing a large foreign protein. Loss of function may have minimal positive selection for these single base changes compared to the unmutated Cas9. The base change rate is low for related *Anopheles* mosquitoes lending further evidence to the long-term stability of these lines (Rashid et al. 2022). Future experiments can quantify selection for these mutations and determine the quality control procedures, but for the purposes of this cost assessment, regularly testing pgSIT activity by crossing the Cas9 and gRNA lines and monitoring for unexpected non-sterile males or females is a functional and inexpensive test that can easily implemented at key points in the production process.

Due to these uncertainties, our cost estimates will be based on both efficient fecundity (300 eggs per female) and less efficient fecundity (210 eggs per female) estimates. There will also be four estimates associated with the larvae food, water consumption, rack and tray system, and sex sorting systems to address the variation in larval survival. These costs will be summarized with minimum and maximum cost estimates.

The costs will be broken into four sections: **2.1.2** Mosquito production requirements, **2.1.3** Description of the mass rearing techniques and costs of equipment and facility development, **2.1.4** Mosquito rearing facility phases and labor costs, and **2.1.5** Development costs of testing and field trials.

#### 2.1.2 Mosquito production requirements

##### 2.1.2.1 Egg production required for The Upper River region

Estimates of the total egg production are needed to determine the facilities required to meet this demand. The mathematical modeling estimates a release scenario of 32 eggs per adult mosquito per week, but this number is dependent on the total number of mosquitoes in The URR. With a population of approximately 265,000 people (Gambia Bureau of Statistics 2015) and an estimate of ∼8 mosquitoes per person on average (∼2.6 per person in the dry season and ∼25 per person at the peak of the rainy season), we estimated the number of adult mosquitoes required per week. The estimates of mosquitoes per person are calculated based on the malaria prevalence in the population in 2017 and the number of mosquitoes that would need to be present to support this rate of malaria. This indirect estimate is based on the malaria rates and population size, but this is a useful approximation as the URR lacks extensive direct entomological studies (Dabira et al. 2022). These estimates give us approximately 1.9 adult mosquitoes to target for suppression. These numbers are multiplied to give us a weekly requirement of just over 60.8 million eggs. Producing eggs simultaneously is not the most efficient use of the equipment. It puts unnecessary strain on the staff and production equipment. To address this, the production of these 60.8 million eggs can be spread across the entire week. This approach gives us a daily egg requirement of just under 8.7 million eggs. The daily egg production can be used to determine the number of adult mosquitoes required to produce the requisite eggs and the infrastructure needed to support this level of mosquito production.

##### 2.1.2.2 Adult production requirements for egg production

Using the daily egg production needs calculated from our release scenario modeling (**Objective 1**) and Section 2.1.2.1, we worked backward to estimate the number of females and total mosquitoes required to achieve these numbers. We accounted for the uncertainties in fecundity (**Table 3**), larval mortality (**Table 4**), and sex selection technologies (**Table 4**) to give us a range of cost estimates. For example, female *A. gambiae* in the wild can lay from 800-1,000 eggs in their lifetime (Clements 1992). This optimal amount is unlikely achievable in mass rearing or even normal laboratory conditions. A conservative estimate of lifetime fecundity per *A. gambiae female* is 300 eggs. This estimate is calculated by multiplying the average number of eggs laid per female during the first oviposition (162 eggs), and the hatch rate of 86% and the average percentage of females that lay eggs per oviposition cycle (85%) (A. S. Yaro et al. 2006; Agyapong et al. 2014; Alpha S. Yaro et al. 2006). This means that at first oviposition, each female produces an average of 118 eggs. As the females age, their fecundity at each oviposition cycle decreases (Agyapong et al. 2014). Using a conservative estimate of a 15% decrease between each egg lay, the second and third oviposition cycles are expected to produce 101 and 86 eggs, respectively. The 15% decrease is a conservative estimate of *A. gambiae* female survival across the two weeks of adulthood, which is usually above 70% survival. An estimated 15% death rate between blood feedings, however, results in the conservative estimate of 60% survival to the end of the two weeks of adulthood. This results in 306 mosquitoes per adult female, which we round down to 300 to be further conservative with our estimate. With careful design and rearing experience, we expect to exceed or at least maintain this fecundity rate. However, we are also including the 30% reduced fecundity rate of 210 eggs per female to provide a worst case scenario estimate. These calculations assume approximately a 65% initial blood feeding rate and a further 35% decline between each feeding. The ideal mating ratio is 1:1 according to the IAEA Protocol (FAO/IAEA 2017), so we double the number of females to meet the daily requirement of adult mosquitoes.

The first column shows the daily mosquito egg production requirements that will be needed for releases in the URR. The daily calculations are derived from the 60.8 million egg per week estimate divided daily across the week. The second column accounts for variations in fecundity in mass rearing conditions. We expect to see at least 300 eggs produced per female. If we have a 30% decrease in production, we expect 210 eggs per female. The number of adult females needed to meet the daily requirement is calculated by dividing the daily eggs by fecundity. At a 1:1 male:female mating ratio(FAO/IAEA 2017), we double the adult female number to get the total number of adult mosquitoes.

##### 2.1.2.3 Mosquito survival to adulthood

To produce the number of adult mosquitoes needed for egg production, we also factored in larval survival to adulthood. In laboratory conditions, the larval survival rate can be quite high, but in a mass-rearing system where rearing densities are high, larval survival is lower. While we have not evaluated pgSIT larval survival on a mass scale, the IAEA *A. gambiae* mass rearing guidelines suggest that larval to adult survival in the larval rearing systems range from 50-75% (FAO/IAEA 2017). The upper and lower range of these survival rates were included in our calculations. To maintain the colony, a minimum of 2.2% of the eggs produced must be used to replenish the lines but to be conservative, we estimate that 3% is needed to support the colony. While this would be sufficient to support the colony, we also want to ensure genetic diversity in the stock and therefore we expand this estimate by 6% of the mosquito production. If producing mosquitoes in a higher quantity, 3% could be sufficient to maintain production during the Active Phase. The 2.2% estimate is derived from the least efficient production scenarios (females produce 210 eggs, 50% larvae to adult survival). If each female mosquito produces 210 eggs, each adult (males and females) produces 105 eggs. Including an 86% egg hatch rate and a 50% larva to adult survival rate, results in 45.15 adult mosquitoes produced per adult mosquito in mass rearing conditions(Alpha S. Yaro et al. 2006). To convert this estimate to percentages, we divide 100 by the number of adult mosquitoes produced per adult mosquito to get 2.2%. Using this conservative estimate, and increasing this estimate to 6%, will ensure we meet our production goals and genetic diversity. These variables result in four different potential daily larvae production rates (**Table 4**).

The larvae rearing numbers are based on the assumption that sex sorting is done on newly emerged L1 larvae.

##### 2.1.2.4 Sorting technology impact on mosquito production

The sex sorting technology, used to separate the Cas9 and gRNA pgSIT lines prior to the crossing of the Cas9 males and gRNA females to generate sterile males, can also impact mosquito production. We assessed two commercially available sex sorting technologies that differ in their workflows due to the life stage at which they sort mosquitoes (**Table 4**). COPAS FP (Union Biometrica, Holliston, MA, USA) sex-sorting strategy relies on the large-particle flow cytometry and enables sex-sorting newly emerged 1st instar (L1) larvae using genetically-encoded sex-specific fluorescent markers. The other technology from Senecio Robotics applies optical sorting of mosquitoes as newly emerged adults. This sorting timeline creates a substantial difference in the number of larvae reared to adulthood because COPAS does this initial sorting upfront in the production process, while Senecio Robotics requires double the Cas9 and gRNA larvae production since both sexes of larvae need to be raised to adulthood before they are sorted. Therefore, the Senecio Robotics sex sorting method nearly doubles the larval production effort and labor, in addition to being slower at sorting. Another critical point to note is that Verily Life Sciences has its own proprietary sex sorting technology on a comparable scale, but it is commercially unavailable.

#### 2.1.3 Description of the mass rearing techniques and associated equipment and facility costs

This section describes the necessary facilities, supplies, and equipment for mass rearing mosquitoes and their range of estimated costs.

##### 2.1.3.1 Egg isolation and hatching methodology

Eggs are laid on a moistened surface and then drained from the water post oviposition. The adult mass-rearing cages have a convenient water trough allowing easy access to the water for this process. The eggs are then added to a strain-specific container and left in water overnight (**Fig. 6**). Within two days, larvae are separated from the eggs and sex sorted by COPAS (**Fig. 7**). Alternatively, the mosquitoes are sorted as adults by

**Fig. 7:**
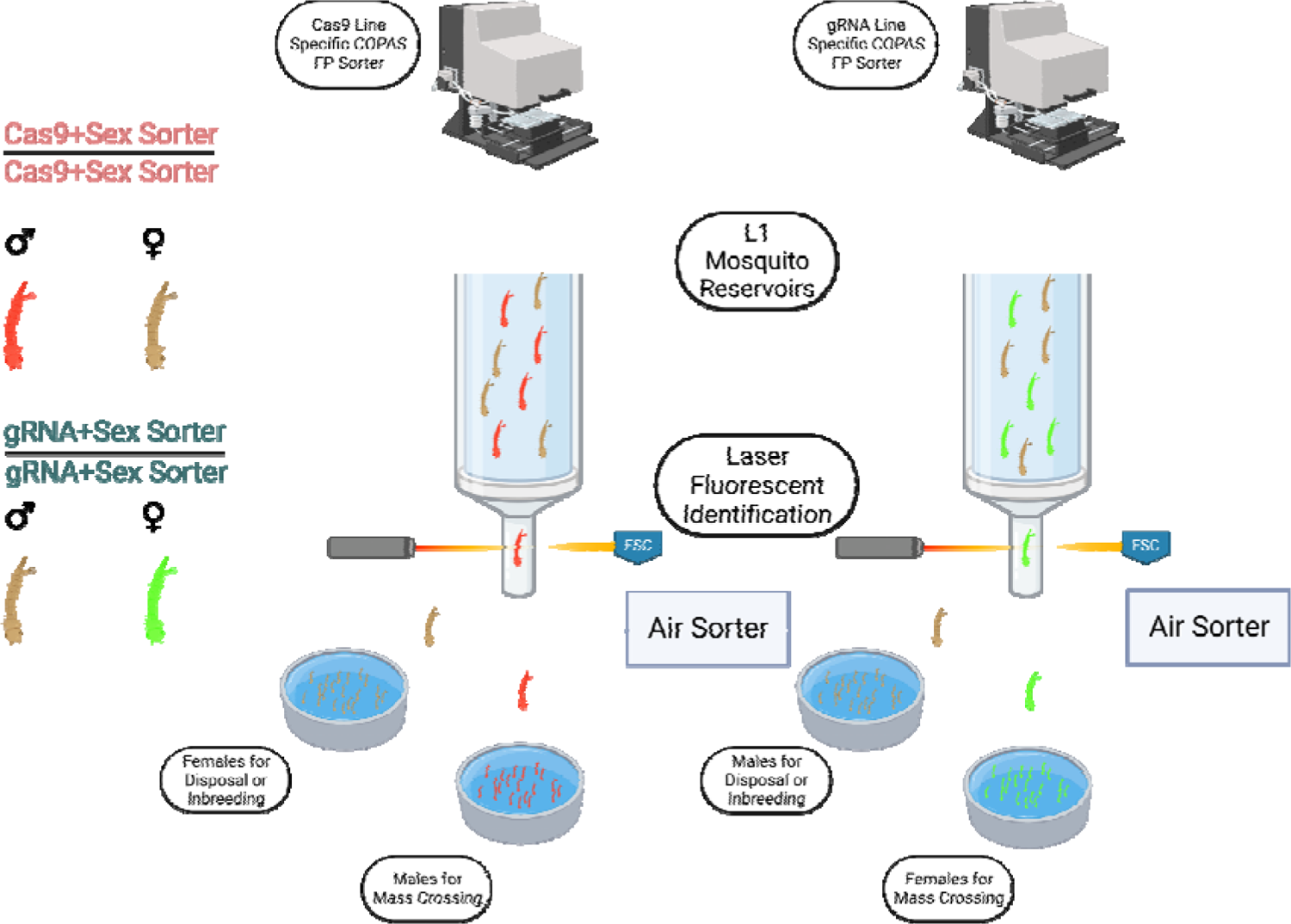
Overview of the COPAS FP-500 larval sorting protocol.

Senecio Robotics sorting machine. Optimization of these procedures will be required to improve efficiency, but there are several options, including a straining system that separates sticky eggs and egg fragment pieces, light aversion techniques to move larvae for retrieval, or swirling or other water isolation techniques. Usage of the COPAS for *A. gambiae* specific applications has been well documented, and robust protocols have been established which can be easily scaled (Marois et al. 2012).

The general process of mass rearing *Anopheles gambiae* mosquitoes when the facility is actively producing mosquitoes for release is shown. Generating pgSIT sterile males (**Factory Stages)**-This begins with hatching the Cas9 and gRNA parent lines(A-B). Assuming that COPAS is used, sex sorting occurs at the L1 nascent larval stage by sex-specific fluorescent markers (C). If Senecio Robotics (or the Verily method) is the sorting method used, sorting occurs at early adult emergence (pupal isolation and adult cage D-E). Following crossing of these lines, offspring larvae are mass-reared in trays for seven to nine days. On days seven to nine, pupae are isolated from the trays and transferred to adult-rearing cages (or are transferred to a screening cage for the Senecio Robotics technology sex sorting approach) (D-E). Males from the Cas9 line and females from the gRNA line will mate *ad libitum* and acclimate for three days (E). Mosquitoes are then blood-fed by an artificial Hemotek feeder, or by a similar method (Section 2.1.3.9) (F). Two days post blood feeding, water is added to the cage trough for egg laying. The following day, the eggs are harvested and distributed to the field (G). The pure-bred lines are used to create the next generation of the parental line and this repeats the cycle at the facility(G). This is key to having continued production.

Maintenance and Ramping Phases have the same Factory Stages and do not have Release Stages.(**Release Stages**)-The egg delivery to the release sites will be done by drone or by other vehicles (H). Once distributed in the field, the larvae will be raised in shallow trays to adulthood, when they mate with wild female mosquitoes (I). This Active Phase production is continued for 12 weeks whereby modeling predicts localized extinction of *A. gambiae* (J). Figure generated in BioRender.com.

##### 2.1.3.2 Egg isolation and hatching equipment costs

Egg isolation requires minimal equipment. A simple metal filter can catch the eggs drained from the trough (FAO/IAEA 2017), or the eggs can be laid on biodegradable filter paper. Eggs from colony maintenance intercrosses within the parent lines will be used to maintain the colony, while eggs from the Cas9 and gRNA crosses will be transferred to the drone delivery system for distribution. At peak capacity, 70-100 cages will be harvested for eggs each day, 3% of which will support colony maintenance. The IAEA and cage suppliers do not provide the exact volume of water in the trough, but looking at the dimensions provided, we estimate two liters per cage. Therefore, 4.2-6 liters of water will be harvested for colony maintenance, while the other 135.8-194 liters will contain pgSIT eggs for processing and field release. These procedures are low cost requiring only plastic buckets, metal sieves, and optionally filter paper to load into drones(FAO/IAEA 2017). Ten-liter buckets can cost about 2 USD apiece, and 40 buckets would capture all the water before filtering the eggs. Therefore, the startup cost of egg isolation is quite minimal (80 USD). Metal sieves would also be inexpensive, with a 38µm sieve costing around 80 USD. The filter paper may not be required and would also be affordable, accounting for a similarly insignificant cost of 1,000 USD per year (84 days of releases multiplied by 10 USD for a pack of large filter paper per day).

The COPAS FP-500 is a larva sorting technology available from Union Biometrica that can sex sort the parent Cas9 and gRNA lines, prior to the production of sterile male offspring. To prevent cross-contamination of the parent lines, larvae can be run through a COPAS FP-500 exclusive to that line of mosquitoes. The fluid dynamics and precise channel sizing result in a single larva flowing through the chamber at a time. Lasers excite the sex-specific fluorescent markers expressed by the larvae to identify the sex of the larvae. If the larva expresses the sex-specific marker of interest, the larvae are diverted and recovered in a bulk vessel to be distributed into mosquito-rearing trays. If the larvae are not the sex of interest, they can be sent to a negative sorted bin for colony maintenance or disposal. Additional options for sample recovery could be used in the rare cases of double feeds into the laser. COPAS sorting can also be used for quality control of the lines by also identifying line-specific transgene expression markers. Additionally, COPAS can be used to count numbers of larvae in the absence of markers if a general quantification is required, such as to normalize the number of larvae in each rearing tray. Figure generated in BioRender.com.

##### 2.1.3.3 COPAS sorting for pgSIT

The COPAS sorting device separates larvae based on fluorescent markers (**Fig. 7**). It can detect the loss of transgene fluorescent markers, which can indicate unexpected loss of the transgene, line contamination, or can be used for sex sorting L1 larvae to set up the Cas9 x gRNA line crosses needed to generate sterile males. Sex sorting can ensure offspring receive Cas9 paternally and gRNAs maternally, the optimal cross-directionality for the current system to function. There can be higher mortality in the *A. gambiae* offspring, for example, when the Cas9 is inherited from the mother (unpublished Akbari lab). pgSIT crosses have also shown better efficacy depending on which parent mosquito provides the Cas9. During the active release phase, daily sex sorting will continually generate more production crosses. During the maintenance phase, the COPAS will primarily be used to periodically screen for the unexpected loss of transgenic markers or contamination. Upon completion of the COPAS sorting, the larvae will be transferred to the rearing racks and trays.

##### 2.1.3.3a COPAS FP 500 Initial Cost

Sex sorting is a key technology for pgSIT implementation as it is the rate-limiting step. Without high-throughput sex sorting technology, the facility will not be able to achieve production levels required for large scale releases. Additionally, these machines are the most expensive equipment required for the facility. Due to the importance of this technology in the pgSIT production workflow, it is important to build redundancies to account for machine malfunction, repairs, and general maintenance activities. To maximize egg production, COPAS flow sorting will be run 24 hours a day during the Active Phase, which should meet our production numbers with the minimum number of machines. We will also have multiple sets of parts onsite for repairs and a spare machine to maintain production levels during repairs. Union Biometrica, the producer of these machines, has given us a preliminary quote for unit cost and a contract for repair parts that will provide all of the common replacement parts. The COPAS FP 500 single laser system with an air compressor for sorting capability will cost 308,174 USD per unit, and the dual laser system will cost 358,898 USD per unit. The dual laser system is desirable for screening multiple fluorescent markers, which is important for the quality control of the parent lines. Union Biometrica offers an annual maintenance fee of 12% of the initial machine cost. This maintenance fee will provide all of the minor and common parts needed to make repairs and remote technical support. They also recommend having spare lasers on site as these are the most expensive components and have a longer lead time than standard parts. The laser costs are included in the annual contract, but a spare onsite laser can avoid long term disruptions in production. Spare lasers cost 46,000 USD each, and it is recommended that we have one spare laser onsite for each type of laser (e.g. two spare lasers are required for the dual laser system).

Each COPAS FP500 can sort up to 1,200,000 larvae daily, yielding approximately 600,000 of the desired genotype if run for 24 hours. Typically, these machines require cleaning between runs, so a more reasonable run time is 23 hours a day for a total of 562,500 larvae per machine. To have a release ratio of 32 eggs per wild adult mosquito, the number of larvae that need to be sorted daily to meet peak Active Phase production levels varies from 81,840 to 175,394 larvae (**Table 4**). If we have efficient fecundity (300 eggs per female), we need to sort 81,840 L1 larvae a day, which requires a fraction of the capability of a COPAS FP 500 machine. Similarly, using the same calculations, if the fecundity of the mosquitoes is more moderate (210 eggs per female) and the survival to adulthood is lower (50%), we will need to sort 175,394 L1 larvae per day, still requiring less than one COPAS FP 500 machine. In order to maintain a functioning facility even in the event of a machine failing, it is likely preferable to have two COPAS FP 500 sorting machines available. This cost, plus two spare lasers, results in a total cost of 948,840 USD for the low fecundity group. These estimates are based on discussions with Union Biometrica, which indicate a sorting rate of 25,000 L1 larvae per hour. During the Active Phase, we plan to maximize production and assume we can sort 575,000 L1 larvae for 23 hours per day. This production level will last the 12 weeks of the Active Phase (**Section 2.1.4**). There is some uncertainty in these production estimates, as these devices have yet to be evaluated or stress-tested at this production rate for that long duration and with that degree of uninterrupted continual use. We plan to test the production capabilities of the COPAS FP 500 in the early phases of field testing to optimize their use and determine their long-term sorting capabilities. The discussion includes additional costs that factor in doubling the COPAS FP machines, and this is taken into account for initial and annual costs (**Tables 29, 31, 48, and S6)**.

Additional costs are associated with the shipping and importing of this equipment to The Gambia. An approximation of these costs, estimated by trade organizations, was at most 25% of the equipment costs, which is included in the total upfront costs estimated in **Table 5**. Since this equipment is used for disease prevention, perhaps reduced import tariffs and taxes commonly afforded to medical equipment can be negotiated with the government to minimize these costs, but currently, 25% is our best estimate of these costs(Framework n.d.).

##### 2.1.3.3b COPAS FP 500 annual cost

The annual maintenance costs for the COPAS are 12% of the initial machine costs. This cost includes necessary maintenance supplies and replacement parts. This service fee will allow us to keep most general parts onsite and restock as needed, except for a spare laser, which is factored into the upfront cost. Restocking of the lasers will be included in the service agreement, but to avoid production delays due to sourcing and shipping, we will keep at least one onsite.

##### 2.1.3.4 Senecio-Robotics mosquito sorting for pgSIT

While other mosquito sorting options exist, we have selected the COPAS sorting devices because they minimize costs and optimize production. COPAS uniquely sorts mosquitoes while they are L1 larvae, eliminating the costs of raising unneeded mosquitoes past the L1 stage. These costs include larvae food, water to rear larvae, rearing equipment (i.e. trays, rack systems), facility space, and the labor required to rear extra mosquitoes to the adult stage, which is the sex sorting stage for most other sex sorting technologies. These additional costs would essentially double the annual costs for the larval activities, and the facility would essentially need to double in size to rear twice as many mosquitoes to adulthood. To reduce cost and optimize production, it is advantageous to use a COPAS or a similar system that sex sorts at earlier life stages.

Nevertheless, adult stage sex sorting technologies are currently available, so it is prudent to evaluate the cost effectiveness of a commercially available adult stage sex sorting technology. Senecio Robotics (Tel Aviv, Israel) uses an artificial intelligence (AI)-based platform to visually identify and sex sort adult insects. Discussions with this manufacturer revealed that custom sex sorting machines can be built to support pgSIT sex sorting, but do have the drawback of having no sex sorting capabilities prior to the adult stage.

Importantly, the current pgSIT design developed and trialed in *A. gambiae* does not support COPAS sorting due to the exclusion of sex-specific fluorophores. This limitation can be remedied in several ways, but release of the current iteration of the pgSIT technology (lacking the sex-specific fluorescent markers) would be lower throughput or require a Senecio Robotics-like optical sorting technology. However, because the next iteration of the pgSIT with sex-specific markers is actively under development, we have planned for a COPAS-based facility bespoke for this upcoming, more optimized system.

##### 2.1.3.5 Larvae rearing rack and tray system

The IAEA has developed a mass-rearing protocol that we utilized to calculate our pgSIT rearing costs. For larval rearing and feeding, the Wolbaki mass rearing rack and tray system is commercially available and can rear approximately 5,000 larvae per tray and contains 100 trays per rack (**Fig. 8**). These racks can be filled with water from the top with an overflow system that will fill the trays below as the tray above fills. These trays are required for mass rearing to provide the mosquito larvae with the shallow water required for larval development. However, *A. gambiae* can be sensitive to being reared in different types of plastics, so trials will be undertaken to verify their survival in these trays before purchase. These larvae will be fed daily a food mix that has been shown to be effective for rearing *A. gambiae* larvae comprising three types of commercially available fish food. A cheaper alternative food mix will be discussed later in this paper (**Table 11**). The feeding schedule is based on the IAEA standard (FAO/IAEA 2017) but is staggered one day to allow unfed larvae to be sorted by COPAS. On days seven through nine post-hatch, it is expected that the larvae will start developing into pupae. Pupa will be sorted and loaded into adult cages, but the mosquito larvae will be returned to the trays for continued growth until day 9. On day 9, depending on need any remaining larvae will either be disposed of or strained out and added to new trays. If mosquito rearing is optimized, however, few larvae should remain by day 9.

**Fig. 8:**
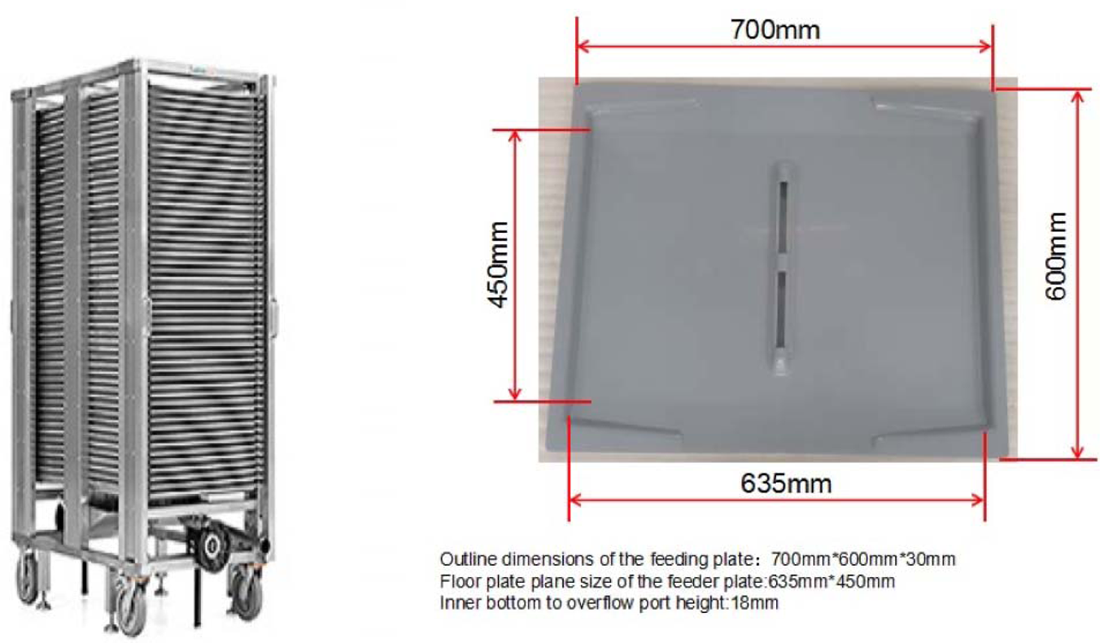
Wolbaki mass rearing rack and tray system.

Production estimates need to consider the mosquito larvae to adult survival rate. The IAEA system uses a predicted larvae to adult survival rate of 50-75% when raised at a density of one larva per ml (FAO/IAEA 2017). While it is possible to test larval survival in our current facilities, due to differences in the environment and procedures in our laboratory and the proposed facility, experiments will be required to ascertain and optimize larval survival during scale-up. Furthermore, the final pgSIT line will have been introgressed into the genetic background of the local native population, which may behave and develop differently altogether. During field trials, we will also better understand the facility’s requirements. The IAEA larvae to adult survival rates range considerably, so until we can assess the survival rate in the facility, we included both the upper and lower IAEA survival rates to estimate the rearing costs. This survival rate will influence the daily sorting rate discussed in the COPAS section and the number of rack and tray systems needed to support larval rearing.

The mass rearing system from *Wolbaki* is the most cost effective commercially available mass rearing system compatible with *A. gambiae* production. It consists of a metal rack and 100 larval rearing trays, each capable of rearing over 5,000 larvae. Each tray has a drain in the center that prevents overfilling and will fill the racks beneath. This overfill may be blocked if there are concerns about cross contamination or control of potential bacterial blooms. This rack system also can tilt the trays in a controlled manner and pour off pupae for isolation(Wolbaki n.d.). The tray dimension image was sent to us via email by Wolbaki.

##### 2.1.3.6 Larvae rearing rack and tray system costs

The most promising mass larvae rearing system was from Wolbaki, which priced its rack and tray system at approximately 22,500 USD. Other companies had higher prices for similar scale systems, while others likely have cost-effective rearing systems but have not been forthcoming with pricing despite communications over several months. To ensure we had multiple, cost-effective options for larval rearing, our next approach was to get piecemeal cost estimates for individual components that comprise a larval rearing rack and tray system. The thermoplastic trays with similar dimensions sell for about 0.99 USD per unit. Acrylonitrile butadiene styrene (ABS) is a common food safe thermoplastic plastic that is inexpensive and can be utilized for this purpose. The racks, while a bit more complex, are aluminum framed and hold 50 trays. Similar aluminum racks have been priced at multiple distributors for around 500 USD. These racks, however, are missing a tilting mechanism that allows water and eggs to be easily removed. Hence, production estimates without this feature must consider alternatives for water removal and egg collection. A simple ratcheting device or similar, which could be placed under the wheels, could achieve this. While a more affordable rack and tray system may become available to meet mass rearing demands, for now we will evaluate the Wolbaki rack and tray system as this is currently commercially available. If lower cost options become available, there will be opportunities to include these options in future facility planning.

Using the 22,500 USD unit price and our known egg production needs (**Table 4**) we can estimate the number of required units and their costs. To estimate the necessary racks, we first take the number of larvae reared per day and divide this by the number of larvae a rack can hold. The Wolbaki rack has 100 trays each that can hold 5,290 larvae per tray, so 529,000 larvae per rack. Larvae are in these trays for up to 8 days. To account for breakage and cleaning time, however, sufficient racks and trays for up to 9 days are beneficial. Cleaning can be done manually, or a machine washing system could be purchased. Therefore, the minimum cost to procure the rack and rearing systems is 45,000 USD. COPAS FP 500 sorting has the advantage of sorting L1 larvae rather than sorting mosquitoes at the adult stage, cutting the rearing numbers in half compared to Senecio Robotics. COPAS’s USD maximum rack and tray cost assumes the lowest fecundity and survival conditions. Senecio Robotics rack costs would be double that of COPAS FP 500.

Additionally, we factored in a 1% annual maintenance cost for the rack and tray systems. The metal rack is the most expensive but sturdy part of the system, so minimal rack repairs are expected with proper use. The trays, on the other hand, are expected to be damaged more frequently but are inexpensive thermoplastic trays that should be replaceable at 1 USD per tray. Therefore, we assume a 1% annual maintenance cost to replace broken trays and for rack repairs (**Table 6)**.

**Table 6:** COPAS sex sorting rack and tray numbers, cost and expected maintenance fees.

| Conditions | Daily Number of Larvae Sorted | Daily Racks | Total Racks in Use | Total Racks (Round Up) | Cost USD | Annual 1% Maintenance Cost USD |
| --- | --- | --- | --- | --- | --- | --- |
| High Fecundity, High Survival Calculation | 81,840 | 0.15 | 1.2 | 2 | 45,000 | 450 |
| Low Fecundity, High Survival Calculation | 116,915 | 0.21 | 1.7 | 2 | 45,000 | 450 |
| **High Fecundity, Low Survival Calculation** | 122,761 | 0.23 | 1.8 | 3 | 67,500 | 675 |
| **Low Fecundity, Low Survival Calculation** | 175,394 | 0.32 | 2.61 | 3 | 67,500 | 675 |

##### 2.1.3.7 Water and larval food requirements

Water and larval food requirements are dependent on the selected larvae rack and tray system. The greatest usage of these resources is during the Active Phase as described later in this report. Still, it is important to account for the Maintenance Phase and Ramping Phase when estimating the usage of these inputs. The Maintenance Phase represents most of the year and is when a minimal colony of the parental lines is maintained. With this in mind, we calculate the use of one mass rearing rack and tray system every three weeks. The mass rearing rack is likely far beyond what is needed, but we preferred to overestimate the numbers to guarantee that we maintain some genetic diversity rather than cause a bottleneck that can negatively affect the genetics and subsequent system function. The Ramping Phase is particularly complex as there will be a logarithmic increase in the facility mosquito rearing numbers. We assume that the six week Ramping Phase will require at least 75% of the resources needed during the Active Phase production. While this is likely an overestimate, there could be unforeseen issues to the rapid scaling up to this production level that these extra costs could cover. Finally, the Active Phase is the most straightforward phase to calculate the water and larval feed usage. In the Active Phase, the facility is working at maximum capacity and will fill a set number of larval racks per week (**Table 7**).

**Table 7:** Annual water usage and cost.

| Conditions | Maintenance Phase Water Used Liters | Total Racks Used | Water Usage per Active+ Ramping Phase Liters | Annual Water Usage Liters | Annual Cost USD |
| --- | --- | --- | --- | --- | --- |
| **COPAS**<br>**Sorting, High**<br>**Fecundity,**<br>**High Survival** | 18,020 | 2 | 19,080 | 37,100 | 371 |
| **COPAS**<br>**Sorting, Low**<br>**Fecundity,**<br>**High Survival** | 18,020 | 2 | 19,080 | 37,100 | 371 |
| **COPAS**<br>**Sorting, High**<br>**Fecundity,**<br>**Low Survival** | 18,020 | 3 | 28,620 | 46,640 | 466 |
| **COPAS**<br>**Sorting, Low**<br>**Fecundity,**<br>**Low Survival** | 18,020 | 3 | 28,620 | 46,640 | 466 |

The water cost is estimated by the rate of water usage associated with each phase. There are 100 trays per rack, and each rack can hold about 530 liters of water. To calculate the usage in the Ramping and Active Phase, the total amount of water for those racks (total racks x 530 liters) is multiplied by 2 to cover water loss and other activities that require water resources (e.g. cleaning, sugar water, egg laying water, and larval feed mixing). Due to increased production in the later stages of the Ramping Phase, water use during four of the six weeks of Ramping Phase will be calculated as equivalent to Active Phase usage, leading to 18 weeks at full capacity. This is unnecessary at this release scale, but this Ramping Phase would be useful to establish the colony at the Active Phase levels prior to releases, producing mosquitoes for suppression at other sites, spread production across the week (if the maintenance phase has synchronized production) and establish techniques to expand production when this facility is scaled up for national or regional suppression. The Maintenance

Phase is estimated to require 18,020 liters of water and was calculated by estimating the water usage in 1 rack over 34 weeks and doubling the water usage to account for water loss and other activities. These estimates are then added together and described below in the cost section.

The larval feed differs from the IAEA protocol due to the development of a more cost effective food formula for *A. gambiae(Bimbilé Somda et al. 2017)*. Additionally, this rearing diet selects globally available ingredients that can be affordably sourced in The Gambia. This mixture consists of tuna meal, Brewer’s yeast, and chickpea powder (**Table 8**). It should be noted that this food mix has not yet been tested in the pgSIT line. However, we are confident that with minor adjustments, it will be sufficient to support larval development. This feed can be stored as a powder at room temperature and is inexpensive compared to other feeds due to the lack of bovine liver powder. Each tray requires about 1.5 liters of this rehydrated food per week following the feed rate from the IAEA protocol (FAO/IAEA 2017) and the increased size of the Wolbaki Trays. This approach results in about 150 liters of this feed per rack per week. Using the same logic as the water calculation, the Maintenance Phase food requirement is 5,100 liters. The specific food requirements are calculated in the same manner as water usage without doubling for cleaning and other uses.

**Table 8:** Larval food cost per liter.

|  | **Grams** | **Kilograms** |
| --- | --- | --- |
| **Tuna Meal** | 7 | 0.007 |
| **Brewer's Yeast** | 1.5 | 0.0015 |
| **Chickpea flour** | 1.5 | 0.0015 |
| **Raw Material Costs** | **USD** | **Cost per Liter USD** |
| **Tuna Meal cost per 100 kg** | 80 | 0.0056 |
| **Brewer's Yeast per 100 kg** | 1000 | 0.015 |
| **Chickpea flour** | 328.26 | 0.0049 |
| **Total:** |  | 0.0255 |

##### 2.1.3.7a Water and larval feed annual costs

Water and larval feed costs are essential annual expenditures to maintain production levels. Water costs in West Africa are typically around 0.01 USD per liter (AAAWater 2019). The market costs for the raw material for the larval food costs are approximately 0.0255 USD of dry powder per liter. The larval feed will be stored as dry material and not require rehydration for use. To be consistent with the IAEA protocol, larval feed will often be referenced in volume rather than weight. These estimates can be used to determine the annual cost of larval feed (**Table 9**).

**Table 9:** Larval food requirements and cost.

| Conditions | Maintenance Phase Food Used Liters | Total Racks Used | Food Usage per Active+ Ramping Phase Liters | Annual Food Usage Liters | Cost USD |
| --- | --- | --- | --- | --- | --- |
| COPAS Sorting, High Fecundity, High Survival | 5,100 | 2 | 3,600 | 8,700 | 222 |
| COPAS Sorting, Low Fecundity, High Survival | 5,100 | 2 | 3,600 | 8,700 | 222 |
| COPAS Sorting, High Fecundity, Low Survival | 5,100 | 3 | 5,400 | 10,500 | 268 |
| COPAS Sorting, Low Fecundity, Low Survival | 5,100 | 3 | 5,400 | 10,500 | 268 |

##### 2.1.3.8 Pupae isolation and transfer into prepared adult cages

A key feature of the larval rack and tray system is that the trays can be tilted to easily pour off the contents. All 50 trays in a rack can be poured into a bucket and then be poured into individual Erlenmeyer flasks to isolate the pupae from the larvae (**Fig. 9**). Automated pupae isolation technologies have been demonstrated, but these devices are limited in their capabilities (Fabrizio Balestrino et al. 2011), often not developed for *A. gambiae*’s unique morphology, and are not commercially available. While there may be opportunities in the future for such pupae isolation devices, manual sorting via swirling in Erlenmeyer flasks is an affordable and effective method to collect pupae.

**Fig. 9:**
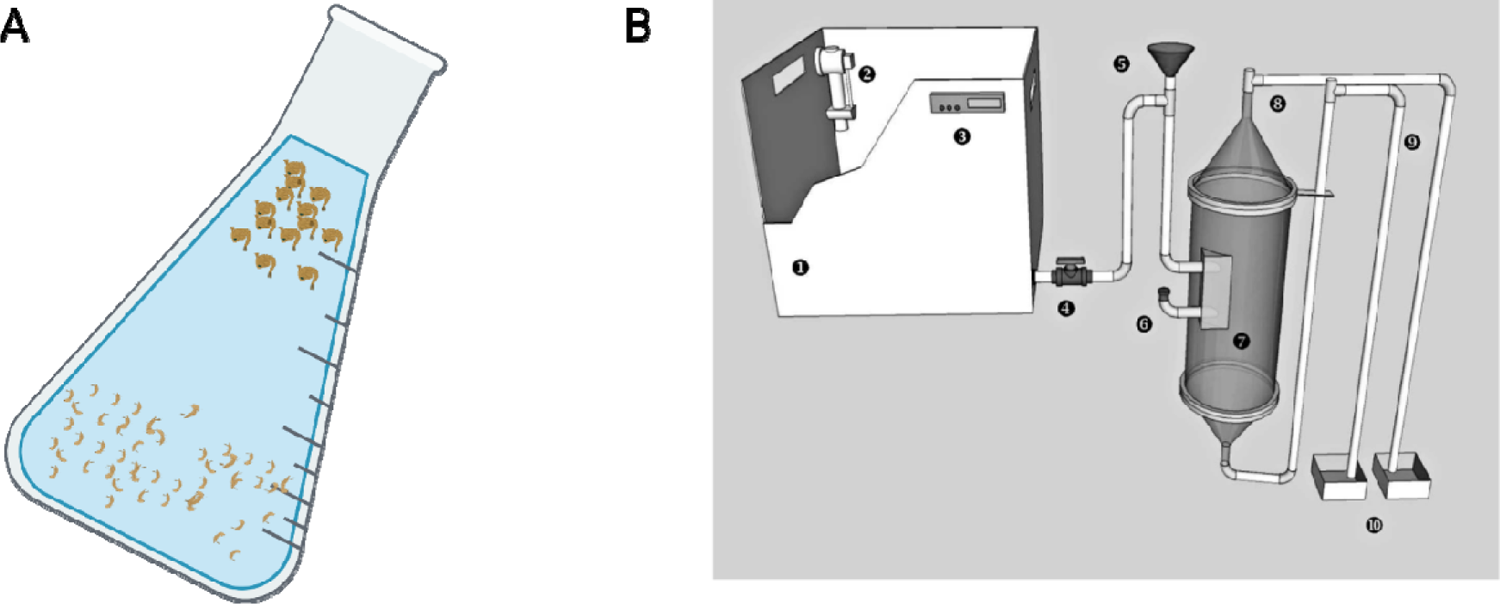
Pupal isolation methods (A) Pupal isolation can be accomplished with an Erlenmeyer Flask. By swirling the flask with a mixed population of larvae and pupae, pupae will become concentrated near the top of the flask. These pupae can then be poured off of the mixture. (B) From (Fabrizio Balestrino et al. 2011) depicts a large mass separator device design. While automated pupae isolation technologies have been made and described in publications, there is a lack of commercially available versions of this technology. Figure generated in BioRender.com.

Pupae can then be measured by volume using a modified 50 ml conical tube with a fine mesh lid as described in the IAEA protocol (FAO/IAEA 2017). This method can be used to estimate the pupae numbers prior to addition to the adult cages. There is a trough at the bottom of the cage where water and pupae can be added. The adult cages will also be outfitted with sugar feeders and Hemoteks, as described previously (**Fig. 10**).

**Fig. 10:**
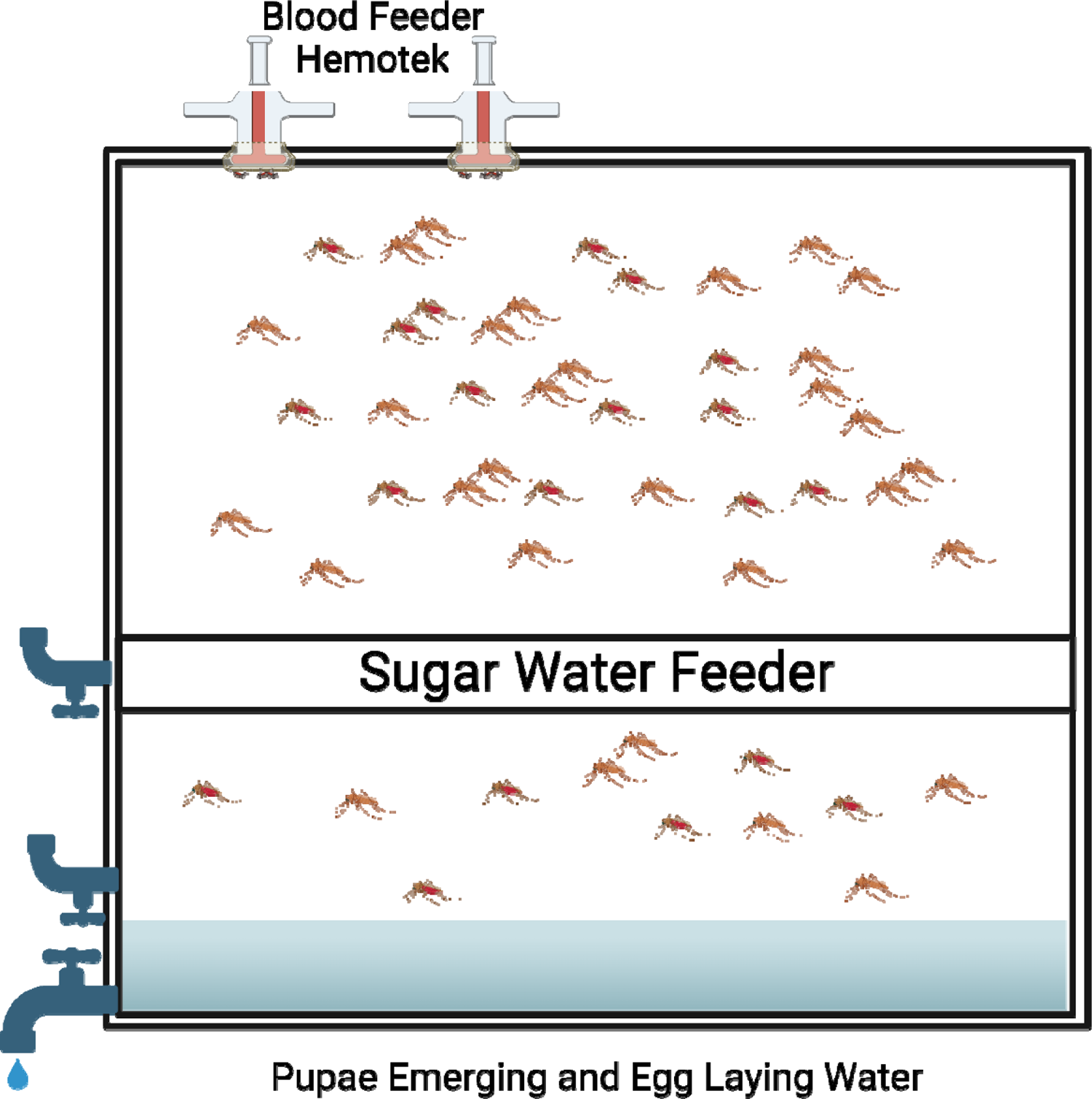
Adult mosquito cage configuration.

The adult cages are thin cubes with the sides covered by screens. Blood feeders are attached to the top to allow for easy blood placement and minimize staff exposure to mosquitoes. A tube system for sugar water also feeds the mosquitoes via a wicking material, as both sexes require a constant source of sugar water to support life. The bottom of the cage has a trough for water input and removal. Water with pupae is added to facilitate the emergence of adults into the cage. Water is then removed until oviposition. Water is added two days post blood feeding for females to lay eggs, and the water containing freshly oviposited eggs is drained off into a filter, which can be distributed in the field or used for mosquito production. Figure generated in BioRender.com.

##### 2.1.3.8a Pupae isolation, transfer, and adult cage costs

One-liter plastic Erlenmeyer flasks can be purchased at 5 USD per unit. Considering that 2 to 3 racks will need pupae isolation daily and that one worker will be isolating pupae from two racks a day, we estimate that we will need two Erlenmeyer flasks per worker. These estimates result in the total cost for 4-6 Erlenmeyer Flasks at 20-30 USD, which includes costs for separate flasks for each line.

The estimated number of adult cages needed to support the proposed production range from 60 (if female fecundity is high, 300 eggs per female) to 90 adult cages (if female fecundity is lower, 210 eggs per female). Many of the manufacturers charge extra for these cages due to their extra features, but the minimum cost for a unit was estimated by other research groups at 250 USD a unit (Maïga et al. 2019). At this price, 60 cages will cost 15,000 USD, and 90 cages will cost 22,500 USD (**Table 10**). A proposed maintenance cost was 5% due to the material used to create these cages being affordable, particularly the netting which will likely be the common breaking point.

**Table 10:**
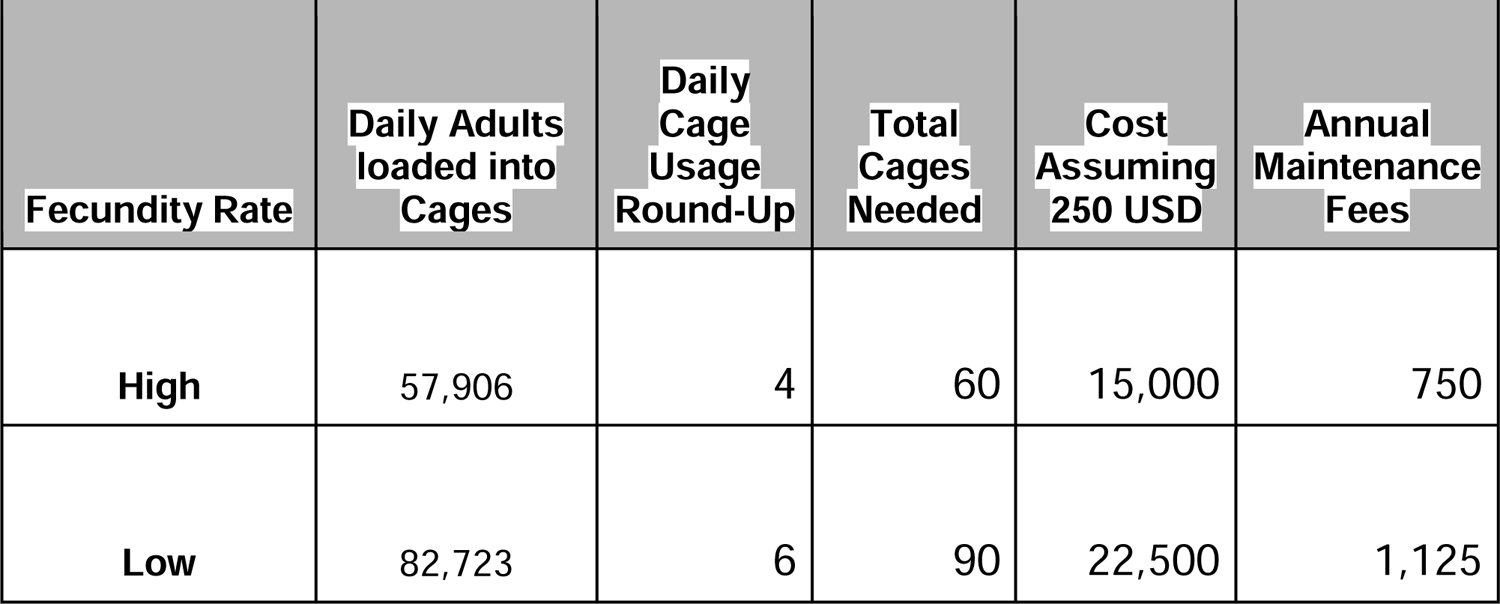
Adult mosquito cage requirements and annual fees.

**Table 11:** Hemotek device cost and annual fees.

| **Requirements and Cost** | **High Fecundity** | **Low Fecundity** |
| --- | --- | --- |
| **Daily Cages** | 4 | 6 |
| **Daily Feeds** | 12 | 18 |
| **Hemoteks Required** | 2 | 3 |
| **Hemotek Cost** | 1,181 | 1,181 |
| **Total Hemotek Cost** | 2,362 | 3,543 |
| **Annual Repair Costs** | 118 | 177 |
| **Active Phase Feeds** | 144 | 216 |
| **Maintenance and Ramping Phase Feeds** | 204 | 204 |
| **Total Feeds** | 348 | 420 |
| **Membrane cost per feed** | 0.03 | 0.03 |
| **Annual Hemotek Costs** | 10.44 | 12.6 |

##### 2.1.3.9 Adult mosquito blood feeding and egg collection

Adult female *A. gambiae* mosquitoes require blood feeding to produce eggs. In the wild, mosquitoes blood feed directly on humans and animals, but within a large-scale facility, this becomes untenable. Fresh blood is the optimal feed for *A. gambiae,* with some attempts to feed frozen blood and lyophilized plasma containing ATP with limited long-term success(Baughman et al. 2017). These options were explored, but fresh blood is the most effective choice and is significantly cheaper when sourced locally (**Table S2**). While human blood is optimal, it has higher biosafety considerations, and sourcing sufficient quantities is difficult. Successful feeding of *A. gambiae* with cow blood has been documented (Lyimo et al. 2012), which we propose as an economically available alternative. Two to three cattle would be required for direct blood feeding a day, but there would be concerns about safety and the ability to handle and maneuver cattle in the facility easily. Live cattle blood sources were considered (**Table S2**), but using artificial feeding devices with drained bovine blood is far simpler and safer. When considering the size of the average cattle in The Gambia, we estimate that the blood from one cow slaughtered each day should be sufficient to support adult feeding during the Ramping Phase and Active Phase, with only a small amount of blood required during the Maintenance Phase. Based on an estimate of 300,000 head of cattle in the nation, local slaughterhouses should be slaughtering a large number of cattle and should be able to meet this demand. Additionally, refrigeration of fresh blood can potentially stabilize this blood for one to two weeks, but this will primarily be used to cover for days fresh blood is not received(Baughman et al. 2017). Feeding on previously frozen or >5 day old blood can cause significant female mortality, so these feeding modalities are not recommended if freshly acquired blood is available. A common brand of synthetic feeding device, the Hemotek membrane feeding system (Hemotek Ltd, UK), uniformly warms blood or blood substitutes within a membrane and attracts mosquitoes allowing for life-like feeding without a host. The device will feed the mosquitoes around 60mL of blood. This approach will require a simple but validated modification to the IAEA protocol to feed mass rearing cages (Damiens et al. 2013; FAO/IAEA 2017).

Two days after blood feeding, water is added to the cage trough to allow the *Anopheles* females to lay eggs on the water. Once a sufficient number of eggs have been laid, the cycle can repeat. 6% of the eggs will be from the Cas9 and gRNA parental intercrosses and will be used to maintain the facility’s mosquito population, and approximately 94% of the eggs will be from the Cas9 x gRNA crosses to be distributed in the field.

##### 2.1.3.10 Upfront adult mosquito blood feeding and egg harvesting costs

The main upfront cost for blood feeding is the purchase of Hemotek devices. While there may be opportunities to reduce the costs in the future and use simpler methods, the Hemotek has been vetted for feeding in many settings, and although expensive, it is a conservative approach for our cost estimate. The basic kit for these devices costs 1,181 USD per unit, but a modified unit with an increased blood feeding surface area is required to feed such a large cage properly. This modification increases the typical blood feeding receptacle from 3 mL to approximately 60 mL. This modification could likely be done in-house for a fairly minimal cost as it only requires larger heating plates and a larger plastic container for the blood meal. The facility requires 35-50 of these devices depending on mosquito fecundity. The total Hemotek equipment costs will be between 2,362 to 3,543 USD (**Table 11**). Notably, there are other means to prepare blood meals that simply require heating the blood and leaving it for direct feeding, which may be lower cost. However, an efficient and reliable blood feeding system is paramount for egg production, and any alternative blood feeding method would have to be extensively tested before implementation at scale (**Table 11 and Table 12**).

**Table 12:** Fresh blood estimate: utilizing locally sourced blood. Pricing is based on blood obtained from food markets in the US. It is expected that fresh blood costs in The Gambia will be less than US prices.

| Fecundity | Total Liters of Blood Per Year | Cost Per Liter (USD) | Total Cost (USD) |
| --- | --- | --- | --- |
| High | 20.88 | <2.24 | <47 |
| Low | 25.2 | <2.24 | <56 |

Blood will be locally sourced, as the costs of internationally shipping blood are too high, and mosquitoes are more optimally reared on fresh blood. There are local slaughterhouses near the planned facility construction site in Banjul. The blood requirement for the facility is about 0.72-1.1 liters per day during ramping or active production. Cattle in The Gambia are primarily *Bos indicus*, Zebu cattle which are about half the size of standard *Bos taurus* Holstein cattle in the United States. American cattle can have approximately twenty liters of blood extracted when slaughtered, so we estimate that Zebu cattle should have about ten liters each. Local butchers may have to be trained on exsanguination protocols to produce blood sufficiently clean for use in the facility. While there is not a direct cost per liter of blood, in The Gambia it is expected that the local slaughterhouses will charge at or below the US market for bovine blood, which is 2 to 4.5 USD per liter, although bovine blood is not often sold in The Gambia (personal communication with Dr. Dalessandro)(B&R Food Services n.d.) (**Table 12 and Table S2**). Due to cattle producing ten-fold the daily requirement when slaughtered, exploring refrigeration options for long term storage will be desirable to increase efficiency if cattle are not being slaughtered daily. It is expected that blood should remain usable for out to five days, but this will have to be confirmed in the context of a mosquito production facility. It is expected to cost less than 3,000-4,000 USD per year to source the blood needed to maintain the colonies (**Table S2**). Other options have been explored, although this is likely suboptimal when compared to sourcing local blood. The cost of blood from non-local sources increases rapidly due to cold chain transportation requirements or freeze drying plasma, which will also not be as effective for feeding mosquitoes. Experts strongly urge the use of fresh blood.

##### 2.1.3.11 Monitoring pgSIT Line Stability

To sterilize mosquitoes for the pgSIT system, it is imperative that the Cas9 and gRNA genetic elements remain stable and effective. In order to estimate the frequency mutations are expected to occur in these key elements, a background mutation rate from closely related *Anopheles* species in the *An. gambiae* complex can be used to estimate the *An.gambiae* mutation rate. Using the spontaneous base substitution rate of1×10^-9^ per site per generation for *Anopheles coluzzii* (Rashid et al. 2022), and the number of bases in the Cas9, gRNAs transgenes and target sites we can determine the likelihood of transgenes of target site mutation. The Cas9 line is inherently more vulnerable to this as it is a larger gene and mutations in most parts of this gene could deactivate this element. The gRNA lines have some resilience as they have inbuilt redundancy in the form of multiple gRNAs for each target gene. A mutation in one of the gRNAs would not result in inactivation of the line. For the gRNA line to be inactivated, all three gRNAs, target sites or some combination would have to be mutated sufficiently to prevent targeting. Not only are these very small sites, gRNAs can tolerate some mutation further reducing the target that would have to be mutated and the target sites are in critical genes that are unlikely to tolerate mutation. The likelihood of this occurring is even less likely as it has to occur on the allele with the original mutations. The chances of this become astronomically small and therefore we focus on the Cas9 mutation rate as this line will have mutations far more frequently. To calculate the Cas9 mutation frequency, we multiply by the number of mosquitoes per generation to achieve the chance of a deactivating mutation occurring per generation (**Table 13**). We would therefore expect 17 generations before a mutation would occur in a Cas9 allele. There are 11 generations in the maintenance phase, and its last generation will begin daily, increased egg production. The ramping phase is approximately 5 generations and the active phase 4-6 generations. Even though the Active Phase has daily mosquito generations, they are still on a 3 week cycle, so we would expect 17-18 generations of maintenance annually. For each of these generations, the same maintenance line amount will be utilized to maintain the stock. Based on the 1×10^-9^ per site per generation mutation rate and assuming 18 generations, we would expect about 1.08 mutations annually in the maintained Cas9 stock of mosquitoes. With minimal fitness cost in the Cas9 lines, it is assumed that these mutations will not undergo positive selection. Additionally, the fitness cost between unmutated and mutated may be especially minimal as a Cas9 protein with a single point mutation will likely have similar fitness costs. At this mutation rate, and assuming a neutral fitness cost of the mutation, Cas9 mutations should only affect 1% of the stock population at about 56 years post-production.

**Table 13:**
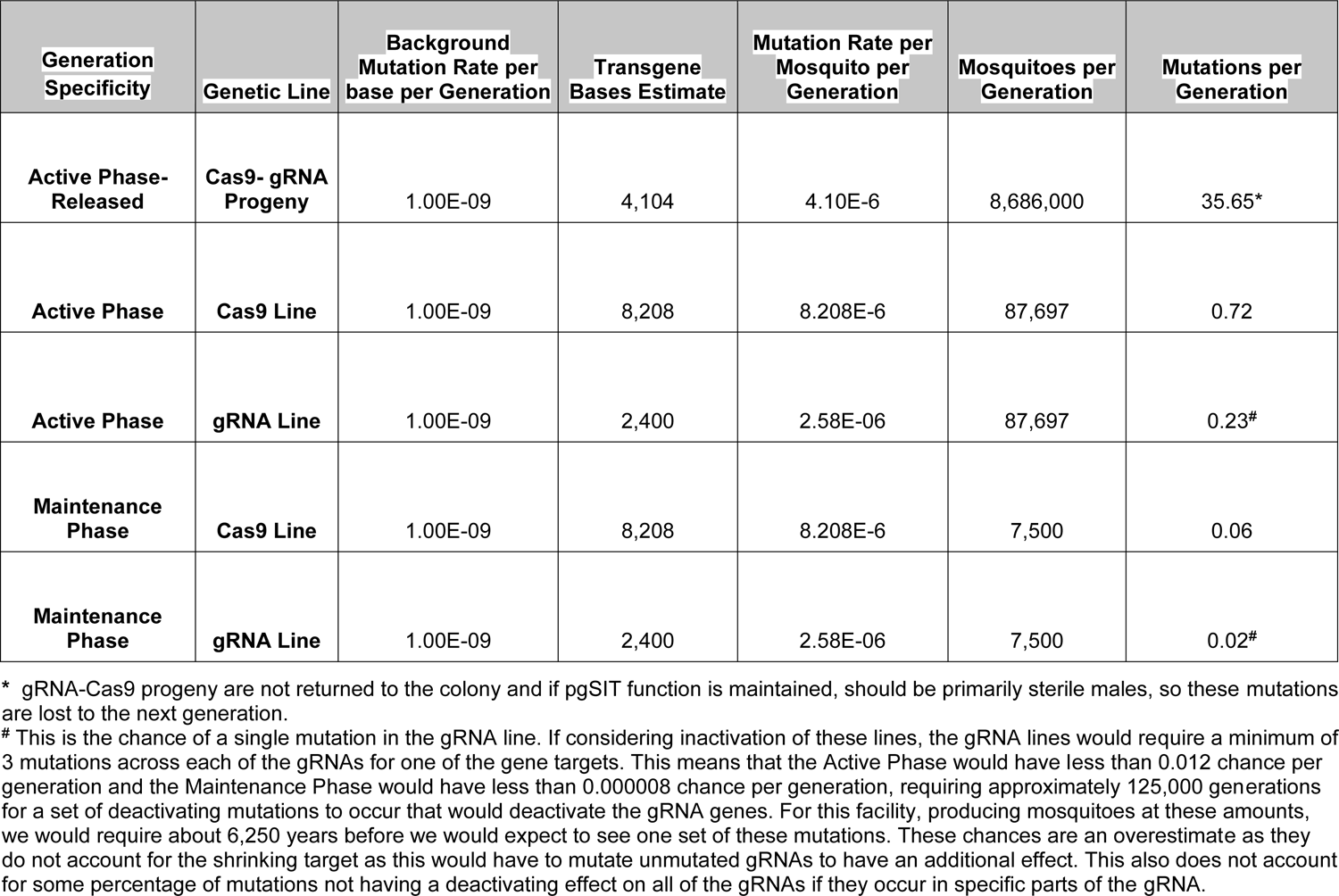
Mutation Rates and Monitoring Assessment: The background mutation rate is based on estimates from another member of the *Anopheles gambiae* species complex, *Anopheles coluzzi* (Rashid et al. 2022). To estimate the expected mutation rate per generation, we use estimates of the genetic element base lengths and the total number of mosquitoes per generation. These calculations can then be used to determine mutation rate over time in the maintenance and active phase, and the expected mutation rate in the Cas9-gRNA pgSIT offspring.

Further work is required to more accurately estimate the mutation rate and selection for transgene and target site mutations. However, these estimates can be improved during the earlier field trials. Selecting lines that have minimal fitness cost can also minimize the risk of loss-of-function mutations accumulating in the stock lines. Quality control procedures can also be implemented to identify and remove mutations that reach high enough frequencies to impact pgSIT performance.

Early field and mass production trials can include monitoring to evaluate and reduce the release of unedited eggs. The pgSIT release and production plan uses 6% more parental mosquitoes than needed for production, which is three times the maintenance requirement. Some of the extra mosquitoes can be used for quality control efficacy and mutation procedures. Cas9 function and the fidelity of the male survival gRNAs can be evaluated by confirming the sex specificity of the sterile males with the COPAS FP 500 sex sorting procedure. To verify sterility the quality control protocols can include periodic sequencing of the editing sites and a portion of the Cas9 x gRNA offspring can be mated with wildtype females to confirm sterility. These additional quality control procedures should not have a large impact on cost.

##### 2.1.3.12 Drone delivery of pgSIT eggs throughout the Upper River region and delivery considerations

About 8,686,000 eggs will need to be flown to the eastern part of the country within 48 hours of laying to ensure the eggs remain viable and fit for release(Mazigo et al. 2019). To maximize efficacy, eggs will be delivered by drone on the day of harvesting or the next day. The predicted dry weight of these eggs is 37.36 grams. However, *Anopheles* eggs cannot be desiccated at any time, so they must be shipped with moisture which will add weight. Drone engineers at Arda (https://www.ardaimpact.com/) have advised on drone egg distribution. They concluded that long-distance drone delivery to the eastern parts of The Gambia (the URR and beyond) is feasible from the capital city of Banjul with current technology (**Fig. 11**). The gasoline-powered drones they have are capable of carrying a 10 lb payload up to 560 km per trip, which means that this can serve much of the far east of the country from the port city of Banjul. We are discussing increasing fuel capacity in exchange for the payload to reach the far east border comfortably. The advisers at Arda predict that a longer range can be achieved by reducing the payload in exchange for increased fuel. The daily delivery will likely be light enough for increased fuel capacity.

**Fig. 11:**
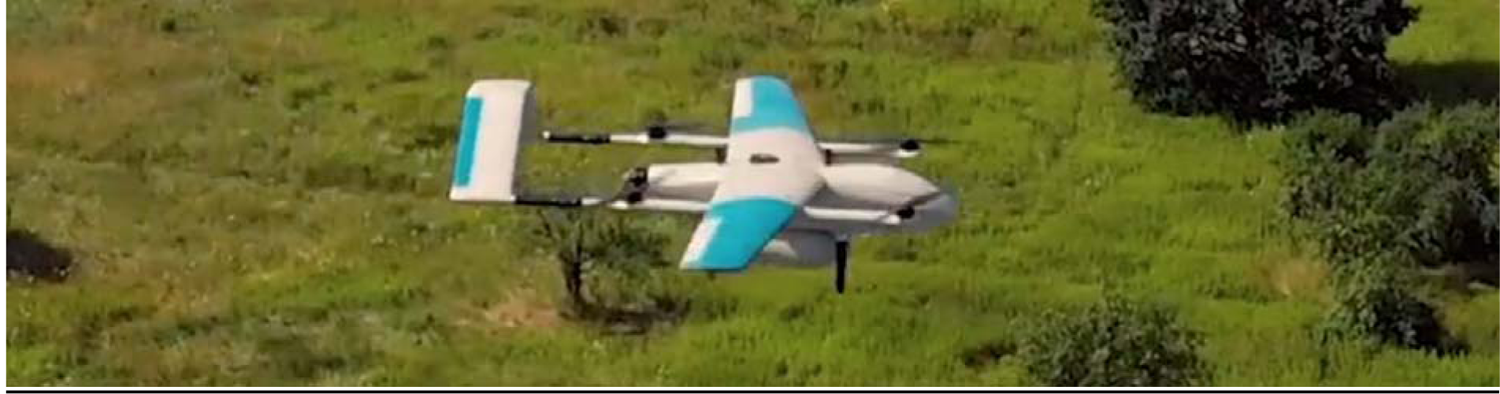
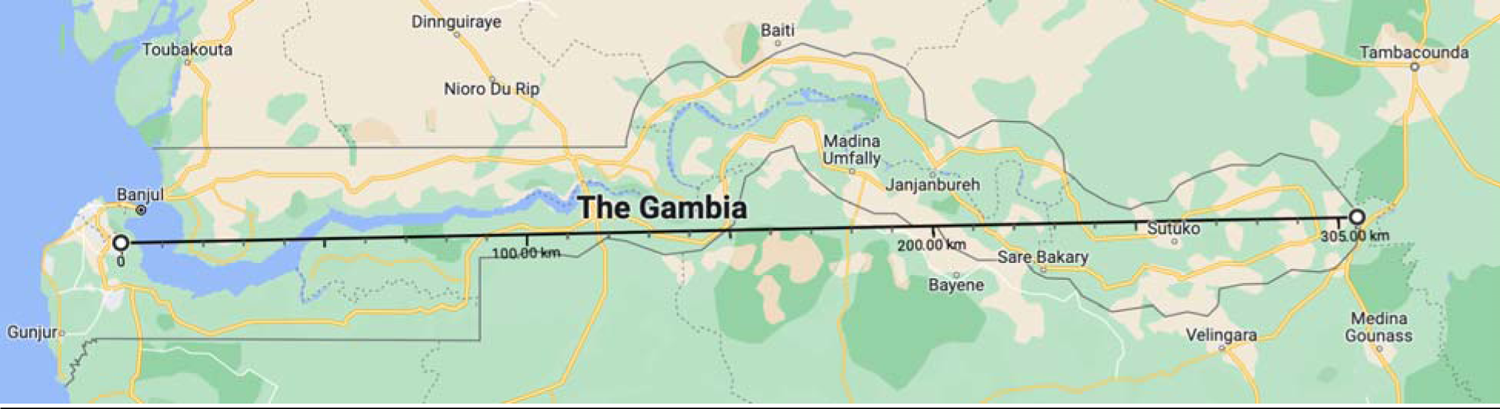
Arda drone and drone flight range with standard fuel tank and range.

Delays in egg delivery should also be considered in the cost estimates. While it is optimal for the facility to keep a daily schedule, inclement weather, drone repair, flooding preventing ground operations, and other issues may cause an interruption in delivery services. Fortunately, *A. gambiae* eggs tolerate cold storage at 4°C for up to 5 days at high density with no noticeable hatch changes (Mazigo et al. 2019). This feature allows for some flexibility in the delivery schedule. Nevertheless, delivering the mosquito eggs within the next couple of days is likely necessary to retain egg viability and consistency in the suppression plan. Importantly hatched nascent larvae can survive at ambient temperature for an additional day, potentially expanding the delivery window to three days if necessary. With the potential requirement of doubling the typical delivery rate over a couple of days, it is important to have excess delivery capacity to provide coverage.

Arda drones are long range with vertical take-off and landing (VTOL) capability for easy take-off and landing without a runway, making them convenient for use in the more urban areas around the facility in Banjul and the more remote areas in the Upper River Region(Arda n.d.).

##### 2.1.3.13 Costs of drone delivery of pgSIT eggs in the Upper River region, storage costs and delivery considerations

The number of drones required can remain constant as these have a set weight capacity and, therefore, a set egg delivery amount per day. Additional flights can be scheduled to accommodate larger egg distributions. Each drone costs about 25,000 USD, and while these are expected to need minimal maintenance according to the vendor, we are estimating two flights a day with two drones in reserve in case of malfunctions in the primary drones or the need for doubling the delivery rate for a day. This strategy gives us an upfront cost of 100,000 USD for four drones. This estimate has factored in ground support and other features that should allow for autopilot flight and flight monitoring (**Table 14**).

**Table 14:** Drone costs and annual fees.

| **Total Drones Used** | **Initial Drone Cost USD** | **Gallons of Gas per Flight** | **Annual Flight Numbers** | **Total Fuel Used Annually** | **Fuel Cost Annually USD** | **Maintenance Fees Annually USD** |
| --- | --- | --- | --- | --- | --- | --- |
| 4 | 100,000 | 0.75 | 196 | 147 | 657 | 10,000 |

The storage of eggs in case of delivery delays also has costs including a large refrigerator to store the eggs at 4°C (Mazigo et al. 2019). Commercial refrigerators cost approximately 1,000 to 3,000 USD, depending on the storage capacity required, but it is likely safe to assume an average cost of 2,000 USD. Additionally, a gasoline generator should be procured for the facility to maintain electricity for refrigeration and other technologies in the facility. For a small factory, a 10,000-20,000 USD generator should suffice. The facility should not require storage of more than five days of eggs as their hatch rate declines substantially after that time point (Mazigo et al. 2019). However fresh eggs are always preferred.

##### 2.1.3.14 Drone delivery alternatives

There are drone systems currently in use in Africa. In particular, Zipline has been using drone technology to deliver medical supplies throughout Africa. Their drone utilizes a catapult and a hook system to minimize the launching and landing space needed for these drones. While this is effective for their purposes, current specifications of these drones from Zipline limit their total flight distance to approximately 161 km. We would require approximately 610 km of total flight distance. It is possible that their drones can be modified to have an extended flight range, and this will be investigated as an option for delivery services if this project moves forward.

Other companies offer similar drones with this flight capability and, most importantly, range. The best approach would likely be to contact multiple companies a year or two prior to the completion of the facility to find the best options for the facility. Regulatory testing will still require around five years to complete, so it is foreseeable that better optimized drones will be available when the facility is constructed.

Additionally, there are options to deliver the eggs via motorized vehicles. These could be included as an option if there is an issue with drone delivery, or as an emergency backup in case of drone malfunctions. Roads in the region, however, are often under maintained, so ground transportation is not optimal for delivery.

##### 2.1.3.15 Mosquito release in the Upper River Region

Drones will deliver the eggs to release sites throughout the Upper River Region where they will need to develop from eggs to adults. There are some native habitats where wild mosquitoes lay their eggs, but these are unlikely to support the development of such a large number of mosquitoes with the exception of rice paddy fields. The likely solution will be to engage and hire members of the local communities to support an onsite mosquito rearing system for the sterile males. This approach would provide an opportunity for community engagement, which is a correlate of public acceptance and program success (Diepeveen et al. 2013). These systems would require a large source of shallow water as a larval habitat which, if not managed appropriately, could provide habitat for wild mosquitoes too. Engineering solutions can be devised, such as a net system or lid that prevents entry of wild mosquitoes, but could be opened briefly when the pgSIT sterile males have reached adulthood. Or a baffling system can be designed to allow adult mosquitoes to exit but not enter. Similar systems have been built for many mosquito trapping and release devices, but numerous other solutions can also be tested as the project is scaled.

##### 2.1.3.16 Mosquito release in the Upper River Region costs

This system would require several community members per site to supply larval food and maintain water levels in the larval habitat; however, the training required for these activities is minimal. Water can likely be supplied by rainfall or other local water sources. The annual costs will consist of training, worker salaries, and the delivery of larvae, feed, and water for rearing.

The primary upfront costs include training, network connections, and rearing trays. Training costs will involve community outreach efforts to gain community feedback and acceptance of the pgSIT program and to train community members to manage their local mosquito releases. Current outreach programs for mosquito net usage and insecticide residual sprays cost approximately 9,000 USD for The URR. The training and community outreach activities for this project are unique compared to other technologies employed in the region. Outreach for pgSIT may be more complex since a genetic technology may be harder to communicate to a broader local audience than the concepts of LLINs and IRS. It is foreseeable that convincing communities to rear mosquitoes (albeit sterile males) to suppress mosquito populations could appear contradictory and more difficult to gain approval compared to LLINs and IRS. However, pgSIT may be simpler to implement for community-wide protection, as it only requires compliance and assistance from a few community members. For example, the rearing of pgSIT mosquitoes is only done by a small percentage of the population, in contrast to the broad community compliance required for LLINs. With these operational considerations, we include two outreach and training cost estimates. The lower estimate (18,000 USD) doubles the current 9,000 USD LLIN and IRS outreach budget. A more conservative estimate per Dr. Umberto D’Alessandro is 500,000 USD, which includes a full team of social scientists and community leaders. Both values are accounted for in **Table 25 and Table 26**, respectively. However, we expect that the cost will drop significantly after validating pgSIT for large scale suppression. Additionally, each field rearing manager would need some means to communicate with the facility in The Gambia. Mobile phone ownership is about one per person across the country, so upfront phone purchasing costs may not be necessary, but phone services will be included in monthly fee calculations (Info 2021).

The final upfront cost is the mass rearing trays for the release sites in The Upper River Region. Thermoplastic ABS trays are inexpensive, likely around 1 USD per tray (**Section 2.1.3.6**). There is the potential to create netted troughs or small ponds for field rearing, but this will require experimentation to determine if this cost saving approach is viable. Larvae may have the potential to feed off of native food sources in these small troughs and ponds. The weekly rearing requirement is linked to the weekly water requirements to rear these mosquitoes, which we can calculate from the weekly egg release numbers. The estimated release schedule is just under 61 million eggs per week, only half of which will hatch because pgSIT females die shortly after hatching. This release schedule means about 30 million larvae will need to be reared to adulthood in The URR weekly. Using a similar density of mass rearing, we assume a rearing density of 1 mL per mosquito larva. These field trays or ponds are expected to have sufficient rearing capacity to achieve this goal as the modeling is built around only 26% of eggs surviving to adulthood, and larvae at this density are expected to have 50-75% survival to adulthood (**Table S1**). The 26% survival is derived from the expected survival as eggs, larvae and pupae in the model. The URR requires 30,000 liters per week to rear the larvae. If we use a tray similar to the Wolbaki trays, we have 5.3 liters per tray, which means we need approximately 5,660 trays for the URR. While there will likely be a range of tray sizes available, and there may be more cost-effective food safe plastic bins, we can still apply this cost to assume the high range cost of 5,660 USD for rearing trays in the URR. Trays could likely be replaced with ponds or troughs with netting, suggesting that the price could be lower than this.

Additional annual costs associated with delivery and rearing in the URR include water, larval feed, and labor. This quantity of water required weekly for the 12 week Active Phases is 359,976 liters. At about 0.01 USD per liter, this costs about 3,600 USD annually. The water costs will likely be lower than this as it will be the rainy season, and therefore opportunities to capture rainwater, but we estimate somewhere around this cost for the entire region if water is purchased at market prices. Alternatively, rain capture facilities could be established at release sites to provide ample water for rearing. Dual usage of this water could also save money, but this will require experimentation to determine best practices. Larval feed is calculated using water usage and the 28.3% food per liter of water estimate (FAO/IAEA 2017). The larval feed does not need to be liquefied, but this makes the food calculations easier. Weekly, the URR would need 8,490 liters of larval feed and about 101,880 liters of feed over the entire time. It will cost around 2,600 USD per year to feed the larval at the release sites in the Upper River. Water can be purchased locally, but the food will require transport and distribution. The current distribution costs of LLINs throughout the URR is approximately 60,000 USD annually, and we assume a similar cost for feed distribution. The weekly weight of the food required is 84,900 grams, and should be delivered as dry food. The annual food requirement is, therefore, 1,020 kilograms for the entire Upper River Region. This amount requires prior delivery to the various release sites to guarantee that they have sufficient feed for the mosquito larvae. Artificial rearing ponds and troughs may reduce larval feed requirements since natural food sources are also available. Pond and trough options are likely viable approaches, but we selected the more conservative and expensive estimate in case a more expensive approach is required (**Table 15**).

**Table 15:** The costs of rearing larvae in The Upper River region.

| Upper River<br>Region<br>Rearing<br>Costs | Training,<br>Public<br>Outreach<br>and<br>Delivery | Upper River<br>Rearing<br>Water Cost<br>USD | Upper River<br>Rearing<br>Food Cost<br>USD | Upper River<br>Rearing<br>Labor Cost<br>USD | Total USD |
| --- | --- | --- | --- | --- | --- |
| **Annual Cost** | 60,000 | 3,600 | 2,600 | 78,750 | 144,950 |

The egg releases in the URR will require management on the ground to rear these mosquitoes from an egg to adulthood. Although it is common for anti-malaria programs to utilize volunteer labor that often is not accounted for in budgeting, we wanted to budget these positions as this would provide better incentives for effectively managing these release sites. When these programs become governmentally run, there may be restructuring of local management to volunteer labor. One to two individuals will likely be able to manage mosquito rearing independent of the rearing density and volume. As a rough estimate, we will assume that there will be one rearing site per thousand people, with one local manager, hired part-time to manage mosquito rearing. This estimate may vary depending on human population density and the size of the release site, so this will require further investigation. For the URR, there are about 292,000 people, so we estimate at least 300 field managers spread throughout the region are required for mosquito rearing (**Table S3**). This job would be part-time so we took half the monthly salary for the main facility. While the actual work will be significantly less, it would be useful to have field managers with enough time available to troubleshoot any issues that may arise. With a monthly wage of 75 USD and 14 weeks of release, the cost would be 78,750 USD. This calculation brings the total cost of 145,000 USD to pay for on site releases in the URR. While water and labor may be received at discount prices assuming there will be cheaper water sources during the rainy season and some level of volunteer effort provided by the community, this is not guaranteed. The mosquito rearing and training costs are described in more detail below.

The larvae rearing will follow the same protocol as seen in the mass rearing facility, but only require rearing the mosquitoes to adulthood (**Fig. 12**). Eggs arrive on day one via drone or other delivery method. These eggs are then hatched in a separate container before transferring to the rearing tray. This procedure allows the trays to be used for the full rearing cycle (Days 2-8). The larvae will then be fed according to IAEA protocol, and we estimate a similar development rate resulting in pupae forming on days 7 through 9. We expect most mosquitoes will emerge by day 9. Once these adults emerge from the pupae, they will be able to mate with and sterilize the wild mosquito population.

**Fig. 12:**
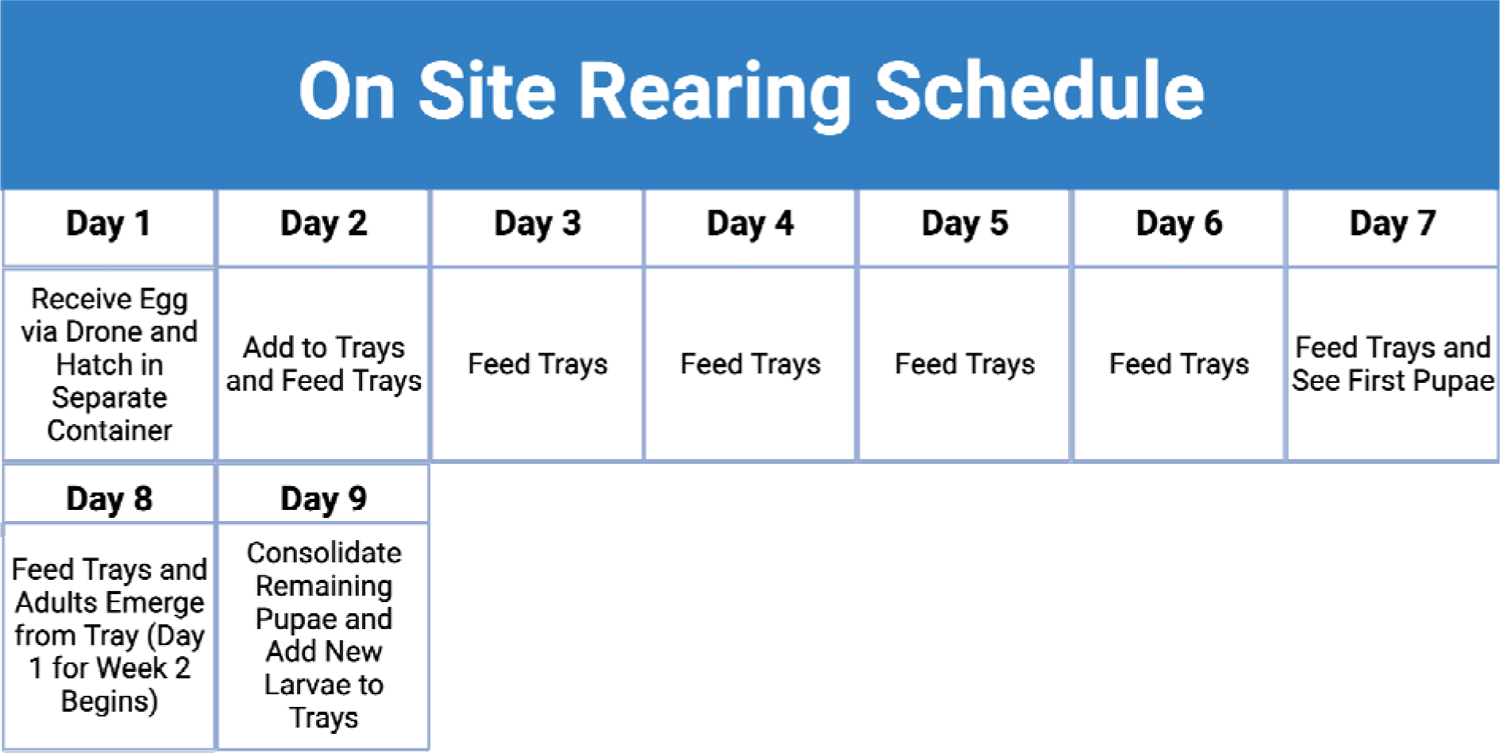
The Upper River on site rearing schedule.

Drones deliver eggs on Day 1, and they are hatched in a separate container. Larvae are fed following the IAEA recommendations, with pupae forming on days 7 through 9 post-hatch. Any pupae remaining on Day 9 are consolidated from the trays to make room for the next batch of larvae. If done in other rearing conditions, adjustments may be needed. Figure generated in BioRender.com.

##### 2.1.3.17 Entomological and epidemiological monitoring

Mosquito population and malaria incidence monitoring will be necessary to determine if the pgSIT sterile males are effective at reducing *A. gambiae* populations and malaria incidence. Two outcomes will determine the success of this intervention: localized suppression of wild *A. gambiae* (entomological impact) and reduction of malaria cases (epidemiological impact) in the treated area. Monitoring mosquito populations requires active efforts to capture/recapture mosquitoes to determine the abundance and composition of the mosquito populations. There are standardized methods for capturing mosquitoes through various trapping methods, i.e., swarm, and insecticide spray captures. Trapping or insecticide spray captures will likely be favored as the swarm captures may remove a significant number of the sterile males. Several sites will be selected throughout the region to account for the differences in landscape and relative location to other release sites. Regions near the center of the release coverage area are expected to have better population suppression due to reduced migration to these sites compared to border regions. Notably, pgSIT mosquitoes can be identifiable with a dye dusting method allowing rapid identification of wild and pgSIT mosquitoes via microscopy once the dead mosquitoes have been delivered to a laboratory (Yao et al. 2022). Monitoring efforts will require collaboration with the London School of Hygiene and Tropical Medicine (LSHTM), or similar organizations to create a new facility or to conduct this work at their field laboratory in Basse with the microscopes and other required equipment.

There are also standardized malaria monitoring programs within these regions and a network of clinics that provide malaria treatment (**Fig. 13**). These clinics can provide malaria incidence data for the region and baseline data to select villages for additional malaria monitoring needed to assess the epidemiological impact of current interventions compared to pgSIT intervention. There is also the option to employ more direct monitoring at the village level if we can allocate the funds for increased surveillance. Diagnostic tools can also be leveraged to better assess the asymptomatic case rate and gain a more in-depth assessment of malaria rates in these regions. An ideal approach may be conducting a cross-sectional survey at peak transmission. These diagnostic tools should be implemented prior to the first release of the pgSIT intervention and for several years after the release.

**Fig. 13:**
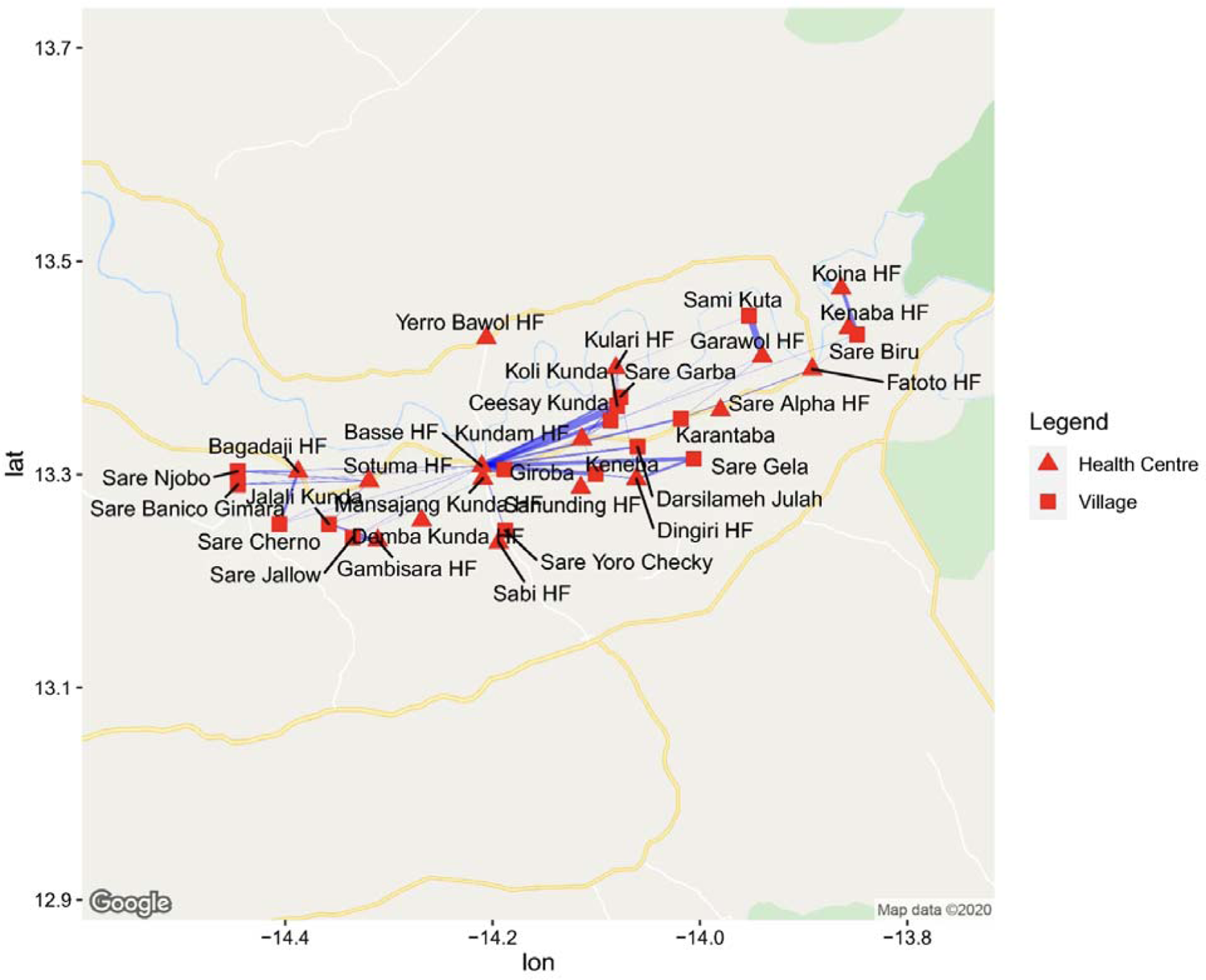
Map of the Upper River region in The Gambia showing the distribution of health facilities that provide malaria treatment and case monitoring.

This map was taken from (Broekhuizen et al. 2021) and shows the general distribution and density of healthcare centers (red triangle) in the Upper River region. The data collected at these facilities can be used to support the selection of sites for additional malaria monitoring and will provide information on the overall case rates in the region.

##### 2.1.3.17a Entomological and epidemiological monitoring costs

The monitoring of the *A. gambiae* population and malaria rates will likely require collaboration with The Medical Research Council (MRC) Unit of the LSHTM in Banjul, The Gambia. Research groups in this region could run the necessary studies to monitor the mosquito population and malaria incidence. To avoid year to year variation, we expect to require five years of monitoring once the facility is fully operational. The preliminary field trials will provide the data needed to set a baseline for the entomological studies. The monitoring costs are shown below. The experimental and operational budget should be fairly low because the monitoring will consist of inexpensive insect traps and communication with the existing medical network to determine the respective entomological and epidemiological impacts. This approach is the minimum required for monitoring, but can be scaled up to meet the project’s needs once baseline estimates are obtained from early field studies(**Table 15**).

Monitoring cost is dependent on the extent of monitoring. From our conversations with experts in the field, an extensive monitoring system could require up to 500,000 USD per year (per communication with Umberto D’Alessandro). We add this cost to the base value set in **Table 16**, and into the calculations in the **Table 27** monitoring section. However, smaller scale field trials should inform the scope and costs of monitoring required to assess safety, and entomological and epidemiological impact prior to large scale release.

**Table 16:** Monitoring costs for the first 5 years of mosquito releases.

| **Monitoring Staff**<br>**and Costs** | **Annual** | **Total** |
| --- | --- | --- |
| **Research Fellow** | 71,000 | 355,000 |
| **Technicians** | 13,500 | 202,500 |
| **Seasonal**<br>**Workers** | 9,000 | 45,000 |
| **Monitoring**<br>**Budget** | 30,000 | 150,000 |
| **Total** | 123,500 | 752,500 |

##### 2.1.3.18 Facility construction and supporting systems

The facility requires space to access the racks and cages, while providing room to move and clean this equipment. The total space for racks and cages was estimated, and the space was squared to account for additional space required for anterooms, and washing, sorting, maneuvering, and storage spaces required in the facility. By squaring the size estimate for the racks and cages, we aim to overestimate the space requirements for this facility (**Table 17)**.

**Table 17:** Facility size, land, and cost.

| **Conditions** | **Racks**<br>**Required** | **m^2 of**<br>**Space**<br>**for**<br>**Racks** | **Cages**<br>**Required** | **m^2 of**<br>**Space**<br>**for**<br>**Cages** | **Facility**<br>**Size** | **Land**<br>**Required**<br>**m^2** | **Facility**<br>**Cost**<br>**(USD)** | **Land Cost**<br>**(USD)** | **Architect**<br>**Fee**<br>**Estimate**<br>**USD** | **Total**<br>**(USD)** |
| --- | --- | --- | --- | --- | --- | --- | --- | --- | --- | --- |
| **COPAS Sorting,**<br>**High Fecundity,**<br>**High Survival** | 2 | 4 | 4 | 16 | 20 | 40 | 32,000 | 350 | 4,500 | 36,850 |
| **COPAS Sorting,**<br>**Low Fecundity,**<br>**High Survival** | 2 | 4 | 4 | 16 | 20 | 40 | 32,000 | 350 | 4,500 | 36,850 |
| **COPAS Sorting,**<br>**High Fecundity,**<br>**Low Survival** | 3 | 9 | 6 | 36 | 45 | 90 | 72,000 | 780 | 10,080 | 82,860 |
| **COPAS Sorting,**<br>**Low Fecundity, Low**<br>**Survival** | 3 | 9 | 6 | 36 | 45 | 90 | 72,000 | 780 | 10,080 | 82,860 |

**Table 18:**
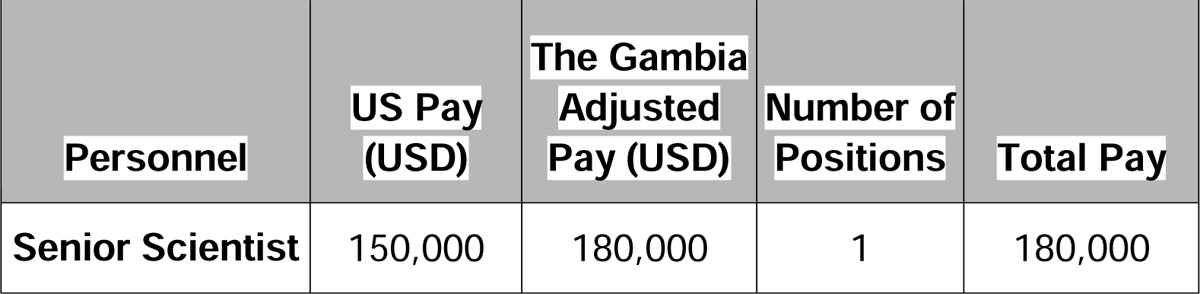
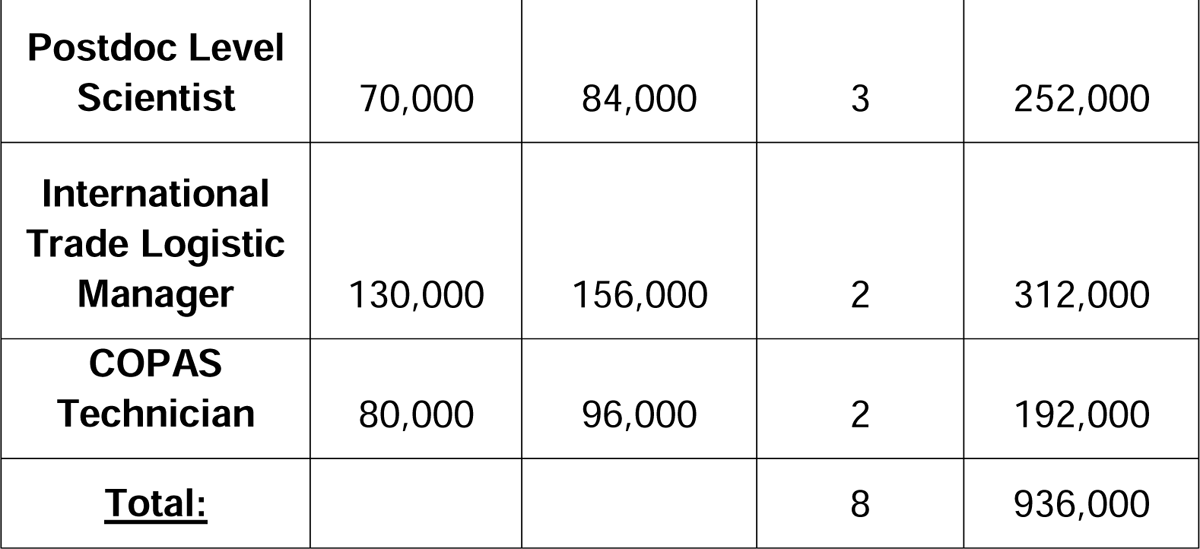
Initial personnel and training costs.

| **Personnel** | **US Pay (USD)** | **The Gambia Adjusted Pay (USD)** | **Number of Positions** | **Total Pay** |
| --- | --- | --- | --- | --- |
| **Senior Scientist** | 150,000 | 180,000 | 1 | 180,000 |
| **Postdoc Level Scientist** | 70,000 | 84,000 | 3 | 252,000 |
| **International Trade Logistic Manager** | 130,000 | 156,000 | 2 | 312,000 |
| **COPAS Technician** | 80,000 | 96,000 | 2 | 192,000 |
| **<u>Total:</u>** |  |  | 8 | 936,000 |

##### 2.1.3.19 Facility construction and supporting systems costs

To estimate the cost of land, we averaged the cost of available plots in Banjul and calculated the cost by square meter, which was 8.65 USD per square meter (**Table 17**). The size of the plot will vary depending on facility size, and we assume 50% more land space than the required facility space to have excess space to expand the facility.

This facility should have many of the structural requirements of a light-duty factory. Food processing plants fall into this category, and we were able to confirm from two sources that the construction costs of a light factory should be approximately 1,600 USD per square meter(Gerhard Brümmer, Lucy McLane, Elisa Campos, Thelma Hlatshwayo 2022; “The Cost of Building in Africa” 2022). With this in mind, the cost of the facility is estimated based on the size required to house the variable number of cages and racks. We also estimated the architectural design cost, which ranges from 7 to 14 percent of the facility size. The higher end 14 percent estimate was used to approximate the architectural design cost (**Table 17**). This preliminary estimate is based on limited data on the construction of such a facility in West Africa, so it may be safer to assume there are some unaccounted for costs associated with this construction. We hope to offset any missed costs by overestimating the size of the facility.

##### 2.1.3.20 Initial facility training

Initial facility establishment requires a team of experts in mosquito mass rearing to plan rearing schedules and operations, supervise the initial day-to-day activities, address issues, and train local staff to troubleshoot problems at the facility. The advisors would work at the facility for up to a year on-site, possibly broken into two separate work periods spread across two years or for a full year with the expectation of committing the maintenance phase to extensively train staff. This plan assumes that the facility begins Maintenance Phase production a couple of months before the expected Ramping Phase and Active Phase (**Section 2.1.4**) and that the professional staff stays a total of six months during the first season. The Ramping and Active Phase consist of five months total, which will be the most difficult time for the facility. If unexpected problems arise, they most likely will occur when the facility is working at 100% capacity. The goal is to have this support team there for a short time before the Ramping Phase, during the Active Phase, and for the return to the Maintenance Phase. Some staff may be required earlier to direct the facility purchasing and stocking or to provide logistical support to ensure the site has the necessary resources for the Ramping and Active Phases.

The professional training staff will have several roles to cover the initial project needs. They will include scientists with experience in mass rearing insects, preferably *A. gambiae*, logistic managers specializing in operations and developing the requisite systems, and technicians with expertise in sex sorting technology. Pay is adjusted according to the State Department’s hardship differential, which is 20%(“U.S. Department of State” n.d.).

First, several scientists with mosquito mass rearing experience should be hired to manage the mosquito racks, cages, egg laying, hatching, and various feeding methodologies. Specifically, at least one scientist with significant experience rearing *A. gambiae* should be included to support the establishment of the colonies and the unique needs of *A. gambiae*. Following colony establishment, a technician or postdoc level scientist could be hired to train local staff on the basics of mosquito mass rearing and rear the Maintenance Phase of mosquitoes. We recommend three technician/postdoc level scientists and one senior scientist to oversee the running of the facility. The average pay for these roles was identified from the LSHTM’s wages, so the adjusted pay scale may be an overestimate. The scientists will assist with training and troubleshooting production as it is scaled in the facility. It is likely ideal to hire both the technician and scientist to work on the ground for the first year of production and to train local staff to manage the insectary and troubleshoot issues. The goal for this first year is to ensure the local, permanent staff can troubleshoot future facility issues and will be able to reach out to experts abroad if further issues arise.

The second group is logistics managers, who can work to develop efficient supply chains and sourcing of resources and manage the complexities of running a large production facility. Ideally, someone with experience working in West Africa with involvement in similar scale agricultural or light manufacturing work. These individuals have the requisite skill set to assist in developing this facility. The range of a logistic manager can range from 80,000 to 150,000 USD depending on experience level. As with the scientist role, if the logistic manager is not local, they will aim to recruit and train local managers to fulfill this role in the long term.

Finally, we require staff with expertise in the selected sex sorting technology. They need to be capable of training local staff and maintaining and addressing any issues that arise with this technology. The average pay for a flow cytometer expert, which is a scaled-down version of a COPAS, is approximately 66,000 USD annually. However, the COPAS FP 500 has fewer technicians explicitly trained on this machine available, likely demanding greater pay. If we can only recruit flow cytometry experts, they would have to undergo training on the COPAS before beginning their year in The Gambia, which would result in similar wages. This technology is the backbone of the facility and is a pivotal component for this facility to function. The importance of this technology makes it a priority to ensure good stewardship of the machines. Senecio Robotics or even Verily may become the supplier of the requisite sex sorting technology. If this occurs, a similar expert will be required, and we would expect similar salaries (**Table 21**).

##### 2.1.3.21 Facility annual labor

The initial labor force is to be eventually fully replaced by citizens of The Gambia to improve community investment in the project, reduce the costs of labor, and create a financial investment into the community. We offer three potential wage estimates spanning the range of expected wages needed to attract labor of sufficient skill. Labor costs may be further reduced as this work would be primarily seasonal, although we are working under the assumption that maintaining a workforce year round would greatly benefit the facility’s stability and training. This work force could be utilized to produce mosquito products abroad to fund this project as mentioned later in the discussion section. These costs were estimated by looking at comparable rates from our collaborators in The Gambia (**Tables 19-21**).

**Table 19:**
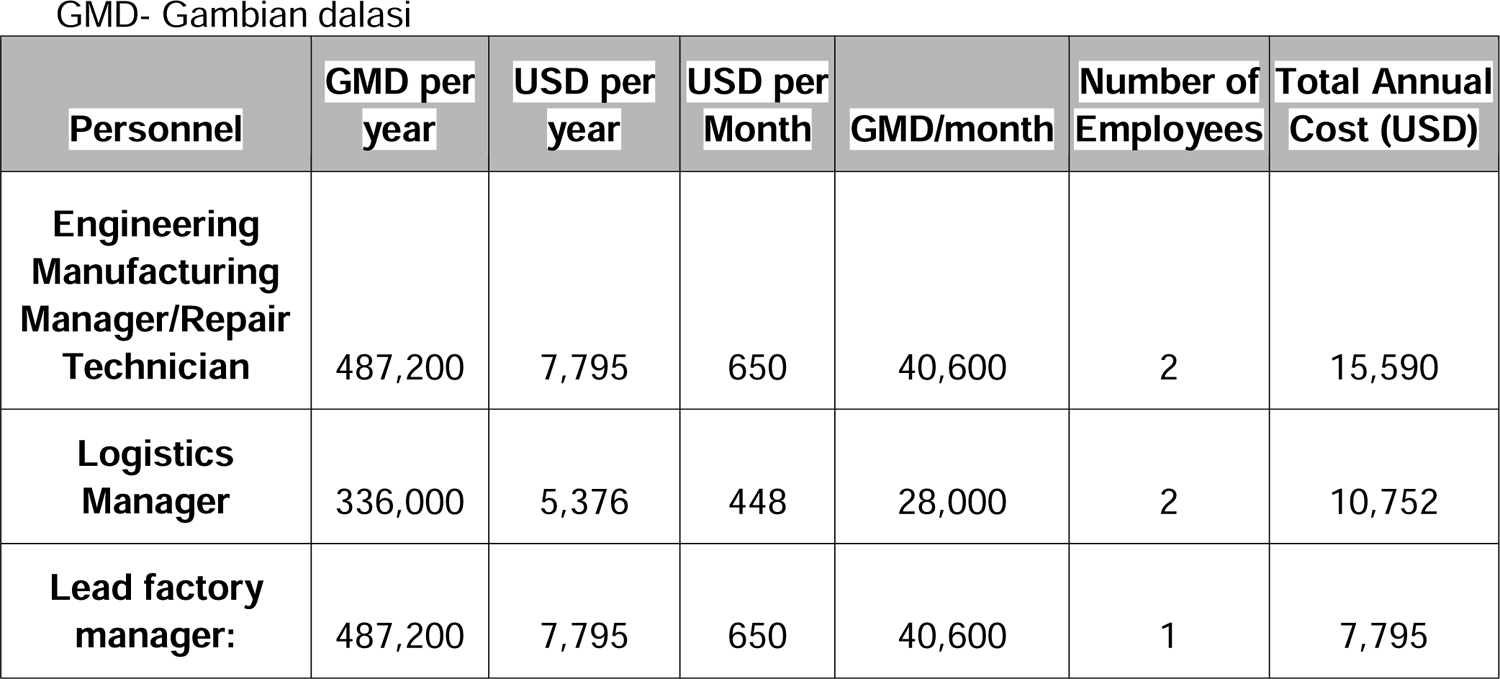
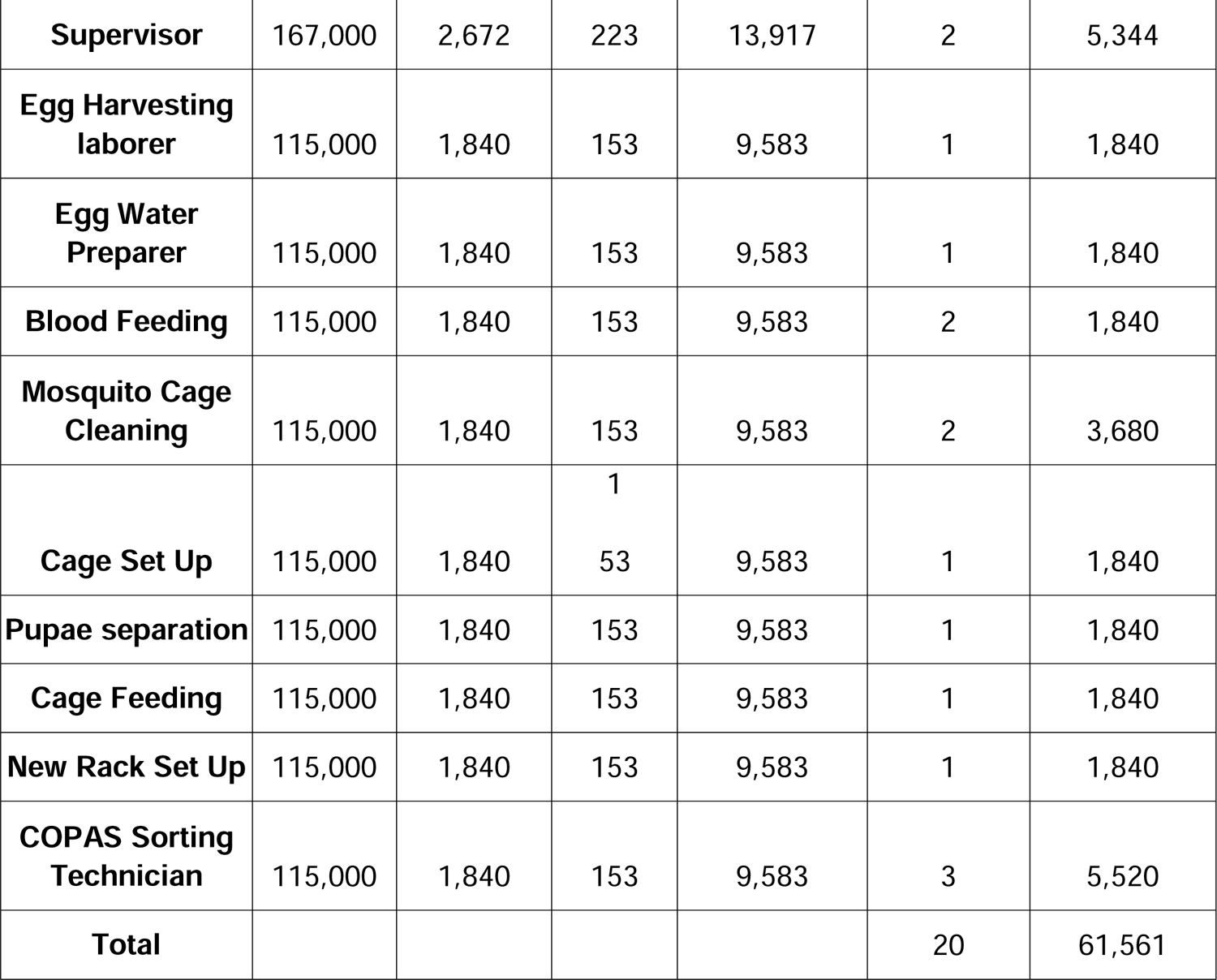
High wage annual estimate.

**Table 20:**
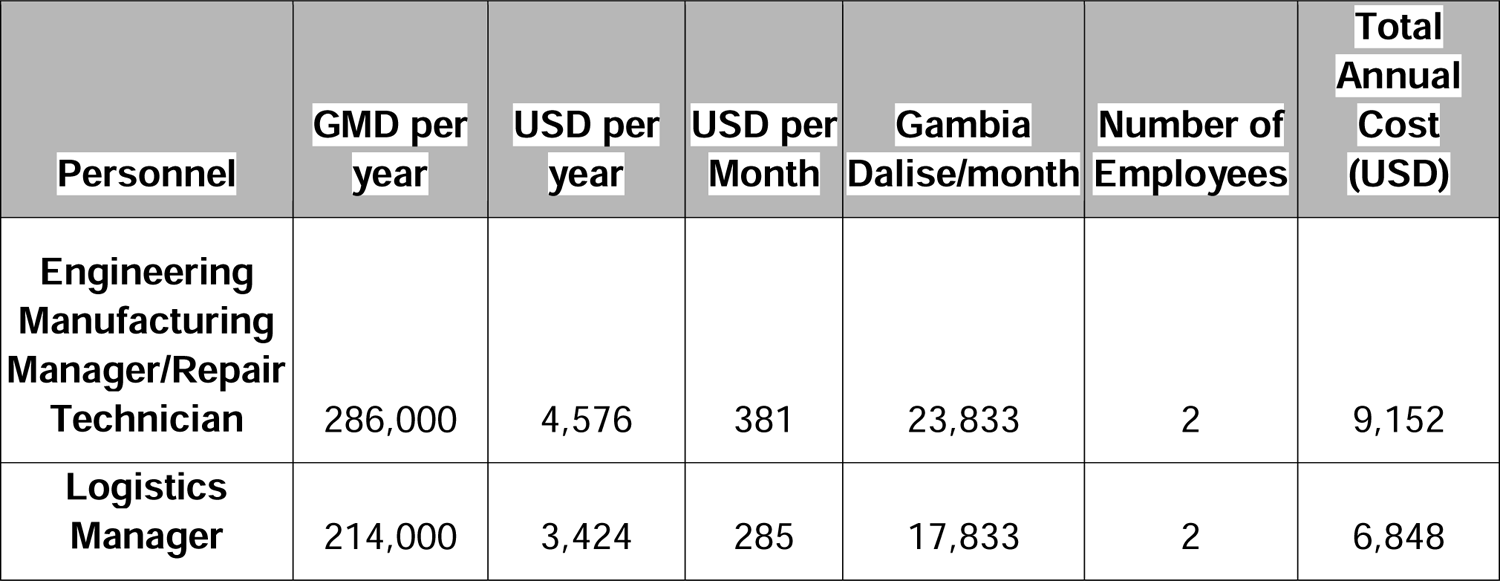
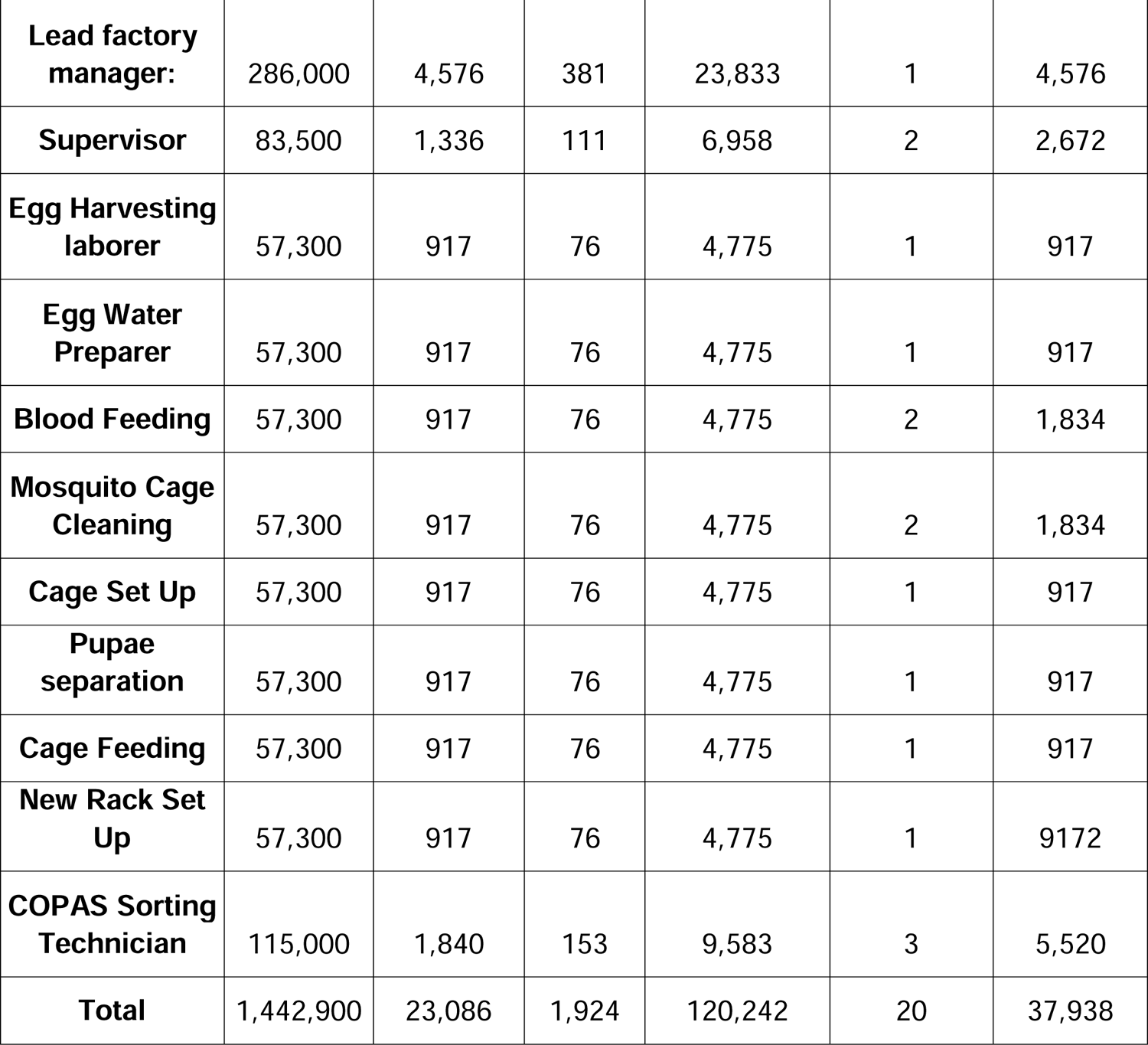
Medium wage annual estimate.

| **Personnel** | **GMD per year** | **USD per year** | **USD per Month** | **Gambia Dalise/month** | **Number of Employees** | **Total Annual Cost (USD)** |
| --- | --- | --- | --- | --- | --- | --- |
| **Engineering Manufacturing Manager/Repair Technician** | 286,000 | 4,576 | 381 | 23,833 | 2 | 9,152 |
| **Logistics Manager** | 214,000 | 3,424 | 285 | 17,833 | 2 | 6,848 |
| **Lead factory manager:** | 286,000 | 4,576 | 381 | 23,833 | 1 | 4,576 |
| **Supervisor** | 83,500 | 1,336 | 111 | 6,958 | 2 | 2,672 |
| **Egg Harvesting laborer** | 57,300 | 917 | 76 | 4,775 | 1 | 917 |
| **Egg Water Preparer** | 57,300 | 917 | 76 | 4,775 | 1 | 917 |
| **Blood Feeding** | 57,300 | 917 | 76 | 4,775 | 2 | 1,834 |
| **Mosquito Cage Cleaning** | 57,300 | 917 | 76 | 4,775 | 2 | 1,834 |
| **Cage Set Up** | 57,300 | 917 | 76 | 4,775 | 1 | 917 |
| **Pupae separation** | 57,300 | 917 | 76 | 4,775 | 1 | 917 |
| **Cage Feeding** | 57,300 | 917 | 76 | 4,775 | 1 | 917 |
| **New Rack Set Up** | 57,300 | 917 | 76 | 4,775 | 1 | 9172 |
| **COPAS Sorting Technician** | 115,000 | 1,840 | 153 | 9,583 | 3 | 5,520 |
| **Total** | 1,442,900 | 23,086 | 1,924 | 120,242 | 20 | 37,938 |

**Table 21:** Low wage annual estimate.

| **Personnel** | **GMD per year** | **USD per year** | **USD per Month** | **Gambia Dalise/month** | **Number of Employees** | **Total Annual Cost (USD)** |
| --- | --- | --- | --- | --- | --- | --- |
| **Engineering Manufacturing Manager/Repair Technician** | 214,500 | 3,432 | 286 | 17,875 | 2 | 6,864 |
| **Logistics Manager** | 160,500 | 2,568 | 214 | 13,375 | 2 | 5,136 |
| **Lead factory manager:** | 214,500 | 3,432 | 286 | 17,875 | 1 | 3,432 |
| **Supervisor** | 62,625 | 1,002 | 84 | 5,219 | 2 | 2,004 |
| **Egg Harvesting laborer** | 42,975 | 688 | 57 | 3,581 | 1 | 688 |
| **Egg Water Preparer** | 42,975 | 688 | 57 | 3,581 | 1 | 688 |
| **Blood Feeding** | 42,975 | 688 | 57 | 3,581 | 2 | 1,376 |
| **Mosquito Cage Cleaning** | 42,975 | 688 | 57 | 3,581 | 2 | 1,376 |
| **Cage Set Up** | 42,975 | 688 | 57 | 3,581 | 1 | 688 |
| **Pupae separation** | 42,975 | 688 | 57 | 3,581 | 1 | 688 |
| **Cage Feeding** | 42,975 | 688 | 57 | 3,581 | 1 | 688 |
| **New Rack Set Up** | 42,975 | 688 | 57 | 3,581 | 1 | 688 |
| **COPAS Sorting Technician** | 86,250 | 1,380 | 115 | 7,188 | 3 | 4,140 |
| **Total** | 1,082,175 | 17,315 | 1,443 | 90,181 | 20 | 28,456 |

##### 2.1.3.22 Testing and field trials costs

Prior to production, the *A. gambiae* pgSIT has to (1) integrate sex specific markers into the pgSIT 1.0 lines to facilitate COPAS sex sorting technologies (pgSIT 2.0 with fluorescent sex specific markers), (2) evaluate pgSIT 2.0 in small scale field testing and (3) obtain regulatory approvals for small and large scale field testing. The currently developed pgSIT system (pgSIT 1.0) in our lab shows 100% female-killing and 100% male sterilization among many thousands of insects tested, demonstrating perfect proof-of-principle of the technology in the species (**Fig. 1**). The manuscript is in the final stages of preparation and is nearing submission. However, these pgSIT 1.0 lines lack a sex-selection marker making optical sorting the only available sorting technology amenable to this system. In the absence of a collaboration with Verily or more precise sorting speed estimates from Senecio Robotics, in addition to the increased costs associated with rearing individuals to adulthood, these are not feasible options for pgSIT mass release. Therefore, this analysis focuses on releasing the pgSIT 2.0 technology, which integrates genetic sex sorting with male sterilization.

##### 2.1.3.22a Development of pgSIT 2.0 to support COPAS sex sorting

The pgSIT technology has been developed in *A. gambiae* has been fully developed (**Fig. 1**), but these lines need to be integrated with sex-specific fluorescent markers to support the COPAS sex sorting technology, which requires these markers to separate the sexes - a critical step for establishing the parental crosses. The pgSIT 2.0 lines that contain these sex-specific markers are currently under development.

The primary cost to update this line is labor and resources for this system to be created and to confirm the efficacy of this line using standard laboratory practices. This work is currently underway; the approximate costs to cover this work correspond to the cost of one postdoctoral scholar, two or three dedicated paid research assistants, research supplies, and overhead. It may take one to two years to complete the creation and assessment of these lines.

##### 2.1.3.22b Field testing and small scale release overview

Prior to large-scale release in The Gambia, field testing and small scale releases will be required to obtain safety and efficacy data, regulatory approval and community support for large scale release. These small scale trials also allow testing and optimization of the mass rearing technologies. There are three phases in small scale testing: (1) initial cage trials and preparation (Phase II), (2) small scale releases, and (3) functional scale release (Phase III, **Fig. 4**). Not only will the technology be assessed for safety and efficacy, but the mass rearing aspects of this technology will be tested, and stakeholders and the community will be involved in this development process.

##### 2.1.3.22c Initial cage trials and small scale release preparation

Cage trials in The Gambia will evaluate the pgSIT technology in the local environment and genetic background by introgressing the pgSIT into local strains. These studies will provide the efficacy data required for small and large scale releases, community engagement opportunities to gain input and buy-in for field releases, and will evaluate and optimize rearing procedures and technologies required to scale up egg production for field releases. These initial experiments will require small scale facilities and are dependent on procuring necessary permits.

The preparation for the small-scale release will require engagement with local communities, The Gambian government, and other stakeholders to obtain their feedback and approval to advance the project to field release. We plan to incorporate many of the core commitments we developed for our gene drive technologies, such as commitments towards fair partnership and transparency in the evaluation and development of the pgSIT technology and field trials, product efficacy and safety, regulatory evaluation, risk and benefit assessment, trial monitoring, and mitigation (Long et al. 2020) to strive for transparency and inclusion of stakeholder perspectives in the development of our trials. We also benefit from the experience of other groups bringing other mosquito technologies to the field (Schairer et al. 2021) and from the experience of our partners at the MRC Unit working with the communities in the URR. Ultimately, these cage trials will provide the data needed to support small field releases and is an important step to obtain feedback from the local community and other stakeholders.

The most cost effective approach to the cage trial is to contract this work to the LSHTM, which has extensive expertise in *A. gambiae* rearing and has led multiple field trials across West Africa. It is likely possible to do this in the field laboratory in Basse, The Gambia. Otherwise, there may be additional infrastructure and equipment costs in a less established area. A rough estimate would be to assume the need for one postdoctoral scholar or research fellow to lead this project, laboratory technicians, and resources to complete these experiments. A research fellow at the LSHTM in The Gambia is paid at most 71,000 USD annually, and local lab technicians are paid approximately 4,500 USD annually. One fellow and three technicians are required to be well staffed. This project would take two to three years to complete introgression and cage trials. Introgression is the process of crossing GE lines to the local mosquito population to improve reproductive compatibility and line fitness upon release (**Table 22**).

**Table 22:**
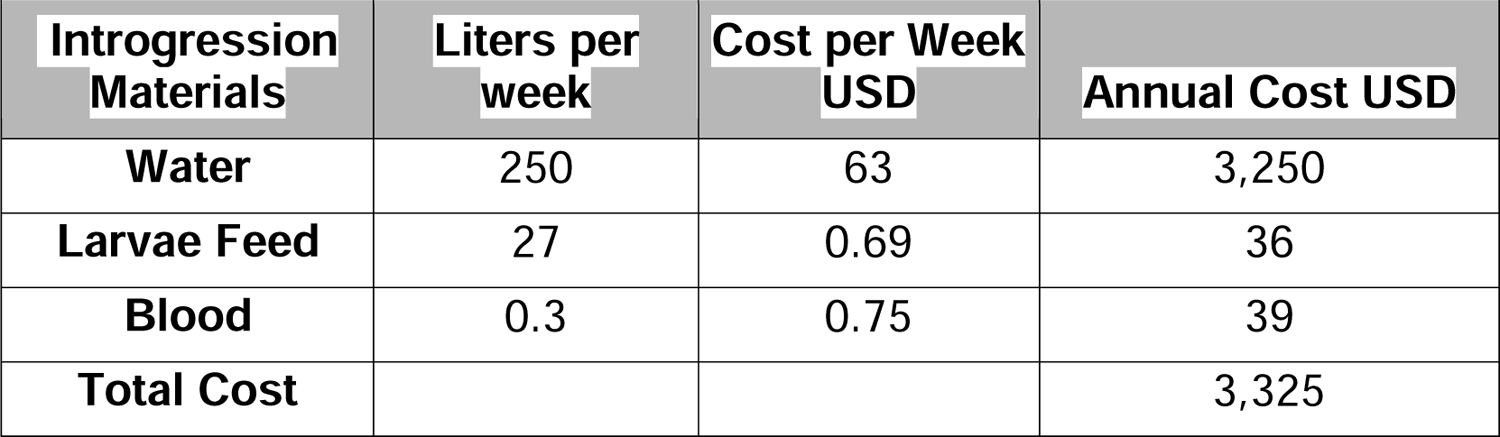
Annual cost of introgression experiments.

During this stage, minimal equipment is needed for mass rearing. These pieces of equipment are one to two Wolbaki mass rearing rack and tray systems and at least four large cages. As stated earlier, the Wolbaki mass rearing rack and tray systems are 22,500 USD each. Cages used for cage trials vary across papers, but it is possible that the mass rearing adult cages could be utilized for the field trials, and these cost 250 USD each. Four of these cages can be purchased as they will be needed for future studies. There may be an additional cost of a fluorescent microscope to screen the mosquitoes, which can range in price from 2,000 to 20,000, but a microscope at the 10,000 price point will likely suffice. The cost of the raw materials would likely consist of mosquito feed, water, large cages for outdoor cage rearing, basic laboratory equipment such as thermocyclers, and sequencing costs for confirming species introgression. Considering that this facility will be rearing up to two racks of mosquitoes year round, we expect at most 3,325 USD annually in food and water costs. However, 100,000 USD in funding for troubleshooting is recommended at this stage because it can be used to solve problems that may be encountered in field trials. Sequencing and other assays associated with introgressing mosquitoes into the local genetic background is expected to be 100,000 USD a year to run necessary experiments. In total, these costs will be approximately 1,150,475 USD. This cost is a conservative estimate to offset unforeseen costs (**Table 23**).

**Table 23:** Total budget estimate of introgression and initial cage trials.

| **Cost Category** | **Annual Cost** | **Total** |
| --- | --- | --- |
| **Facility Rental** | 100,000 | 300,000 |
| **Research Fellow** | 71,000 | 213,000 |
| **Technicians** | 13,500 | 121,500 |
| **Cages** | 250 | 1,000 |
| **Wolbaki Mass Rearing Racks** | 22,500 | 45,000 |
| **Fluorescent Microscope** | 10,000 | 10,000 |
| **Experiment Budget** | 150,000 | 300,000 |
| **Mosquito Rearing Input** | 3,325 | 9,975 |
| **Total** | 370,575 | 1,150,475 |

##### 2.1.3.22d Small scale field trials

Pending the necessary approvals and regulatory guidance, the likely next step is a small scale trial to confirm the efficacy and safety of pgSIT sterile males in the field (Phase III, **Fig. 4**). These studies will assess pgSIT mosquito movement, distribution, longevity, and any unexpected introgression of pgSIT genes into the local mosquito population. These studies will include preliminary research of the entomological impact of the technology, but this phase will focus on the safety and risk assessment information required for large scale studies, which can be easily acquired with minimal releases.

Small scale releases are also useful for the initial testing and optimization of mass scale rearing. We can evaluate the proposed equipment and procedures, for example, to obtain better survival and fecundity rate estimates, which will allow us to more accurately estimate the upfront and annual facility costs and the resources required for scaling egg production for mass release. The egg production needs of a small scale study may be too low to necessitate sex sorting technologies, but we may be able to demo these systems for a limited time, or begin optimizing the sex sorting technologies in the larger scale release phase.

When considering the costs of the small scale field trial, it is important to note that costs can build off of equipment, resources, knowledge, and expertise gained during the cage trial. The primary changes to the budget will be resources for release monitoring t and potentially the purchase of sex sorting devices. The small scale field trial can utilize the mass rearing racks and cages and should take one to two years to accomplish. There may be some initial monitoring experiments the year before the small scale release to gauge the native mosquito population abundance and distribution. Pre-release monitoring can also facilitate site selection and identify controls for the pilot study. Purchasing a single COPAS FP 500 first may be the best approach, whereas Senecio Robotics systems are custom and may be trickier to acquire and later scale up individual systems. There may be options to purchase a smaller version of Senecio Robotics that could be scaled up. Either way, further testing and collaboration with both sex sorting companies would be appropriate to confirm efficacy before purchase (**Table 24**).

**Table 24:** Total budget estimate of small scale trial.

| **Cost Category** | **Annual Cost** | **Total** |
| --- | --- | --- |
| **Facility Rental** | 100,000 | 200,000 |
| **Research Fellow** | 71,000 | 142,000 |
| **Technicians** | 13,500 | 81,000 |
| **COPAS FP 500** | 308,000 | 344,960 |
| **Part time Workers**<br>**for Release Season** | 4,500 | 9,000 |
| **Experiment Budget** | 100,000 | 200,000 |
| **Mosquito Rearing**<br>**Input** | 3,325 | 6,650 |
| **Total** | 600,325 | 983,610 |

**Table 25:** Total budget estimate for large scale field trials.

| **Cost Category** | **Annual Cost** | **Total** |
| --- | --- | --- |
| **Facility Rental** | 100,000 | 200,000 |
| **Research Fellow** | 71,000 | 142,000 |
| **Technicians** | 13,500 | 81,000 |
| **COPAS FP 500** | 36,960 | 73,920 |
| **Part time Workers for Release Season** | 9,000 | 18,000 |
| **Experiment and Production Budget** | 100,000 | 200,000 |
| **Mosquito Rearing Input** | 3,325 | 6,650 |
| **Total** | 333,785 | 667,570 |

**Table 26:**
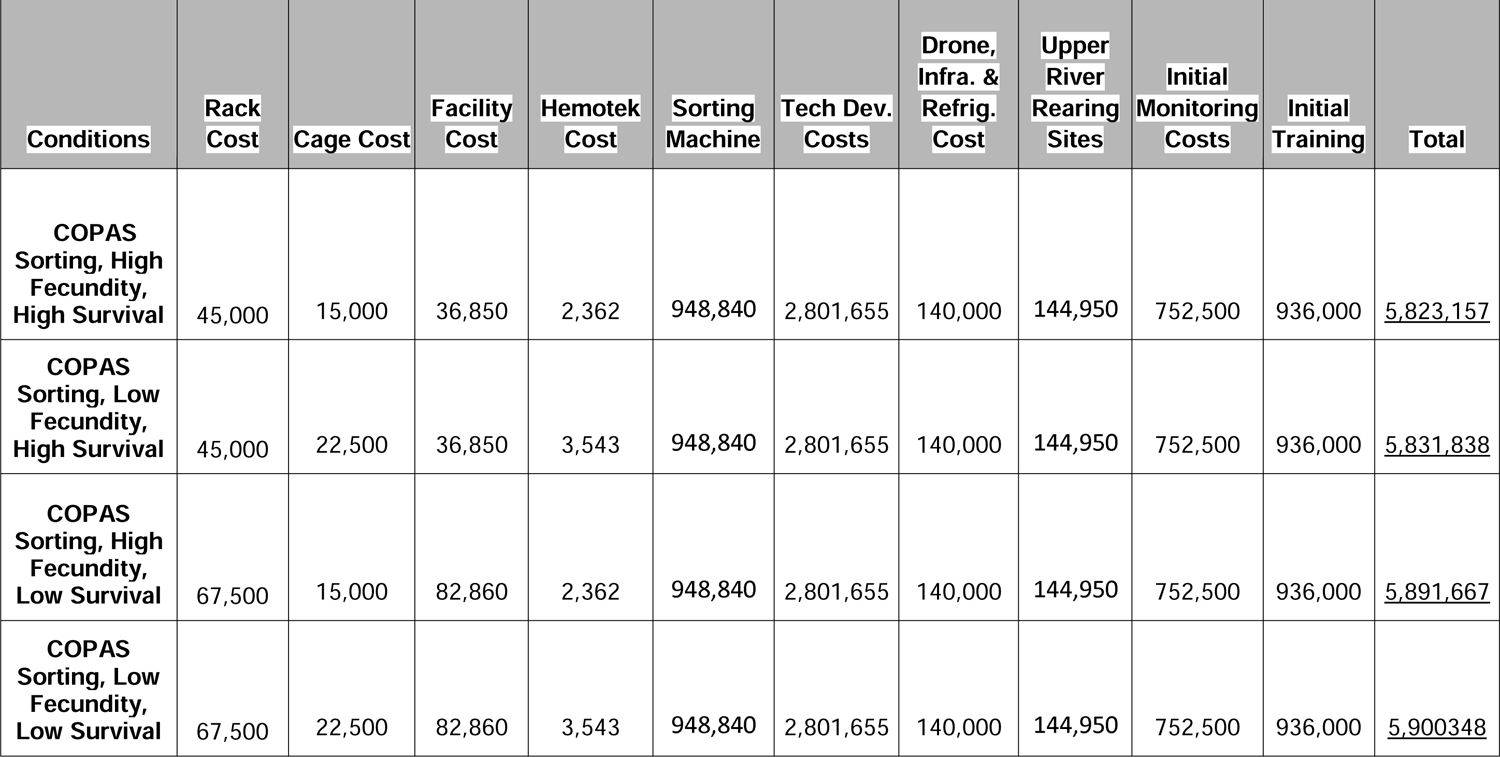
Total initial costs with least expensive trials: This cost assessment is based on the direct cost estimates of trials based on assumed reagent requirements, wages for local employees in The Gambia and other expected expenses. A high cost can be seen in Table 27.

**Table 27:** Total initial costs with more expensive trials.

| Initial Cost | Rack Cost | Cage Cost | Facility Cost | Hemotek Cost | Sorting Machine | Development Costs | Drone, Generator & Refrigerator Cost | Upper River Rearing Sites | Initial Monitoring Costs | Initial Training | Total |
| --- | --- | --- | --- | --- | --- | --- | --- | --- | --- | --- | --- |
| COPAS Sorting, High Fecundity, High Survival | 45,000 | 15,000 | 36,850 | 2,362 | 948,840 | 5,801,655 | 140,000 | 144,950 | 3,252,500 | 936,000 | <u>11,323,157</u> |
| COPAS Sorting, Low Fecundity, High Survival | 45,000 | 22,500 | 36,850 | 3,543 | 948,840 | 5,801,655 | 140,000 | 144,950 | 3,252,500 | 936,000 | <u>11,331,838</u> |
| COPAS Sorting, High Fecundity, Low Survival | 67,500 | 15,000 | 82,860 | 2,362 | 948,840 | 5,801,655 | 140,000 | 144,950 | 3,252,500 | 936,000 | <u>11,391,667</u> |
| **COPAS**<br>**Sorting, Low**<br>**Fecundity,**<br>**Low Survival** | 67,500 | 22,500 | 82,860 | 3,543 | 948,840 | 5,801,655 | 140,000 | 144,950 | 3,252,500 | 936,000 | <u>11,400,348</u> |

##### 2.1.3.22e Large scale field trials

Large scale releases can provide additional safety and essential efficacy data to evaluate pgSIT sterile male suppression of mosquito populations. During the cage studies and small scale releases, we will have built relationships with the community and other stakeholders, and accumulated data from our studies, local health centers, and vector management organizations, to identify multiple areas for larger-scale trials. Community members will also become more involved in the rearing of sterile male mosquitoes from eggs to adulthood within their village.

The scale of these studies will be informed by the data collected during the cage and small field trials as well as data and input from local stakeholders. The scale of these studies will increase and will require moderate scaling of egg production. A village in the URR, for example, may have a population of approximately 1,000 people, so the mosquito sorting requirements are no longer manageable manually, as about 58,000 larvae will need to be sorted daily. During this phase, a COPAS FP 500 or another sorting technology will need to be implemented to support egg production. This approach will allow us to test and optimize sex sorting technologies in the field. In particular, these sorting devices should be pushed to their limits to determine their capabilities and stress-test their hardware.

The cost of the large scale field trial can utilize any devices purchased during the small scale field trial. The primary costs are the same as in previous trials, but with an increase in the experiment budget for monitoring (**Table 25**). Additional fees will be needed to maintain the COPAS FP 500, and the monitoring and releases should continue for two years, which is accounted for in the budget. We also added some seasonal workers to support the mosquito field collections. These individuals would only be hired for a month or two per year.

Depending on the results of the previous studies and technical needs, a larger scale cluster randomized trial similar to the recent trial at Dabira could also be implemented (Dabira et al. 2022). A large scale cluster randomized trial in the region cost approximately 2.5 million USD (As per communication with Umberto D’Alessandro). This cost is included in **Table 26** and would increase the initial cost of development but would not have long term effects on the annual costs. There are outstanding questions as to the scale and design of the field trials, but as the pgSIT technology advances in development, we will acquire the data to address these questions.

##### 2.1.3.22f Regulatory approvals for large scale deployment

Throughout the field trials and the transition to large scale use, we will engage regulatory stakeholders to ensure compliance with the relevant laws and regulations. The Gambia has yet to deploy GE mosquitoes, and has not yet established laws and governance for such activities. The Gambia is expected to follow guidelines set out by the Cartagena Protocol on Biosafety. Organizations such as Target Malaria in other African regions may have paved the road for future GE mosquito releases on the continent. Still, there is also the possibility that our project may prompt the creation of a country-specific governance or a regulatory body in The Gambia. No matter the approval process, frequent engagement with regulators early and throughout this process should establish the regulatory pathway for the pgSIT technology. Other GE mosquito releases have occurred elsewhere in Africa and the rest of the world, so there is a precedent for these releases. Once the regulatory requirements are better defined, additional costs may be required to support regulatory compliance activities. Because the development of regulations can take time, we plan to engage the Gambian government early in the project planning to identify the best path toward regulatory approval.

##### 2.1.3.23 Summary of pgSIT facility costs

The initial construction and equipment purchase costs for the facility for this project range from approximately 6 to 11.5 million USD (**Tables 26 and 27**). These costs include the construction and setup of the necessary infrastructure, such as equipment, sex sorting devices, initial labor investment of scientists and experts, and the mass rearing facilities. The wide range in costs is primarily due to the uncertainties associated with key factors, such as fecundity, larval survival rates, and the implementation of sex sorting technology. These uncertainties make it challenging to determine the exact financial requirements for the project. **Table 26** summarizes these inputs. One important note is that the field trials would have already procured a COPAS FP 500, so this cost is subtracted from the COPAS sex sorting costs. However, as field work often results in wear-and-tear on the equipment, replacements may be required in the future. Extra sex sorting machines and more costly field and monitoring trials are accounted for in **Table 27**.

In addition to the upfront facility costs, there are also annual costs associated with the operation and maintenance of the facility. These costs are about 315,000 USD per year (**Table 28**). The annual expenses cover various aspects, including staff salaries, ongoing maintenance, and repair, purchasing feed and supplies, as well as funding to support release sites in the URR (**Table 28**). For simplicity, only the high salary estimates were used in the calculations, but this can likely be reduced. Additionally, there are estimates to double the number of sex sorting machines if we need to offset additional maintenance time for these machines (**Table 29**).

**Table 28:**
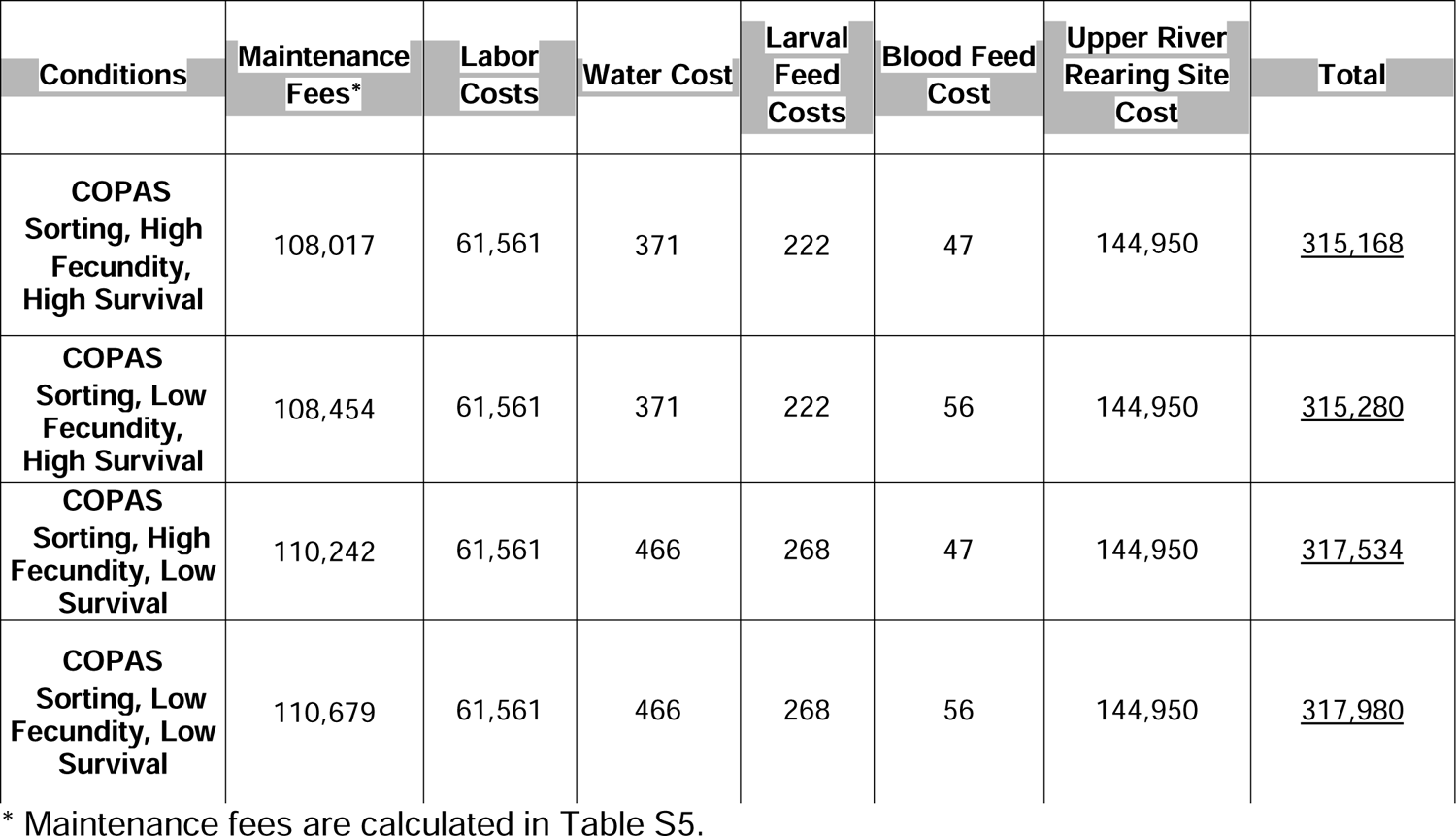
Total annual costs: The total annual cost includes facility costs, including maintenance fees for the equipment and the high estimated cost in The Gambia, for labor, local resources and rearing in the URR. Rearing costs may be overestimated, however, as other malaria interventions rely on some volunteer labor. Maintenance of larvae at sites in the URR may also utilize other more affordable, local resources rather than imported mosquito feed.

The broad range in both upfront and annual costs highlights the need for further testing and evaluation of the pgSIT technology at smaller scales. By conducting smaller scale trials and experiments, we can gather more accurate data on the fecundity of the organisms, survival rates of larvae, and the effectiveness of sex sorting technology using the exact equipment used at scale. These findings will provide valuable insights and help refine the estimates for the project’s costs. Additionally, testing at smaller scales allows for the identification of potential challenges. It provides opportunities to develop strategies to mitigate risks before scaling up the operation, ensuring a more cost-effective and efficient implementation of the project in the long run.

This table includes additional costs associated with the larger and more expensive field trials and additional machines for sex sorting. The additional field trial cost are: 1) 500,000 USD per year for the five year monitoring period, 2) an additional 2.5 million USD for the larger scale cluster randomized trial (approximate cost of a previous large scale trial(Dabira et al. 2022) as per communication with Umberto D’Alessandro) and 3) 500,000 USD to pay for a social science team to manage public communication of pgSIT technology. As a facility of this scale has not been developed, and extensive testing has not yet been conducted with these machines, this may need to be tested. These estimates should represent the upper limit of initial costs, but there are significant uncertainties on whether these expenditures will be necessary.

#### 2.1.4 Mosquito rearing facility phases and labor costs

pgSIT mosquitoes do not need to be constantly released throughout the year to reduce mosquito abundance and malaria transmission, since *A. gambiae* in the URR and many other regions throughout their range only have seasonally elevated populations (in The Gambia this is the rainy season beginning June 1^st^-late October). The rest of the year, the *A. gambiae* population size is too low to sustain malaria transmission. To minimize the cost of rearing mosquitoes for a large-scale facility, the facility’s year can be divided into 3 phases for consideration: Maintenance Phase (34 weeks), Ramping Phase (6 weeks), and Active Phase (12 weeks). The Maintenance Phase is designed to keep a minimal population of the parent Cas9 and gRNA lines alive and reproducing to minimize labor. The Ramping Phase is used to rapidly increase the number of mosquitoes to the numbers required to meet the egg production needs in the Active Phase while spreading the daily egg batches across three weeks to maximize the efficiency of the facility. The release schedule is based on weekly releases of eggs to parts of The Upper River. Functionally, this production will be spread throughout the week rather than shipping the eggs once a week. The Active Phase is the full sterile egg production phase for the delivery of sterile male pgSIT mosquitoes to the field sites. During the active phase, the facility is running at near maximum capability with contingencies to account for equipment failures or distribution problems. Upon completion of the Active phase, which is at the end of the mosquito season, mosquitoes are culled down to Maintenance Phase numbers.

In order to time the releases with the beginning of the rainy season, the Active Phase egg hatching should start on May 16th, two weeks before June 1st. The Active Phase will continue to produce daily batches until 14 weeks later, on August 22nd. This last batch can be culled to Maintenance level production on approximately September 12th. There may be a need to stagger egg production by a couple of days to align the Maintenance Phase schedule to the Ramping Phase Schedule. The first Ramping Phase would need to occur nine weeks before the start of the Active Phase. March 14th will be the approximate start of the last Maintenance Phase, where there will be three egg harvesting cycles to begin the shift to the Ramping Phase. To prepare for the beginning of the Active Phase on May 16th, April 4th is the approximate date of the beginning of the Ramping Phase. This cycle will repeat annually.

##### 2.1.4.1 Maintenance phase

The Maintenance Phase was designed to minimize cost while sustaining a large, stable population for expansion during the Ramping Phase. This option has been outlined for larger scale production or to further reduce maintenance phase work. Synchronization of the mosquito life cycle in the Maintenance Phase will necessitate a Ramping Phase to spread production over a week and minimize the burden on COPAS sex sorting and other activities. We aimed to (1) minimize the work hours required to manage a large-scale facility and (2) minimize resources used to sustain the mosquito population during this phase.

The Maintenance Phase achieves minimal costs by maintaining small batches of mosquitoes in a 3-week synchronized schedule (**Fig. 14**). The Active Phase, where the production is at its highest, has life cycles beginning every day throughout this period, but the Maintenance Phase is designed to minimize labor and resources when sterile male egg production is not needed. So, production is started once every three weeks instead. While the rearing phases have similarities, the Maintenance Phase prioritizes minimizing costs and, therefore, only harvests eggs once during the life cycle of the mosquitoes. Specifically, during the Active phase, multiple generations of mosquito are ongoing concurrently, in parallel, each staggered by a day. In the Maintenance phase, only one generation is ongoing at a time. This phase will still require two blood meals, but this is only to increase longevity and maintain the regularity of the cycle. There is the potential to harvest eggs during the first blood feeding shown in **Fig. 14** and refrigerate the eggs for up to five days to maintain the same schedule, but viability testing would be required prior to implementation. Our experts do not recommend this, however, so we selected the slightly more expensive double blood feed strategy. During the Maintenance Phase, the goal is to extend the egg harvesting to the end of the third week to maintain a regular cycle, which minimizes the cost of this approach.

**Fig. 14:**
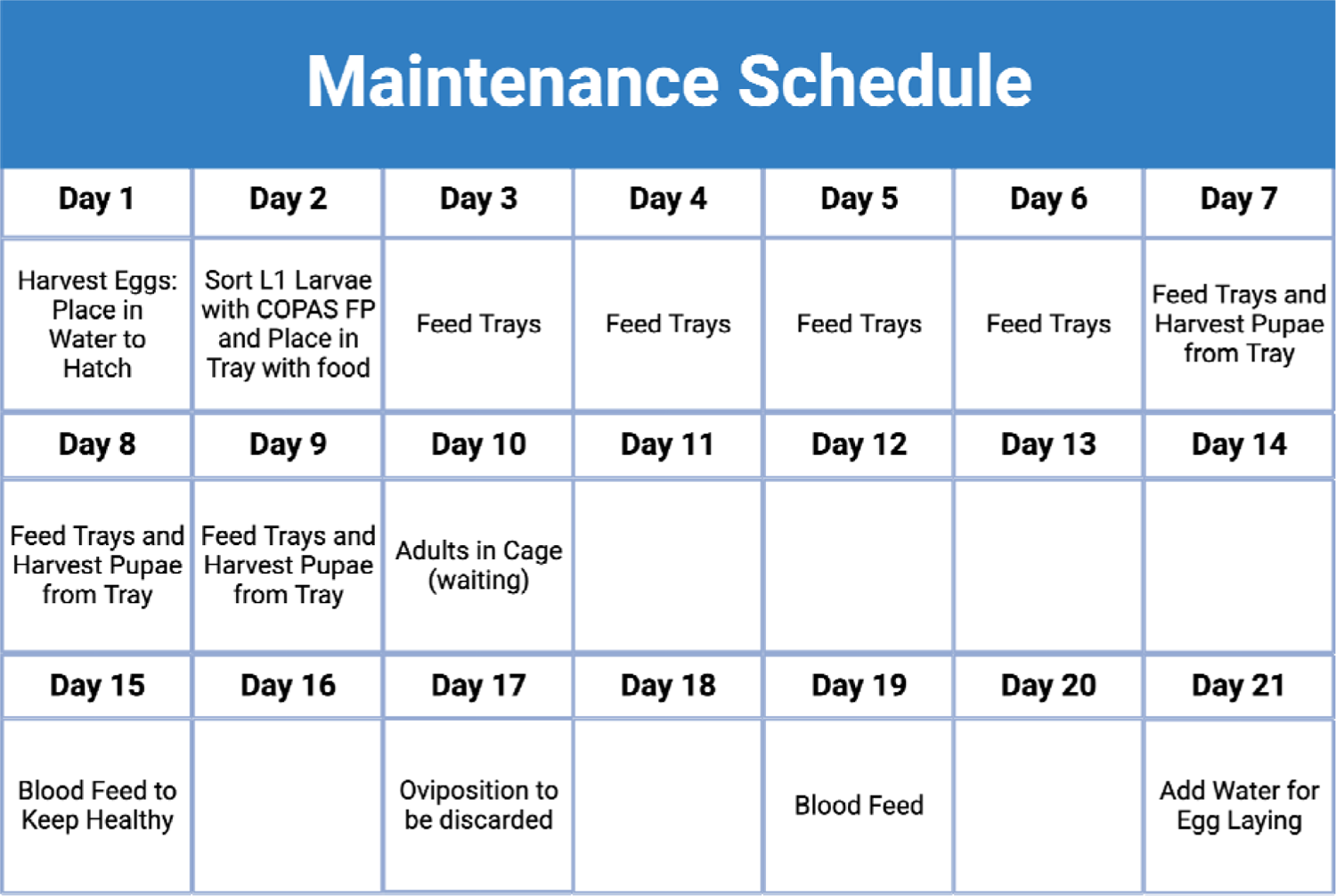
Maintenance Phase schedule.

Mathematical modeling suggests that only 14 weeks of active mosquito releases and an additional six weeks to scale production to the Active Phase is required. This scheme leaves 34 weeks of the year during which *Anopheles gambiae* mosquitoes are maintained at low population sizes. This phase utilizes the standard rearing schedule but stretches the rearing procedures to a three week cycle to minimize food, water, and labor inputs. This phase differs from the Active Phase in that it restarts a cycle once every three weeks. In the Active Phase, a new production cycle occurs daily to maximize efficiency. Figure generated in BioRender.com.

##### 2.1.4.2 Ramping phase

When transitioning from the Maintenance Phase to the Active Phase, a six week Ramping Phase is required to build the colony to support the egg production numbers required in the Active Phase (**Fig. 15 and 16**). While this facility does not need six weeks to expand the colony to meet the production requirements for the Active Phase, the six week Ramping Phase is necessary to synchronize daily production. This ramp-up of the colony is accomplished by increasing the blood feeding and egg harvesting frequencies. The last Maintenance Phase batch of mosquitoes is blood fed two additional times to maximize egg harvest and spread egg production across multiple days instead of weeks. This change starts the Ramping Phase, with each of these harvests being reared and harvested across three days per week. These collections are staggered to provide better coverage of the entire three week cycle. This strategy will provide nine cycles that can be further expanded to 27 individual cycles at the end of the Ramping Phase. At this point, eggs will be produced each day, providing complete coverage for the Active Phase.

**Fig. 15:**
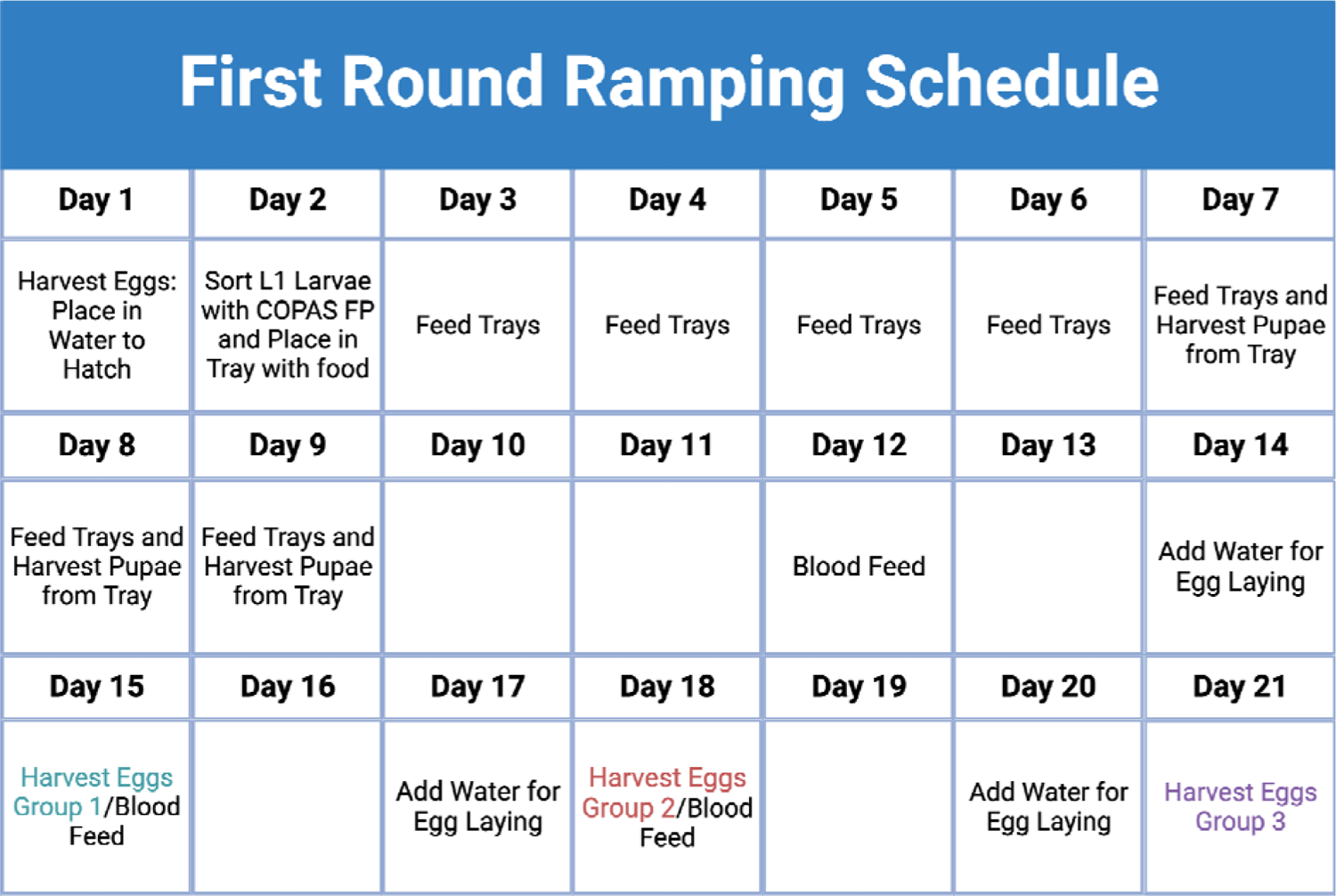
Maintenance to Ramping Phase schedule.

**Fig. 16:**
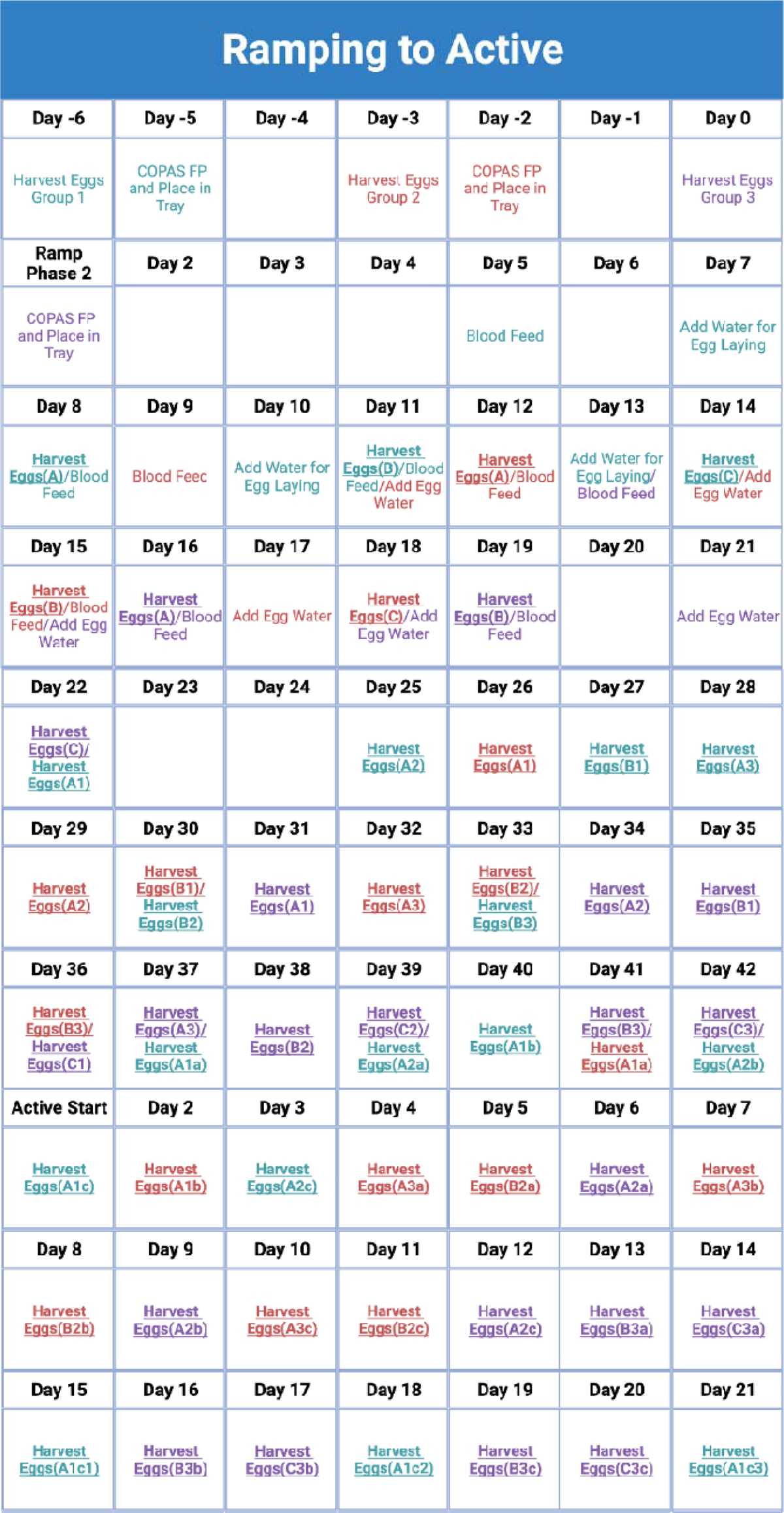
Ramping Phase to Active Phase Schedule.

The first step in the transition from the Maintenance Phase to the Ramping Phase maximizes egg production in the late Maintenance Phase while also spreading the production out across multiple days. This transition is important because production cycles need to begin daily during the late Ramping Phase before the transition to the Active Phase (**Fig. 17**) and requires spreading production across a three week cycle. Figure generated in BioRender.com.

**Fig. 17:**
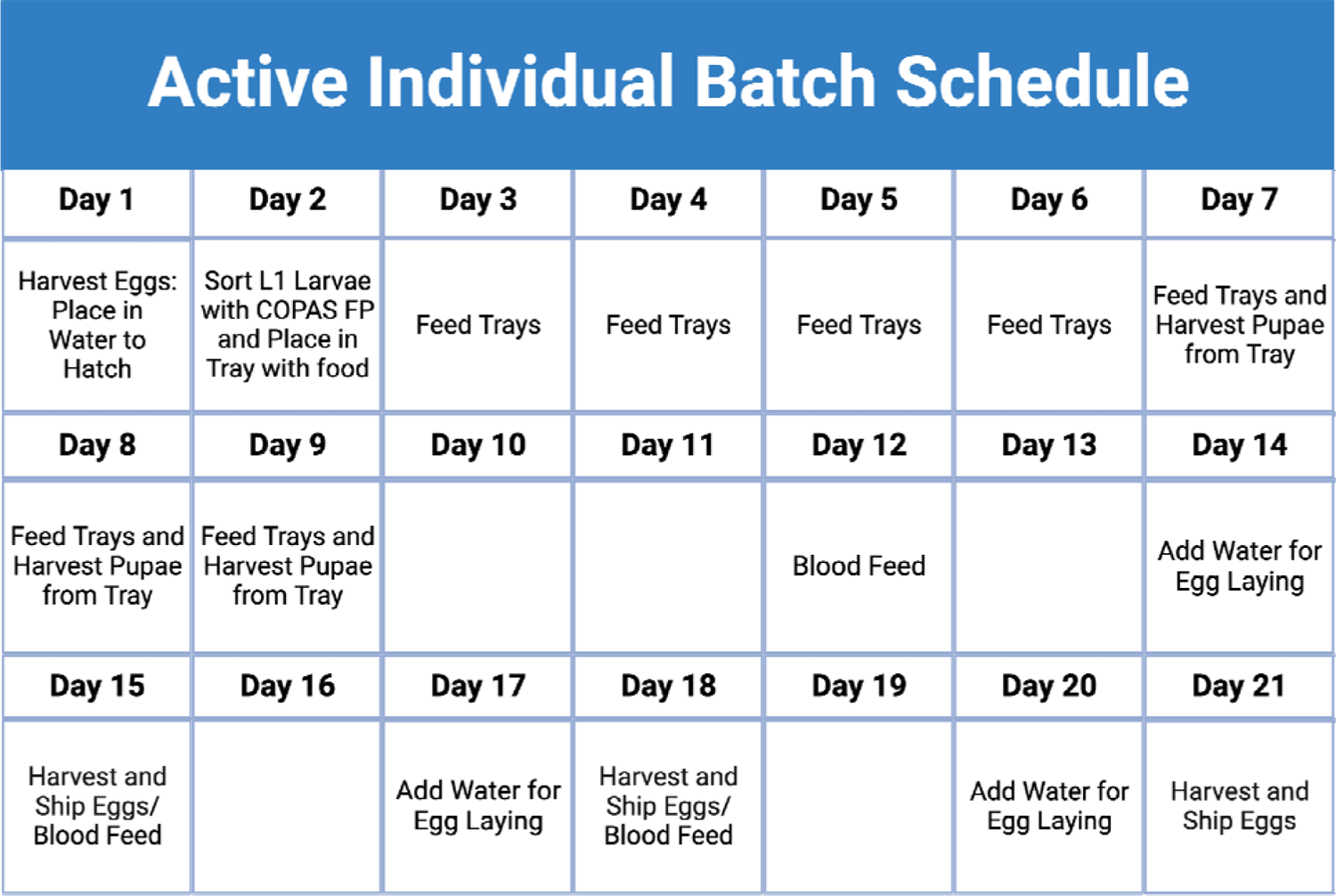
Active Phase Schedule.

The Ramping Phase’s goal is to increase the *A. gambiae* colony size to support the egg production requirements in the Active Phase, while also expanding to daily egg production. While this level of production does not require expansion, the early and late Ramping Phases can increase the population by 50 to 100-fold per egg lay depending on the estimated larval survival to adult. This could result in a 1.25e^5^ to 1e^6^ increase in daily mosquitoes, suggesting that Maintenance Phase population levels can remain low as needed. There is a lower limit of the mosquito population during the Maintenance Phase as genetic diversity needs to be maintained. The Ramping Phase also demonstrates that egg production can rapidly be transitioned from once every three weeks to daily in three week cycles once the Active Phase begins. There is an overlap in the egg batches for many days in the Active Phase, so to simplify this, redundant egg hatchings are excluded for clarity. Figure generated in BioRender.com.

##### 2.1.4.3 Active Phase

The Active Phase has a new cycle (or cohort) of mosquitoes being hatched and sorted each day (**Fig. 17**). Importantly, this schedule is designed to minimize the number of COPAS sorting machines required to produce the weekly egg requirements. The COPAS Sorting Machines are the highest cost equipment for this project and are limited in the number of larvae that can be sorted in 24 hours. Therefore, it is essential to maximize the efficiency of each machine. To this end, production is staggered across every day of the week during the full three week cycle. A daily production schedule also reduces the total number of rearing racks, cages, and drones needed to obtain production numbers. This schedule has additional benefits by standardizing lab procedures and staff roles. Staff will specialize in a particular mosquito rearing role, which should increase efficiency.

Rearing mosquitoes in a large facility is a complex and critical process that requires standardized protocols and a team of skilled staff. A well-trained management hierarchy will be needed to ensure uniformity in practices and to meet the facility’s production goals.

Each day in the Active Phase, a single batch of mosquitoes follows the three week schedule shown above. Rather than attempting to produce the weekly requirement of mosquitoes in one group, a new batch of mosquitoes is started in this cycle each day of the Active Phase. By spreading this production across each day, the number of mosquitoes at each stage of the lifecycle are reduced to a minimum, and the facility can maximize efficiency. For example, if the weekly requirement for sex sorting was used in one day, we would require seven times the number of COPAS FP 500 sorting machines to meet this demand. By spreading this across the week, the facility can be reduced to a seventh of the size. Each batch of mosquitoes is blood fed three times to produce three batches of eggs across the lifespan of this batch of mosquitoes. The youngest cohort will produce, on average, the most viable eggs (118 per female), the cohort on their second egg lay will produce 101 viable eggs, and the oldest cohort will produce 86 viable eggs. This estimate means that each day during the active phase, the facility produces the same number of eggs per day as one cohort of mosquitoes will produce during their lifetime. Figure generated in BioRender.com.

##### 2.1.4.4 Conclusions: Phase Summary

The implementation of a phased production plan offers significant benefits in terms of minimizing resource and labor costs during specific periods when the sterile pgSIT males are not being released. The plan consists of a 34-week Maintenance Phase, a 6-week Ramping Phase, and a 12-week Active Phase annually.

During the 34-week Maintenance Phase, the facility focuses on colony maintenance and preparation for future production. This phase involves insect rearing, monitoring mosquito health and development, and ensuring the overall health and quality of the colony. While this phase does not involve the daily production required for releases, it plays a crucial role in maintaining a healthy and productive population of sterile pgSIT males.

Following the Maintenance Phase, the facility enters the 6-week Ramping Phase. This phase serves as a transition period to shift from colony maintenance to the active production required for releases. Within these six weeks, the facility gears up its operations, increasing the daily production rate to meet the demand. This phase includes scaling up the breeding efforts, optimizing production processes, and streamlining workflows to efficiently generate the required number of sterile pgSIT males.

By dividing the production process into these distinct phases, the facility can optimize its resource allocation and labor utilization. During the Maintenance Phase, when sterile pgSIT males are not released, the focus is on maintaining the colony and ensuring its long-term sustainability. This approach minimizes unnecessary production costs during this period. In contrast, the Ramping Phase allows the facility to swiftly transition from the maintenance phase to full-scale production, ensuring a seamless shift in operations while minimizing any production gaps.

Overall, the phased production plan ensures that resources and labor are efficiently utilized, reducing costs during periods when releases are not required and facilitating a smooth and efficient transition to meet the necessary production demands during the active phase.

### 2.2 Predicted health benefits of implementing pgSIT in The Upper River Region of The Gambia

The health benefit modeling methods were discussed previously in this paper, but the annual health benefits, the annual cases, and the deaths averted were not addressed in detail. These calculations are useful for interpreting the direct health benefits of the pgSIT technology. In this model, these epidemiological outcomes are age-stratified, which facilitates the calculation of the number of life years saved due to the pgSIT intervention. While the value of life estimate used in this study does not take age into account, the number of life years saved does include this important metric (**Table 29 and Table 30**). There are multiple ways to calculate the value of life, each having intrinsic biases in their calculations. A common value of life calculation accounts for the number of life years lost. This calculation method is favorable for malaria prevention as malaria primarily kills children. We use life-years to estimate QALYs in addition to the other measurements.

**Table 29:** Estimated age stratified life years saved.

| **Age Bracket** | **0 to 5 Years** | **5 to 17 Years** | **17 to 40 Years** | **40 to 60 Years** | **60 Years and Older** |
| --- | --- | --- | --- | --- | --- |
| **Average Age** | 2.5 | 11 | 28.5 | 50 | N/A |
| **Years Saved** | 60.1 | 51.6 | 34.1 | 12.6 | 1 |

**Table 30:** Life years saved annually.

| Intervention<br>Year | 0-5 years | 5-17 years | 17-40<br>years | 40-60<br>years | ≥60 years | Total |
| --- | --- | --- | --- | --- | --- | --- |
| 0* | 3 | 6 | 1 | 0 | 0 | -5 |
| 1 | -2 | -5 | 0 | 0 | 0 | 133 |
| 2<sup>#</sup> | 2,855 | 5,897 | 556 | 43 | 2 | 9,353 |
| 3 | 3,446 | 7,2790 | 689 | 53 | 3 | 11,461 |
| 4 | 3,433 | 7,389 | 705 | 54 | 3 | 11,584 |
| 5 | 3,443 | 7,538 | 725 | 55 | 3 | 11,763 |

\* Year 0 has no intervention.

**Table 31:** Sick days prevented by age over four years of pgSIT interventions.

| Intervention<br>Year | 0-5 years | 5-17 years | 17-40<br>years | 40-60<br>years | ≥60 years | Total |
| --- | --- | --- | --- | --- | --- | --- |
| 0* | 2 | 17 | 21 | 4 | 3 | 47 |
| 1 | -1 | -11 | -14 | -3 | -2 | -31 |
| 2<sup>#</sup> | 1,673 | 13,974 | 18,045 | 3,599 | 2,198 | 39,488 |
| 3 | 2,020 | 16,897 | 21,887 | 4,374 | 2,673 | 47,851 |
| 4 | 2,012 | 16,835 | 21,844 | 4,374 | 2,678 | 47,743 |
| 5 | 2,016 | 16,879 | 21,934 | 4,400 | 2,698 | 47,928 |

\* Year 0 has no intervention.
<sup>#</sup> Year 2 the intervention begins part way through the year and this model assumes an accumulative suppressive effect, year to year.

The life years saved are calculated by subtracting the average age of the age bracket from the life expectancy in The Gambia, which is 62.6 years. For the 60 and older category, the average is not easily attainable and is expected to be over the average life expectancy for the region. With this in mind, we assume that those who die of malaria in this category died one year before their time.

To calculate the number of life years saved per age group per year (**Table 30**), the years saved in each category is then multiplied by the deaths prevented in **Table 2**. The majority of the life years saved are in the younger age groups, reflecting the results in **Table 2**, which shows the number of years of life that are being saved for each category.

The number of sick days prevented is another quantifiable health outcome that can be extrapolated from the modeling data. The incapacitation rate, which divides the number of work days lost by the total number of absentees, is an effective means to estimate the average number of sick days per case of malaria (**Table 31**). A study that monitored the economic effects of worker absences due to malaria calculated an average incapacitation rate of 3.7 days (Lukwa et al. 2019). To calculate the number of sick days saved due to the pgSIT intervention, the cases prevented in Table 1 are multiplied by the average sick day rate (**Table 31**).

Once this intervention has been applied for multiple years, approximately 48,000 sick days are saved (**Table 31**). Sick days have a direct economic impact on the community, but it is harder to quantify the indirect effects of sick days, such as the loss of educational opportunities for school-aged children. Notably, sick days in the 0 to 5 years and 5 to 17 years groups also cause economic loss if parents need to take time off to care for a sick child.

### 2.3 Quantifying the economic benefits of pgSIT in The Gambia

Five predicted values can be estimated when quantifying the economic benefits of a malaria disease intervention: (1) the value of life saved per year, (2) the value of sick days saved, (3) medical intervention costs saved, (4) insecticide intervention costs reduced and (5) additional economic benefits. Many of these values are directly quantifiable.

#### 2.3.1 Value of statistical life (VSL) saved per year

To quantify the value of lives saved per year, the number of lives saved and VSL saved in The Gambia must be identified. As shown in the predicted health benefits of implementing pgSIT in The URR of The Gambia, approximately 230 lives would be saved annually (**Table 2**). This number then needs to be applied to the VSL value. VSL is the local tradeoff between risk of death and money which is then used to derive an estimate of local money that would be paid to prevent a death. For example, if individuals willingly pay 1,000 USD in extra risk prevention measures to reduce an annual one in 10,000 chance of death to 0, then 1,000 USD would be multiplied by 10,000 to get a VSL of 10 million USD. Studies evaluate the VSL for many risks, and the USTD uses a selection of these calculations with economic adjustments to estimate the VSL in the United States. For 2022, the United States VSL is estimated to be 12.5 million USD (“Departmental Guidance on Valuation of a Statistical Life in Economic Analysis” n.d.).

Without a direct estimate of the VSL for The Gambia, we use the US VSL as a benchmark, a practice used frequently in the literature (Viscusi 2020). Therefore, we used the United States Transportation Department (USTD) VSL and converted the GDP per capita purchasing power parity (PPP) in The Gambia to the United States GDP per capita(Kip Viscusi and Masterman 2017).

This method of VSL calculation can be used to estimate the VSL in The Gambia by comparing the GDP per capita and adjusting by PPP, a metric to equalize purchasing power between countries. This calculation is done because the per capita GDP is linked to the VSL for a region. GDP per capita adjusted by PPP affects the VSL as the amount able to invest into the cost associated with a reduction in death is limited by budget. Therefore, comparing the United States per capita GDP to The Gambia’s per capita GDP adjusted by PPP can determine the VSL for The Gambia. The Gambia’s GDP per capita adjusted by PPP is 2,215 USD, and the United States GDP per capita is 70,248 USD. Utilizing these values and the previously shown United States VSL, The Gambia has a VSL of 394,139 USD. With an average of 230 lives saved yearly, this pgSIT intervention saves approximately 91 million USD in VSL (**Table 32**). When considering the cost of this program ranges from 315,000 to 318,000 USD annually (**Table 28**), the annual cost of this program would save lives at about 0.3% of the VSL.

**Table 32:**
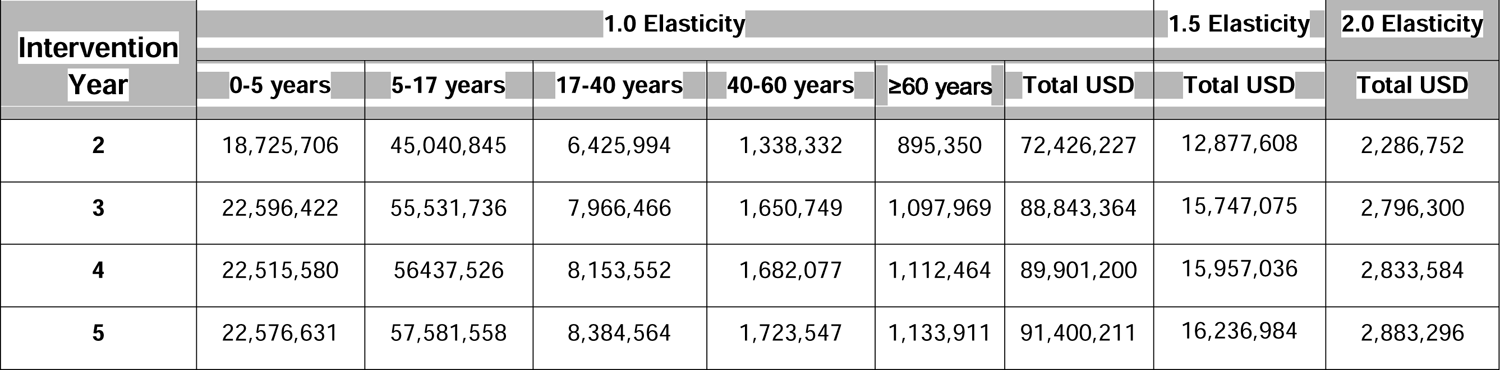
Age-stratified and total value of life saved in USD by VSL.

The VSL can change with income. Income elasticity measures the percent change of VSL as income changes by 1%. Based on VSL studies in multiple countries, lower per capita income nations have a higher percentage of their income allocated to cost of living and, therefore, will have less to spend on extra methods to reduce the risk of death and disease.

Thus, VSL increases with income. To capture the variation in how this is quantified, we applied elasticity of 1.0, 1.5 and 2.0 in order to capture the range of potential VSL estimates (**Table 32)**. This approach uses The Gambian PPP adjusted GDP, dividing this by the United States’ GDP, and raising this fraction to a power based on the elasticity (1, 1.5, 2). The VSLs for The Gambia at elasticities of 1.0, 1.5 and 2.0 will be 394,139, 69,987 and 12,428 USD, respectively. This range is extreme, however, as income elasticities of greater than 1.5 are unlikely. Direct VSL studies in sub-saharan Africa are uncommon and we could not find a study in The Gambia to compare to these estimates. Masterman (2017) suggests that an income elasticity of 1 is most appropriate for assessing most countries outside of the US and thus the age group and total estimates are appropriate at an elasticity of 1 (Kip Viscusi and Masterman 2017). Fortunately, even with the most extreme income elasticity, this intervention would still be cost-effective if calculating the cost of VSL saved per investment **(Table 42)**.

Additionally, QALY and WTP attempt to capture a further range of potential benefit estimates, with WTP being related to VSL in that both are an attempt to capture local willingness to prevent the risk of death.

#### 2.3.2 Value of Life Years based on Quality Adjusted Life Years (QALY) saved annually

In addition to calculating the value of life saved via the VSL, the value of life saved can be calculated by the number of life years saved and applying a value to each year saved. The average value for the loss of a quality adjusted life year (QALY) in the United States is 104,000 USD (Vanness, Lomas, and Ahn 2021). This value is calculated in an assortment of ways, but at the core, it is an estimate of the payment willingly made to increase life expectancy by one QALY. This estimate is primarily based on insurance rates, but other methods have been explored. This method of life measurement has been primarily used in the United States and Europe, but a similar approximation to VSL could be used for QALY to take into account the investment that would be made in The Gambia to gain one QALY. Again factoring in GDP differences and the PPP, as seen in the VSL section, the same equation can be used to get a QALY value of 3,280 USD. This value could almost be doubled by using the 200,000 USD QALY, bringing it to 6,308 USD. The QALY values can then be multiplied by the life years saved annually to show the value of life saved using the QALY metric (**Table 33 and Table 34**).

**Table 33:**
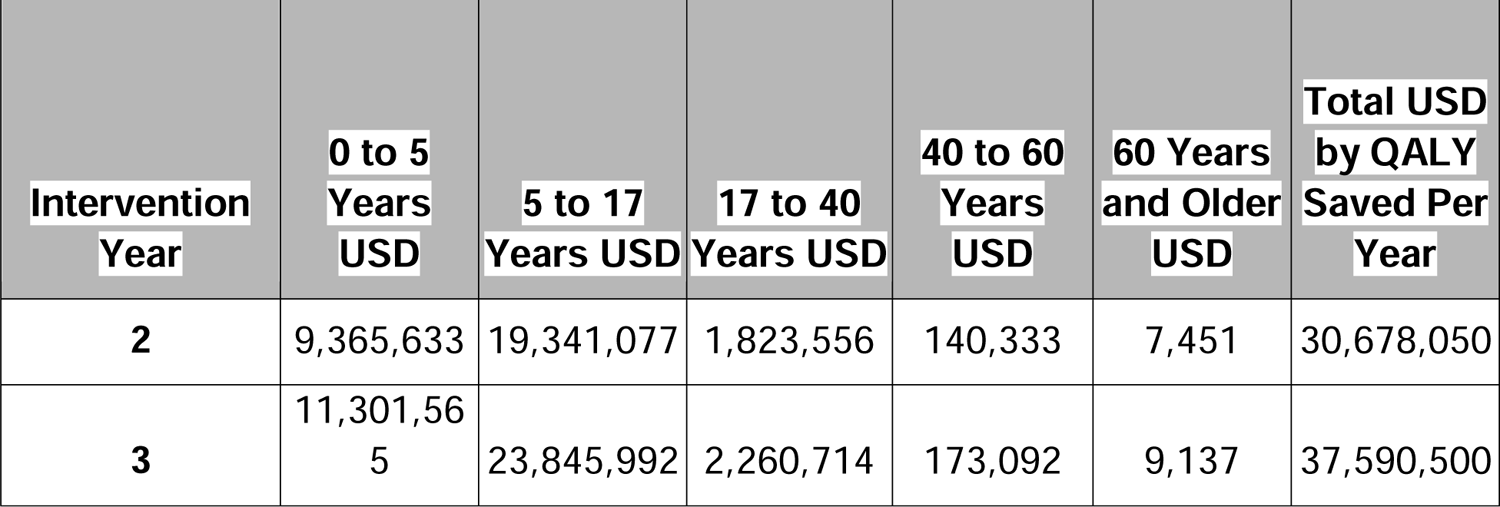
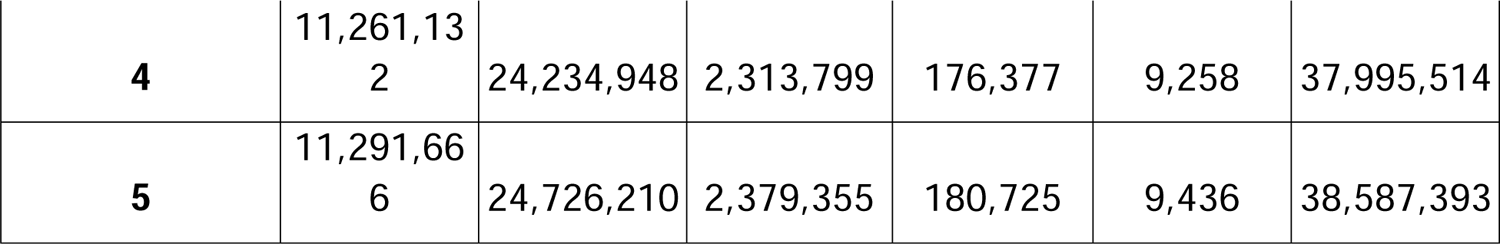
Value of life saved in USD based on average QALY value.

**Table 34:**
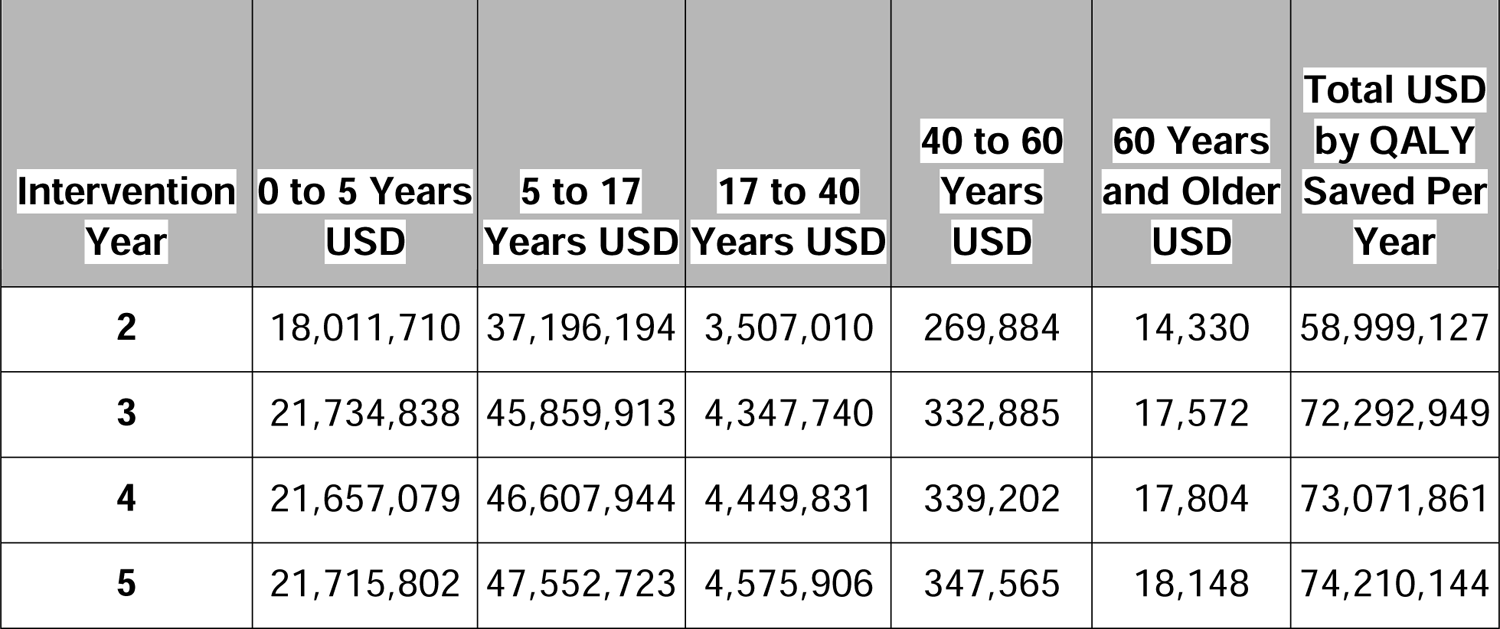
Value of life saved in USD based on high QALY value.

| **Intervention Year** | **0 to 5 Years USD** | **5 to 17 Years USD** | **17 to 40 Years USD** | **40 to 60 Years USD** | **60 Years and Older USD** | **Total USD by QALY Saved Per Year** |
| --- | --- | --- | --- | --- | --- | --- |
| **2** | 18,011,710 | 37,196,194 | 3,507,010 | 269,884 | 14,330 | 58,999,127 |
| **3** | 21,734,838 | 45,859,913 | 4,347,740 | 332,885 | 17,572 | 72,292,949 |
| **4** | 21,657,079 | 46,607,944 | 4,449,831 | 339,202 | 17,804 | 73,071,861 |
| **5** | 21,715,802 | 47,552,723 | 4,575,906 | 347,565 | 18,148 | 74,210,144 |

**Table 35:**
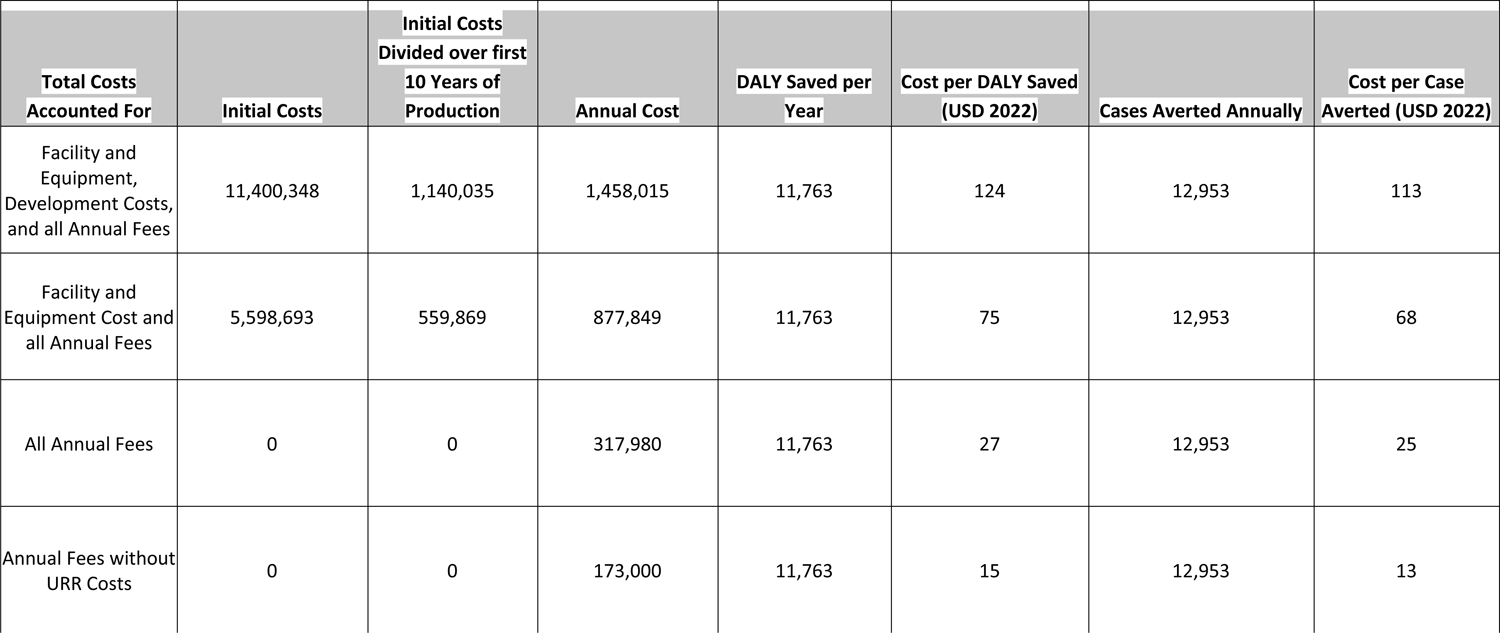
Cost per life year saved and cost per case averted.

This QALY estimate, while significantly less than the VSL estimate, will still pay off the investment in the first year of intervention. This variation is likely due to these methods differing in their means of calculating the value of human life. In the following years, the intervention will cost less than 0.8% of the average QALY saved annually.

#### 2.3.3 Cost per DALY Saved and Cost per Case Averted by Intervention Annually

In addition to utilizing life years saved to monetize the value of this intervention, determining the cost per disability adjusted life year (DALY) saved is a means to compare this intervention to other health interventions. For pgSIT, there are several costs useful for these comparisons, because current interventions do not include facility upstart and research and development costs and determining annual costs is important to compare the cost per life year saved. Every cost will factor in annual costs of the facility, but the other estimates will add the costs of rearing in the URR, along with start up costs and an estimate that includes the research and development costs. The first estimate includes all costs and represents the cost per DALY of the first 10-years of the first facility. The second estimate removes the research and development costs involved and would represent the second facility of the same size being developed. The third estimate only includes the annual costs, which includes both the facility production costs and rearing costs of the URR. The last estimate only includes the facility production costs as the URR rearing costs may be covered by volunteer labor as many LLIN costs are covered by and feed for mosquitoes could be sourced locally from detritus that the wild mosquitoes feed on. The initial costs for the first 2 estimates are spread over the first 10 years of the facility. These 4 estimates allow for comparison to a variety of interventions throughout the use of this intervention.

These costs are then applied to the predicted DALY averted or case averted. We can then provide cost estimates per DALY averted or per case averted. The cost per DALY averted and cost per case averted can then be compared to other interventions, both malaria interventions and other life saving interventions when considering the cost per DALY averted. This can be used to compare the cost efficiencies of these interventions to save lives or prevent disease.

This table presents four estimates of cost per DALY saved or cases averted based on the following: Cost based on the annual cost of the 1) Total Costs including Research and Development, Facility Construction and Annual Fees, 2) Costs of Facility Construction and Annual Fees, 3) Annual Fees including the URR costs and 4) Annual Fees including only the facility production costs. This metric will be used to compare the cost of saving life years or preventing illness compared to other interventions. These calculations are done in 2022 USD and will require conversion if compared to calculations using USD from other years.

#### 2.3.4 GDP growth benefit associated with malaria prevention

The effect of malaria on the growth and development of countries is commonly understood to be one of the primary sources of poverty in Africa and in other tropical regions affected by the disease (Sachs and Malaney 2002). Malaria has an estimated annual reduction in the growth of GDP by 1.3% (Gallup and Sachs 2001). Additionally, there are examples where only a 10% reduction in malaria results in a 0.3% increase in GDP (Gallup and Sachs 2001). This 10% reduction is convenient as the target region of The URR is 11.1% of the population, meaning that reducing malaria by more than 10% in this region would, therefore, result in at least a 0.3% increase in GDP. Accounting for growth benefits in this way is unique compared to other methods in that it compounds annually upon deployment and results in exponential growth and benefits from this intervention. This intervention will likely be deployed by 2030 as several field trials require completion before wide scale implementation. Utilizing the current 10-year average GDP growth rate of 3.5%, we predict the expected growth rate until 2030. The GDP without intervention and GDP with intervention then diverge, with the column without intervention maintaining the predicted GDP rate and the GDP with intervention a 3.8% growth rate. The annual effect of intervention is then the difference in GDP between the predictions with and without intervention. It is expected that this intervention will have an exponential effect on the economic growth of the country (**Table 36**).

**Table 36:** Predicted GDP increase with pgSIT intervention in the Upper River Region.

| **Year** | **GDP without Intervention (USD)** | **GDP with Intervention (USD)** | **Annual Effect of Intervention (USD)** |
| --- | --- | --- | --- |
| **2021 GDP of The Gambia** | 2,038,000,000 | 2,038,000,000 | Pre-intervention |
| **Predicted 2022** | 2,109,330,000 | 2,109,330,000 | Pre-intervention |
| **Predicted 2023** | 2,183,156,550 | 2,183,156,550 | Pre-intervention |
| **Predicted 2024** | 2,259,567,029 | 2,259,567,029 | Pre-intervention |
| **Predicted 2025** | 2,338,651,875 | 2,338,651,875 | Pre-intervention |
| **Predicted 2026** | 2,420,504,691 | 2,420,504,691 | Pre-intervention |
| **Predicted 2027** | 2,505,222,355 | 2,505,222,355 | Pre-intervention |
| **Predicted 2028** | 2,592,905,138 | 2,592,905,138 | Pre-intervention |
| **Predicted 2029** | 2,683,656,817 | 2,683,656,817 | Pre-intervention |
| **Predicted 2030** | 2,777,584,806 | 2,785,635,776 | 8,050,970 |
| **Predicted 2031** | 2,874,800,274 | 2,891,489,936 | 16,689,662 |
| **Predicted 2032** | 2,975,418,284 | 3,001,366,553 | 25,948,270 |
| **Predicted 2033** | 3,079,557,924 | 3,115,418,482 | 35,860,559 |
| **Predicted 2034** | 3,187,342,451 | 3,233,804,385 | 46,461,934 |
| **Predicted 2035** | 3,298,899,437 | 3,356,688,951 | 57,789,515 |
| **Predicted 2036** | 3,414,360,917 | 3,484,243,132 | 69,882,215 |
| **Predicted 2037** | 3,533,863,549 | 3,616,644,371 | 82,780,821 |
| **Predicted 2038** | 3,657,548,773 | 3,754,076,857 | 96,528,083 |
| **Predicted 2039** | 3,785,562,980 | 3,896,731,777 | 111,168,797 |
| **Predicted 2040** | 3,918,057,685 | 4,044,807,585 | 126,749,900 |
| **Predicted 2041** | 4,055,189,704 | 4,198,510,273 | 143,320,569 |
| **Predicted 2042** | 4,197,121,343 | 4,358,053,663 | 160,932,320 |
| **Predicted 2043** | 4,344,020,590 | 4,523,659,703 | 179,639,112 |
| **Predicted 2044** | 4,496,061,311 | 4,695,558,771 | 199,497,460 |
| **Predicted 2045** | 4,653,423,457 | 4,873,990,005 | 220,566,548 |

This increase in GDP may be underestimated if this system is leveraged to suppress malaria throughout the region. Without the concern of malaria infection in the URR, it is foreseeable that outside investment into this region would be considerable.

Using the growth benefit of GDP alone, the economic benefits of this intervention pay off the investment in only two years. The valuation of these methods likely has some, but not complete, overlap making the addition of these values inappropriate and incomplete estimates of the benefits.

#### 2.3.5 Value of sick days saved per year

In addition to the VSL, there are approximately 48,000 sick days saved annually. This savings is due to preventing nearly 9,000 cases per year in the URR. There are multiple ways to estimate the value per sick day, but a common approach is taking the QALY and applying this daily. We have shown this with previous estimates converted to The Gambia values. This can be converted to a daily value of 12.96 USD and then be used to calculate the total QALY saved per year by preventing malaria infections. On average, 620,000 USD is saved in QALY annually from the pgSIT intervention. While the main value saved is the VSL, it is notable that this value saved will have a more immediate impact on the economy by saving these sick days in that year, whereas the VSL value is spread across many years (**Table 37**). The value of the sick days saved is nearly double that of the annual cost to run this program.

**Table 37:** Value of sick days saved.

| Intervention<br>Year | 0-5 years | 5-17 years | 17-40<br>years | 40-60<br>years | ≥60 years | Total |
| --- | --- | --- | --- | --- | --- | --- |
| 2 | 21,688 | 181,100 | 233,858 | 46,644 | 28,480 | 511,770 |
| 3 | 26,174 | 218,991 | 283,652 | 56,684 | 34,646 | 620,147 |
| 4 | 26,070 | 218,187 | 283,101 | 56,684 | 34,703 | 618,745 |
| 5 | 26,133 | 218,749 | 284,267 | 57,025 | 34,971 | 621,144 |

#### 2.3.6 Value of medical intervention saved per year

There are also medical intervention costs saved by malaria case prevention. Potentially all of these medical costs can be reduced if pgSIT eliminates malaria in the region, but this is uncertain until large-scale field testing is completed. The immediate cost that should be reduced is the estimated medical intervention expenditure. For the URR, it is estimated that approximately 36,000 USD is spent on medical interventions to treat malaria (**Table S3**). This approximation is likely an underestimate as this only accounts for governmental investment in treatment and does not include individual costs. This cost for medical intervention would likely scale with malaria cases, but even assuming this cost remains static, there is the additional cost paid by patients seeking treatment. On average, each patient is paid 4.55 USD for treatment in addition to lost wages and the cost of medication (**Table 38**) (Broekhuizen et al. 2021). This result leads to approximately 59,000 USD being saved annually (**Table 39**). All of the medical intervention or associated costs will be immediately saved if the pgSIT intervention is as effective as predicted, as these medical interventions will not be required.

**Table 38:** Costs associated with malaria treatment seeking.

| **Years** | **Complicated Cases** | **Uncomplicated Cases** | **Household Costs for seeking treatment GBP** | **Household Costs for seeking treatment USD** | **Average Per Household per Case USD** | **Average GBP to USD by Year** |
| --- | --- | --- | --- | --- | --- | --- |
| **2013** | 1,515 | 45,479 | 145,021 | 226,233 | 4.81 | 1.56 |
| **2014** | 933 | 26,806 | 85,613 | 141,261 | 5.09 | 1.65 |
| **2015** | 1,273 | 52,007 | 164,288 | 251,361 | 4.72 | 1.53 |
| **2016** | 1,262 | 34,167 | 109,368 | 147,647 | 4.17 | 1.35 |
| **2017** | 526 | 13,821 | 44,293 | 57,138 | 3.98 | 1.29 |
| **Average** | 1,102 | 34,456 | 109,717 | 164,728 | 4.55 | 1.48 |

**Table 39:** Treatment seeking costs saved from preventing malaria cases.

| **Intervention Year** | **0-5 years** | **5-17 years** | **17-40 years** | **40-60 years** | **≥60 years** | **Total** |
| --- | --- | --- | --- | --- | --- | --- |
| **2** | 2,058 | 17,184 | 22,190 | 4,426 | 2,702 | 48,560 |
| **3** | 2,484 | 20,779 | 26,915 | 5,379 | 3,287 | 58,844 |
| **4** | 2,474 | 20,703 | 26,862 | 5,379 | 3,293 | 58,711 |
| **5** | 2,480 | 20,756 | 26,973 | 5,411 | 3,318 | 58,938 |

**Table 40:** Additional potential cost savings: Value saved by URR intervention per year.

|  | Medical<br>Intervention<br>Saved USD | IRS/LLIN<br>Interventions<br>Saved (Potential<br>Savings) |
| --- | --- | --- |
| Average<br>Annual Cost | 36,000 | 310,000 |

**Table 41:** Willingness-to-pay for malaria prevention in the URR.

| Year | Population Estimate | Willingness to Pay<br>USD |
| --- | --- | --- |
| 2022 | 300,149 | 4,541,254 |
| 2023 | 307,713 | 4,655,698 |
| **2024** | 315,467 | 4,773,016 |
| **2025** | 323,417 | 4,893,299 |
| **2026** | 331,567 | 5,016,609 |
| **2027** | 339,922 | 5,143,020 |
| **2028** | 348,488 | 5,272,623 |
| **2029** | 357,270 | 5,405,495 |
| **2030** | 366,273 | 5,541,710 |
| **2031** | 375,503 | 5,681,360 |
| **2032** | 384,966 | 5,824,536 |
| **2033** | 394,667 | 5,971,312 |
| **2034** | 404,613 | 6,121,795 |
| **2035** | 414,809 | 6,276,060 |
| **2036** | 425,262 | 6,434,214 |
| **2037** | 435,979 | 6,596,362 |
| **2038** | 446,966 | 6,762,596 |
| **2039** | 458,229 | 6,933,005 |
| **2040** | 469,777 | 7,107,726 |

**Table 42:**
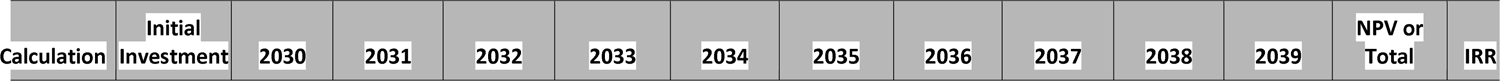
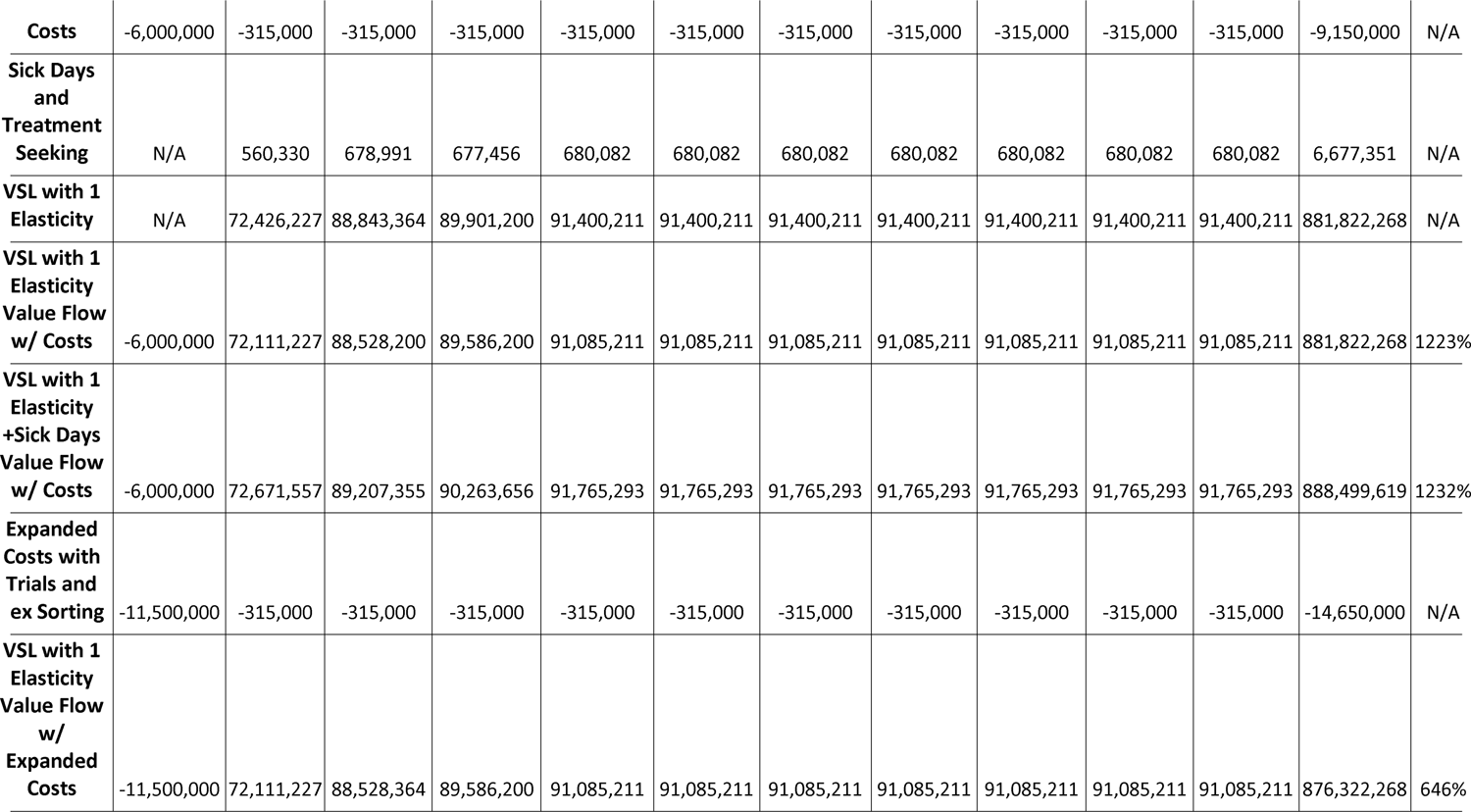
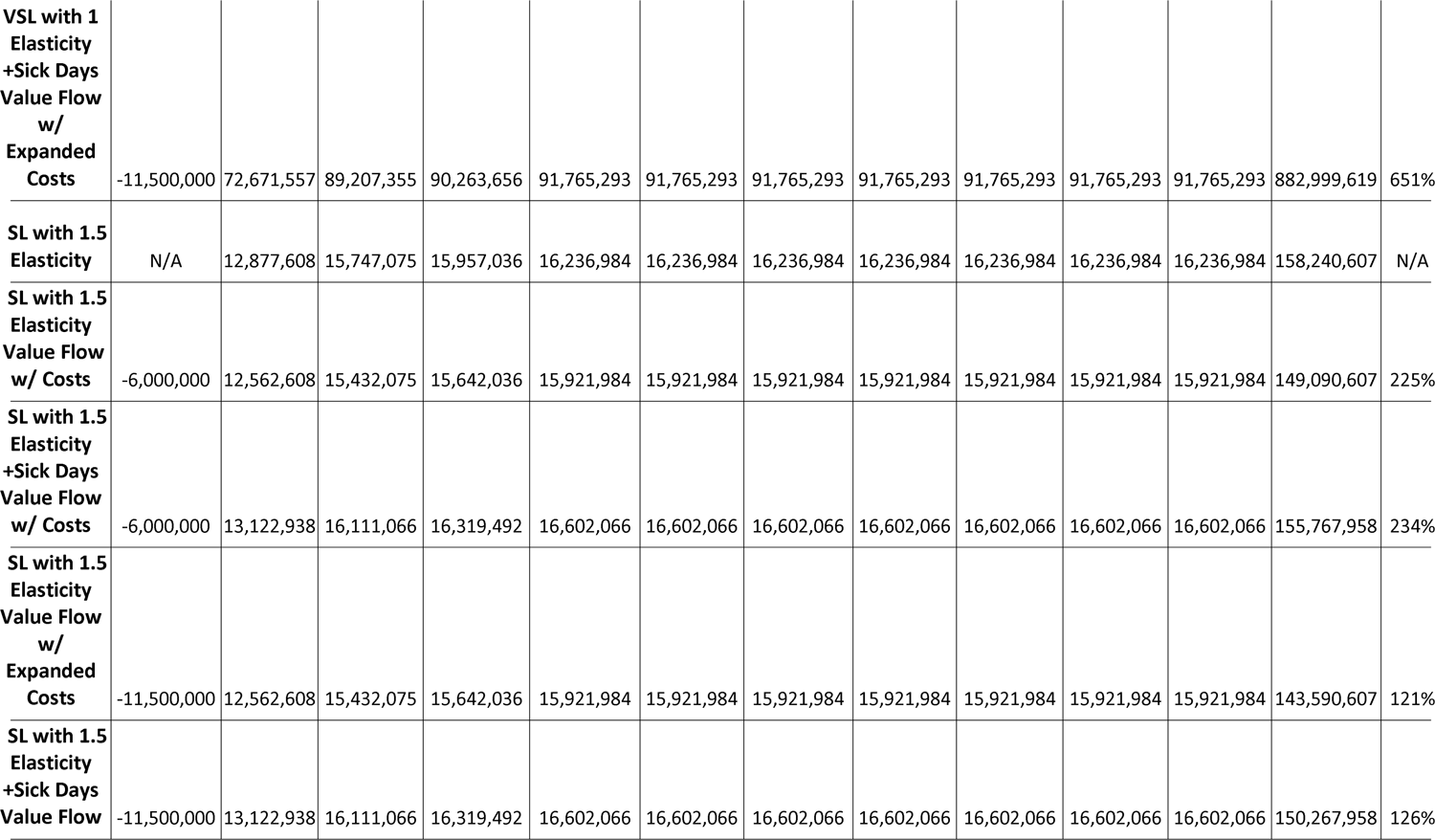
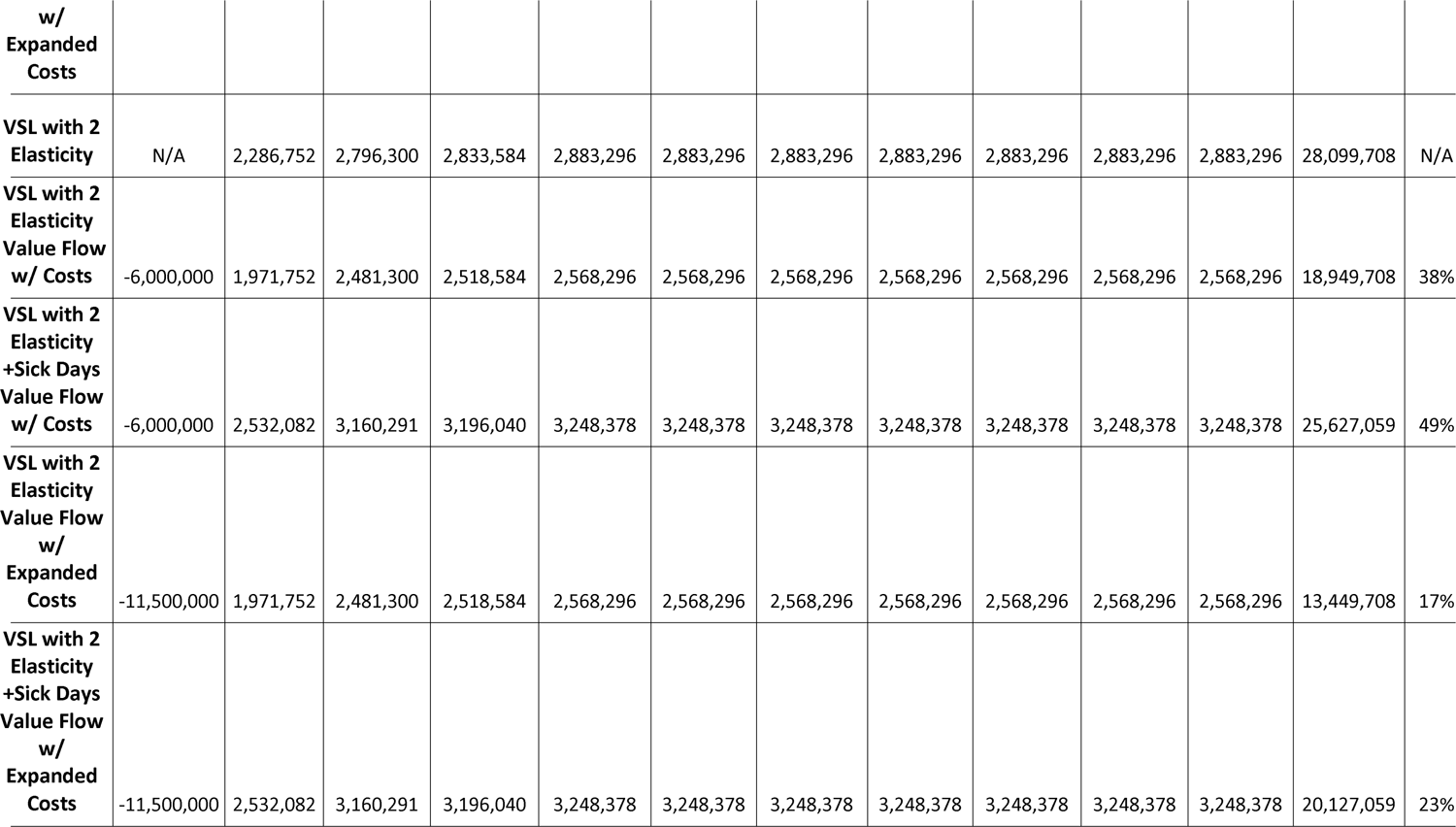
Net present value (NPV) over 10 years and internal rate of return (IRR): value of statistical life (VSL)

Adapted from(Broekhuizen et al. 2021) with conversions from British pound (GBP) to USD.

#### 2.3.7 Value of IRS and LLINs saved per year

The current cost of malaria interventions for the URR of The Gambia is approximately 310,000 USD when excluding the medical intervention costs (**Tables 38, S3**). This cost will be reduced if pgSIT alone is sufficient for mosquito suppression in The Upper River Region. The model predicts the local extinction of the mosquito population (**Fig. 3**). Therefore, interventions that are currently being funded, including long lasting LLINs and IRS, may not be necessary in the future. With this in mind, the current intervention budget could eventually be budgeted to fund pgSIT, potentially covering 50% to 100% of the facility’s annual cost. With this budgeting, there would not be a significant increase in cost to the long term financing of mosquito suppression in the region.

Whether pgSIT is capable of being the sole malaria intervention tool in the region is unknown, but this table focuses on additional costs saved, including a minimum value saved in VSL, sick days, and medical intervention with potential maximum savings if IRS and LLINs are no longer needed.

#### 2.2.1 Willingness-to-pay value per year

“Willingness-to-pay”(WTP) is a commonly cited means to determine the amount that a person is willing to pay to avoid acquiring a disease. This measure captures the risks of the disease, such as the risk of pain and suffering, the inability to work, disability, and death. Parsing these risks in the WTP metric is difficult, but we can estimate how much the populace would be willing to pay annually to fund a pgSIT system. The average WTP metric for malaria prevention and treatment using current methods is 15.13 in 2022 USD (Trapero-Bertran et al. 2013). As current methods are unable to achieve malaria elimination in many regions, it is foreseeable that the WTP could be increased for this intervention. Using the current population of the URR and applying the expected population growth rate, the annual WTP for malaria prevention can be predicted. There are similar studies that look at The Gambia population’s willingness to pay for a national health insurance system(Njie et al. 2023). This study includes all medical interventions and puts the WTP at 23.27 USD on average, which corroborates the WTP for malaria interventions as it is a significant fraction of the total medical intervention cost.

The WTP estimate could also be used as a rough approximation of the value assigned by the local populace to malaria prevention interventions or as an individual cost contribution estimate for pgSIT if the citizens of the URR contributed financially to the facility as an alternative malaria intervention. This calculation shows that this project is priced to support interest from the local population (**Table 39**). It should be noted that this WTP is based on the willingness to pay for LLIN, IRS interventions, and treatments that will reduce the individual risk of infection and treat the disease. It is foreseeable with this intervention being more effective that the WTP for GE sterile mosquitoes would be greater than the WTP we are using. Importantly, the WTP should not be expected to fund the development of this project. The WTP money is an estimate of what the affected population would pay to prevent malaria with current methods and may not be readily taxable for The Gambia to fund this effort. This WTP price suggests that the local populace would be willing to be taxed up to 15.13 USD to prevent malaria, which could indefinitely support a government funded program. The annual cost of maintaining and running the facility is 5-8% of the WTP value, which is a small fraction of that value. This percentage will consistently decrease as the population of the URR increases, and more people invest in this intervention, making the total Upper River WTP increase. These local contributions facilitate the local sustainability of the project and would, therefore, along with any export income, limit the need for indefinite funding from external funding sources. Eventually, the project would be transferred to the local government, which would be responsible for providing the funds to support the intervention. The WTP value indicated that local financial support is likely in the long term making this project sustainable economically in the URR of The Gambia.

#### 2.2.2 Additional economic benefits

There are additional less quantifiable economic benefits of implementing pgSIT in the URR. Building the facility within The Gambia will have economic and security benefits for the country. All of the WHO approved LLINs and IRS insecticides are produced outside of The Gambia. LLINs are all produced in East Asia, Europe, and Tanzania, and all IRS chemicals are produced outside of Africa except for one facility in South Africa (Unicef and Others 2020). The pgSIT approach provides an additional financial benefit to The Gambia by providing investment, development, and financial input into the local economy that would typically be directed to foreign countries. There would be an initial investment in the necessary infrastructure as well as annual investment of 60,000 USD to the economy and workforce of The Gambia annually. In Tanzania, a similar benefit was seen when building a local factory to produce LLINs(Masum et al. 2010). There are also additional security and stability benefits that will be discussed in the discussion section (**Section 2.4.1.2 Safety benefits pgSIT**). Utilizing these facilities to produce pgSIT for export can supplement the income of this facility.

#### 2.2.3 Summary of economic benefits

The implementation of the pgSIT program has substantial economic benefits by the value of statistical life saved, sick days prevented metrics, as well as other potential advantages. It is estimated that a pgSIT program would save approximately 91 million USD in human life per year (VSL) once the intervention is functioning at full effect (**Table 31**). The development of a pgSIT-based malaria control program requires an initial investment ranging from 6 to 11.5million USD (**Tables 25 and 26**). These funds are utilized to establish the necessary infrastructure, acquire equipment and resources, and conduct research and development activities. The annual operational costs of the program range from approximately 315,000 - 318,000 USD (**Table 27**), covering expenses such as staffing, maintenance, supplies, and administrative requirements.

The first year of the deployment of the intervention is projected to save less compared to the following years as the model is accounting for release late in the second year. Once this program is functioning at full capacity, the facility will pay off the initial investment in the first year. The program’s positive impact on health outcomes and productivity led to a reduction in healthcare costs, increased workforce participation, and improved overall well-being. These factors contribute to significant financial savings, highlighting the cost-effectiveness and positive return on investment (ROI) of the pgSIT malaria prevention program.

Furthermore, since malaria is both the cause and consequence of poverty, it has been reported that investors interested in building factories are hesitant to do so with the current malaria climate (Sachs and Malaney 2002). Eradicating malaria would make this investment more feasible, beginning the process of raising the living and working conditions for those in the area.

Considering the program’s economic benefits, initial development costs, annual operational expenses, and the substantial savings expected in the first year alone, it becomes evident that the pgSIT program is cost effective. The potential for continued savings in subsequent years reinforces the long-term value and sustainability of the program, making it a worthwhile investment in both human lives and economic prosperity.

Notably, as this is a novel intervention, this is a preliminary model-derived estimate of disease prevention and lives saved. Field and safety trials will be conducted that will improve our estimates of production capabilities and the efficacy of mosquito population supression. The valuation of these benefits is performed to better compare this investment’s net benefit to other philanthropic investments.

#### 2.2.4 The unaccounted economic benefits derived from public health interventions

Public health experts agree that eliminating or drastically reducing child mortality is key for economic development. This value may be captured in part or whole by the predicted increase in GDP associated with preventing malaria in over 10% of the population, but there may be additional unaccounted value. By eliminating childhood mortality - primarily caused by malaria - women are likely to have fewer children and more likely to invest in these children and their own education. With this shift, the populace becomes better educated and therefore become higher skilled labor for investors looking to build facilities in the region. With new industries come new markets, increased wages, buying power, and ultimately improved quality of life. Eventually, this nationalized development can elevate the populace out of poverty. However, this potential progression is currently being held back by malaria. We, therefore, want to emphasize that the economic impacts of malaria eradication extend far beyond those direct and quantifiable calculations outlined here and extend into the possibility of opening up an entire nation to unprecedented economic development.

#### 2.2.5 Internal rate of return, net present value, and tracking expected return on investment

The internal rate of return (IRR) and net present value (NPV), along with timelines for the ROI also provide metrics to understand the costs and benefits of the project. The formula for calculating NPV is:

NPV = summation = R*_t_*/(1+*i)^t^*

NPV = net present value

R*_t_* = net cash flow at time

*t i* = discount rate

*t* = time of the cash flow

This equation tracks the investment into the facility and the value gained by the intervention at various times and compares this to the discount rate or bank rate which is the interest rate associated with bank loans. In other investments, where the profit is tangible finances, this is important to determine when an investment would be paid off.

The IRR is a metric used to determine the profitability of an investment. The NPV is set to 0 over the timespan of interest, and the discount rate is varied to allow NPV to equal 0 over that time period. For this, we assume an initial investment of either 6 or 11.5million USD and an annual facility cost of approximately 315,000 USDexpected. Nonetheless, we also wanted to account for the costs of more extensive trials and doubling the number sex sorting machines if we later find these are necessary. We then include these estimates in the various metrics for measuring the value saved:

1. VSL with all associated sickness costs. (**Table 42**)
2. QALYs with all associated sickness costs (High and Medium Estimates). (**Table 43**)
3. WTP (**Table 44**)

**Table 43:**
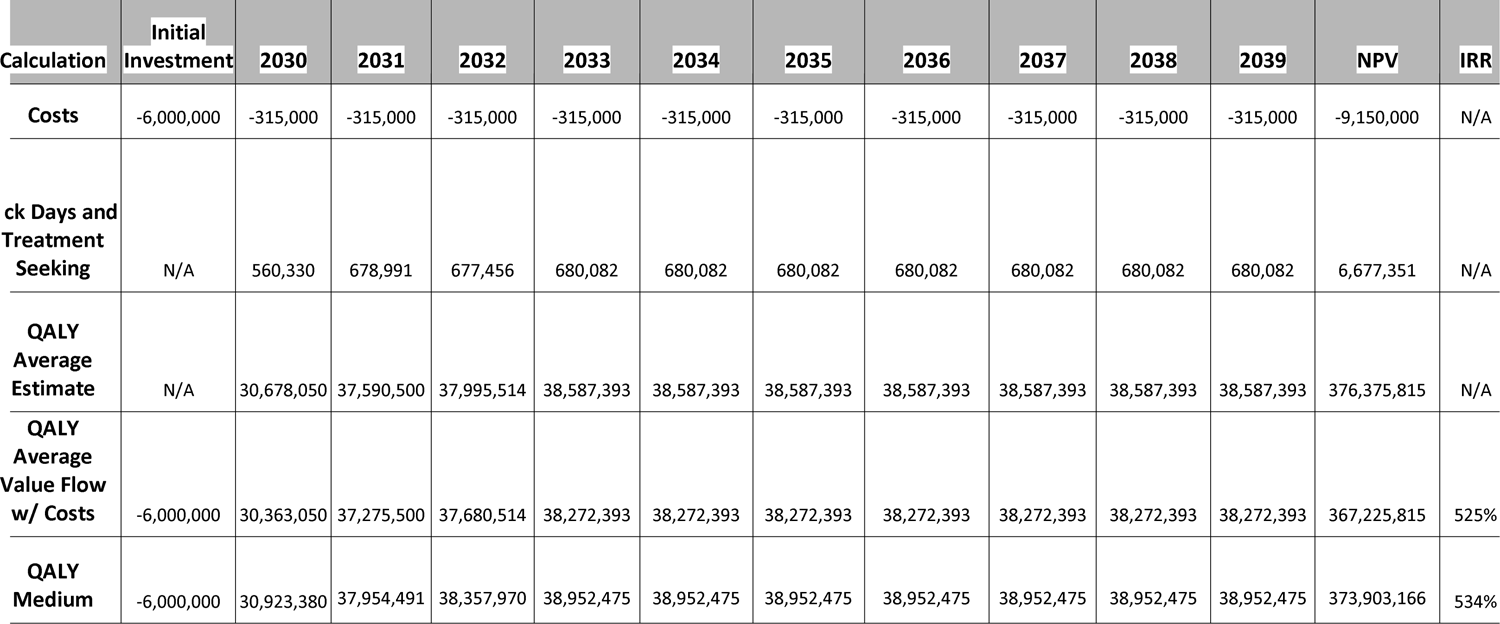
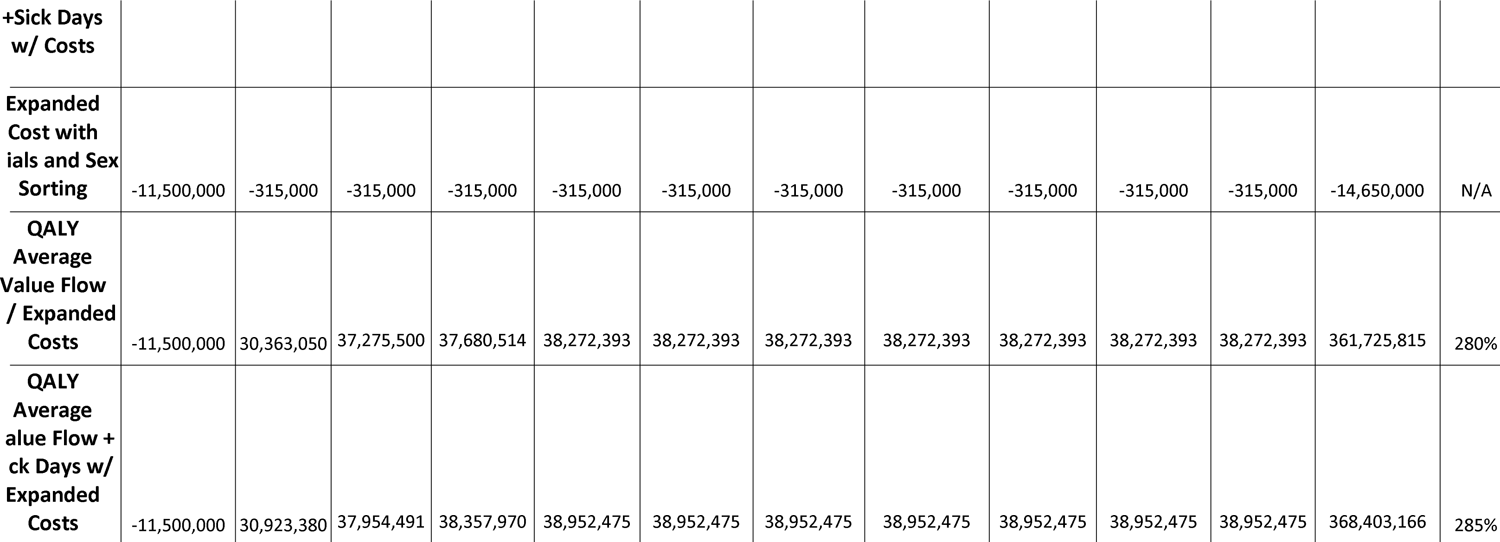
Net present value over 10 years and internal rate of return: QALY average estimate.

**Table 44:** Net present value over 10 years and internal rate of return: willingness-to-pay (WTP)

| Calculation | Initial<br>Investment | 2030 | 2031 | 2032 | 2033 | 2034 | 2035 | 2036 | 2037 | 2038 | 2039 | NPV or<br>Total | IRR |
| --- | --- | --- | --- | --- | --- | --- | --- | --- | --- | --- | --- | --- | --- |
| Costs | -6,000,000 | -315,000 | -315,000 | -315,000 | -315,000 | -315,000 | -315,000 | -315,000 | -315,000 | -315,000 | -315,000 | -9,150,000 | N/A |
| **WTP** | N/A | 5,541,710 | 5,681,360 | 5,824,536 | 5,971,312 | 6,121,795 | 6,276,060 | 6,434,214 | 6,596,362 | 6,762,596 | 6,933,005 | 62,142,950 | N/A |
| **WTP Value**<br>**Flow w/ Costs** | -6,000,000 | 5,226,710 | 5,366,360 | 5,509,536 | 5,656,312 | 5,806,795 | 5,961,060 | 6,119,214 | 6,281,362 | 6,447,596 | 6,618,005 | 52,992,950 | 90% |
| **WTP Value**<br>**Flow w/**<br>**Expanded**<br>**Costs** | -11,500,000 | 5,226,710 | 5,366,360 | 5,509,536 | 5,656,312 | 5,806,795 | 5,961,060 | 6,119,214 | 6,281,362 | 6,447,596 | 6,618,005 | 47,492,950 | 47% |

Notably, there are additional benefits that are not accounted for in these calculations, but these should include the main expected benefits. Each of these metrics is beneficial for assessing the cost effectiveness of the intervention. VSL is useful as an age agnostic method to calculate the value of life, although the value is significantly higher than many of the other methods. QALY accounts for age and has closer to a median benefit estimate when compared to the other interventions. WTP is mostly useful as a metric to estimate The Gambia’s ability to fund this project when management and funding are turned over to them. WTP is different from VSL or QALY as it is not calculating the value saved but the willingness to prevent harm.

The cost estimates will be subtracted from each of these groups to obtain the net value gained each year. Cost estimates are based on the highest estimates, with the minimum and absolute maximum number of sorting machines and simple and exhaustive field trials and monitoring. Additionally, VSL and QALY can each be added to the combined value of sick days and treatment seeking saved. VSL and QALY only account for the value of life saved, so adding the value of the cases avoided (sick days and treatment seeking) can give a more complete assessment of the total value saved without overlap of the values. Additionally, there are other benefits that could be added to this, but these benefits may not be fully realized with this system.

The net value estimated by each of these metrics is a net gain after ten years, with most net gains within the first couple intervention years, with the exception of WTP. The WTP value is both an underestimate of the actual WTP and an incomplete metric for the value of the intervention. Tracking of the predicted net costs is shown in **Table 44**. GDP is also included in this comparison to track the net predicted benefit. This exercise is not an extensive economic analysis of all potential outcomes though, and only compares the value gained to the most costly estimates for the intervention. If the intervention is less expensive than these estimates, which is the expectation, the intervention will be more cost effective.

These metrics all show a significant gain of value in terms of IRR over the ten years of implementation. All relevant calculations show value gains in the 100 millions, with the annual cost accounting for a small fraction of the value gained each year. The IRR similarly shows rates well above what would be desired for investment, with WTP being the lowest. The IRR ranges from 47%-1232%. WTP is likely significantly lower for several reasons, namely the WTP is adapted from current intervention options rather than pgSIT in particular. Since pgSIT has the potential to be a highly effective intervention, the WTP would likely be increased for pgSIT. WTP also does not capture the value saved from disease prevented and death prevented, making this an incomplete way to measure the value of intervention. Despite these limitations, this WTP is still shown to be cost effective.

The costs used are 6 or 11.5million USD initially and approximately 315,000 annually, which are the highest expected costs rounded up. The expanded costs capture increased trial costs increasing the initial cost to 11.5million USD. Calculations are done with both the standard and expanded cost estimates. The VSL and associated sickness costs are modeled out to 2034 as the prevention of disease and death is predicted to that point. We assume that years after that will be best reflected by the suppression seen in 2032-2034 when suppression is at full effect, so these values are averaged and projected out to year 2039. The VSL per year has the annual cost subtracted from it per year to get the VSL value flow and predict the IRR. For completeness, the cost of sick days and treatment seeking is added to the VSL value flow to obtain the NPV per year and predict the IRR. IRR was calculated using Excel Spreadsheet’s IRR function.

The costs used are 6 to 11.5million USD initially and 315,000 annually, which are the highest expected costs rounded up. Calculations are done with both the standard and expanded initial cost estimates. The QALY and associated sickness costs are modeled out to 2034 as the prevention of disease and death is predicted to that point. We assume that years after that will be best reflected by the suppression seen in 2032-2034 when suppression is at full effect, so these values are averaged and projected out to the year 2039. The QALY per year has the annual cost subtracted from it per year to get the QALY value flow and predict the IRR. For completeness, the cost of sick days and treatment seeking is added to the QALY value flow to estimate the NPV per year. The average estimated QALY was also used to predict the IRR. IRR was calculated using Excel Spreadsheet’s IRR function.

The costs used are 6-11.5 million USD initially and 315,000 annually, which are the highest expected costs rounded up. The expanded costs capture increased trial costs to 11.5million USD. Calculations are done with both the standard and expanded initial cost estimates. The WTP is calculated from similar WTP studies and the expected population growth for The Upper River Region. The annual WTP has the annual cost subtracted to obtain the WTP value flow and to predict IRR. The cost of sick days and treatment seeking is added to the WTP value flow to obtain a more complete NPV per year and to predict IRR. IRR was calculated using Excel Spreadsheet’s IRR function.

#### 2.4.1 Discussion

##### 2.4.1.1 Cost effectiveness summary

First and foremost, this cost feasibility study shows that investment in pgSIT will result in a highly cost-effective approach to preventing malaria-associated disease and death in The Gambia. Even when accounting for WTP, an estimate of the affected population’s willingness to pay, the facility would have a net gain within six years of operation. When accounting for the value of life, sick days, or GDP associated metrics, the facility is cost effective by the second year of operation, paying off the investment. It can likely be cost effective by the first year too, if the first year of releases is modified to effectively suppress the *A. gambiae* population during that first year of release. With the highest total VSL, sick days, medical intervention, and prevention costs saved, **pgSIT would save an estimated 91 million USD annually. The initial cost to implement this technology ranges from 6 to 11.5million USD, and the annual cost of about 315,000 USD**. Therefore, this approach is saving lives and health at least 13% of the value for statistical life and health benefits in The Gambia in the second year of operation and sub 1% annually thereafter. This approach is extremely cost-effective when considering the economic losses prevented by this technology. There is usually an expectation for public health interventions that it will take many years for the project to break even on the initial investment. This facility would not only pay for itself four times over in VSL in the first year of operation, it will continue to save life at a fraction of the statistical cost. Other metrics provide a more conservative estimate of the value saved, with QALY life savings being 32% of investment in that second year with life being saved at 0.8% of the QALY value saved annually. The higher percentage initially is due to the inclusion of the initial development costs. The GDP growth effect shows that this could have exponential benefits to the country, potentially accelerating its economic development, which will generate many other indirect benefits. Finally, WTP shows that the community in the URR may be willing to partially fund this intervention in the future, which strongly suggests this will be a sustainable project by the local government.

Second, pgSIT is highly cost effective when compared to current interventions. For the URR, LLINs, IRS, and medical treatment cost approximately 345,000 USD annually but excludes the value of volunteer labor that is difficult to account for. As cost effective as the current interventions are, these interventions have limitations, and the implementation of pgSIT has the potential to completely eliminate malaria in the region. The current methods are quite cost effective, but they are likely at their peak capability due to the limitations of low compliance rates, growing insecticide resistance, and the lack of outdoor interventions (Wubishet et al. 2021; Fru et al. 2021; Aweis et al. 2023; Kayedi et al. 2007). These issues make it highly unlikely that these approaches could eliminate malaria in these regions, even if the investment is increased. With the growing issue of insecticide resistance, current interventions will inevitably decrease effectiveness. Alternative insecticides in the development pipeline are often more costly and without proper management may also rapidly become ineffective for vector control due to insecticide resistance (Hemingway 2014).

Additionally, it is useful to compare the cost of saving disability adjusted life years to other public health technologies. Depending on how much research and development is included in the cost, this **intervention is expected to save DALY at 15 to 124 USD (2022) per life-year saved. This intervention prevents malaria infections at 13 to 113 USD per case**, costing less than 17% of GDP per capita to save a life year. Even though this intervention is additive, the cost per DALY is comparable to the current cost per DALY averted of current malaria interventions. Horton et al. 2017, summarizes estimates of the costs per life-year saved in 2012 USD for many malaria interventions, which we used to convert to 2022 USD, which is about 21 to 95 USD, quite comparable to our estimates for pgSIT(Horton 2017). Notably, the high estimates include research and development costs that may not be accounted for in other interventions. The pgSIT intervention outperforms other interventions, but initially pgSIT will be additive to current standard of care interventions. While it is possible that pgSIT could supplant current interventions in the future, large scale implementation is likely needed before this is known.

##### 2.4.1.2 Other unknowns for cost effectiveness

There are several unknowns that could increase the cost efficiency of pgSIT. The first unknown is the year-to-year impact of pgSIT on the mosquito population. It is assumed that there is some level of migration of mosquitoes, but the extent and range of these migrations are not well understood. *Anopheles gambiae* populations drop below detectable levels seasonally in the URR and in many other malaria endemic regions which leads some to speculate that during the dry season these mosquitoes are locally extinct and rely on long distance migration to repopulate these regions(Faiman et al. 2020; Sanogo et al. 2021; Huestis et al. 2019). Whether these mosquitoes have small local populations maintained during the dry season or they rely on long distance migration, both complement pgSIT mosquito suppression strategies. If *A. gambiae* persists in small undetectable populations during the dry season, a year-to-year carryover effect that eliminates this following dry season population could allow for the production at this facility to be retargeted to other regions (Baber et al. 2010). This result could potentially allow this facility to slowly facilitate the local extinction of *A. gambiae* throughout the region, significantly expanding the effectiveness of this facility. On the other hand, if *A.gambiae* are reliant on long distance migrations to repopulate regions after the dry season, there should be an initial time when migration has only brought a few mosquitoes to the area. Therefore, if pgSIT releases are also timed before the population increases substantially, a lower amount of pgSIT mosquitoes will be required to treat the region, and consequently, there will be enough individuals available to release pgSIT into larger areas, or other regions. With long-distance migration being a dominant theory for *A. gambiae* repopulation of seasonally populated regions, this early release approach has been evaluated in our models and will likely be used in the field. Either way, there may be the potential to cause the extinction of these population reservoirs during the dry season to have a greater effect on the mosquito population. It is suspected that riparian habits by rivers are a dry season reservoir of *A. gambiae*. Targeting dry season releases to these sites could potentially lead to greater suppression effects at an even cheaper cost than laid out in this report.

The migration range is, therefore, important for planning large scale suppression of *A. gambiae.* There is a wide range of predicted migration distances from below 50 kilometers to over 100 kilometers. If the migration distance is below 50 kilometers, it is likely that large-scale suppression of the URR will have core regions that are protected indefinitely as long as the regions surrounding maintain reduced population levels. If this migration is over 100 kilometers or has no clear upper limit, regions will unlikely be able to achieve permanent suppression until pgSIT is spread to additional regions.

Another relevant unknown previously discussed in the economic benefits section is the potential to switch to a pgSIT-only approach to malaria prevention and, therefore, save costs associated with current prevention activities (LLINs, IRS), and medical interventions. If pgSIT achieves local extinction of *A. gambiae* as predicted in regions with only *A. gambiae* malaria vectors, it may be possible to rely on pgSIT alone to prevent malaria transmission. While this will require monitoring before withdrawing, these other preventative interventions, if large population reduction is achievable, will allow pgSIT to become the only approach required for malaria prevention in the region.

##### 2.4.1.3 Safety benefits pgSIT

pgSIT will have multiple biological safety benefits, not only compared to more invasive interventions, like gene drives, but also when compared to current interventions. First, pgSIT is species-specific or species-complex-specific and will only target and suppress *A. gambiae* in particular or the complexes of subspecies. There are many malaria vectors in the *A. gambiae* species complex (e.g. *Anopheles coluzzii* and *Anopheles arabiensis*), and *A. gambiae* is currently undergoing a speciation event, so multiple subspecies may be targetable by the release of *A. gambiae* pgSIT. Since many of these species and subspecies also spread disease to humans, it would be ideal to suppress these as well, but whether pgSIT can accomplish this requires further exploration. Many current control methods use insecticides, which have off-target effects on beneficial insects and local biodiversity. Additionally, as mosquitoes become increasingly resistant to insecticides and require more intensive application of insecticides to maintain the same effect, these negative off-target effects will only increase.

The pgSIT technology is also a dead end for synthetic transgenes, whereby the sterile males do not produce viable offspring, and therefore, these synthetic genes are rapidly eliminated from the population. If there is an unpredicted issue with the genetics of the line, simply halting their release will rapidly eliminate their genes from the populations. If a gene drive technology or another technology that spreads and persists in the environment is released in the area in the future, there will also be no pgSIT genes in the population to potentially complicate the introduction of these technologies.

##### 2.4.1.4 Investment benefits and biomedical supply chain security

The facility in The Gambia has additional economic and biomedical security benefits. Producing pgSIT mosquitoes in The Gambia reinvests the money sent to foreign nations to produce anti-malarial technologies into the local economy. This project would provide 60,000 USD in wages to the local economy of The Gambia that would normally support facilities in foreign countries. A local pgSIT facility, therefore, can have many downstream benefits to support the development of new industries in The Gambia. Other African nations produce their own anti-malarial interventions, for example, the AtoZ insecticide treated net manufacturing facility in Tanzania(Masum et al. 2010), which provides over 8,000 jobs to Tanzanians. This production facility also gives Tanzania a more secure biomedical supply chain, as it relies on fewer imports to produce its malaria prevention technologies.

Reliable and sustainable malaria programs are important to achieving malaria elimination. During the recent coronavirus pandemic, malaria prevention programs were disrupted by lockdowns and supply chain issues. Consequently, malaria deaths increased by 12% during this period, likely due to the disruption of these services (“More Malaria Cases and Deaths in 2020 Linked to COVID-19 Disruptions” n.d.). Future events that disrupt biomedical supply chains are likely and will cause similar disruptions in malaria prevention programs. A pgSIT facility in The Gambia should be able to source most of its supplies for mosquito production from within the country, making pgSIT more resilient to external supply chain disruptions.

##### 2.4.1.5 Alternative production at the pgSIT facility

The development of a pgSIT *A. gambiae* mass production facility would represent a significant investment of capital that would only be ramping up and running at full capacity for approximately 20 weeks. This production schedule leaves an additional 34 weeks of the year that can be leveraged for other work. A pgSIT mass production facility can be utilized for a couple of key purposes, including the production of other mosquito species, or other control technologies. In the future, the facility could be used for the production of more invasive interventions, such as gene drives. In the short term, this could consist of mass rearing the pgSIT technologies to control the dengue vector, *Aedes aegypti*, which has eggs that can be stored for months and stockpiled before release. While dengue is not a major concern for The Gambia currently, it is an emerging disease in many parts of Africa, including neighboring Senegal, which has had multiple outbreaks since 2017 (Dieng et al. 2021). This facility could produce, stockpile, and release these *A. aegypti* pgSIT mosquitoes to curtail future dengue outbreaks in the region. If not needed in The Gambia, or Senegal, the dried Aedes eggs can be exported to other countries, and the revenue from this production can offset some of the facility costs.

The facility could also support the development of pgSIT technologies in other malaria vector species, or other research activities. Space and equipment in the facility could be rented to researchers and local universities, such as the International University in Banjul. These activities would not only support the development of additional technologies to support mosquito control but can also offset some facility maintenance costs.

Finally, this facility creates the critical infrastructure necessary for alternative genetic suppression technologies in the future, such as the more invasive gene drive suppression technologies, or any future genetic suppression technology. The pgSIT approach is an effective and controllable genetic suppression technology as it produces sterile males that prevent the persistence of transgenes in the population. The current approach is a safe version of genetic suppression as it does not introduce any transgenes into the wild population. Large-scale releases of this technology into the Upper River Region will likely have the ability to cause localized extinction or near extinction of *A. gambiae*. Moreover, given that many of the other malaria vectors can mate with *A. gambiae* males - it is possible that pgSIT may also suppress populations of these related species, thereby providing unprecedented multi-species protection.

These studies are a key step to understanding the environmental impacts of pgSIT and localized *A. gambiae* extinction in the region and are important for regulatory approval of genetic technologies in the field. Once the impact of pgSIT and its greater ecological effects is better understood, this may pave the way for more invasive technologies to be utilized in The Gambia and beyond or may simply show that technologies such as pgSIT are sufficient for sustainable malaria control. Many more invasive interventions, such as gene drives, have technological limitations, and safety concerns, and consequently, their regulatory pathway is less certain, so it may be many years before they are available to malaria control programs, if ever. Therefore, this facility will provide a valuable intervention that can support the deployment of any future mosquito genetic suppression technologies in the region.

##### 2.4.1.6 Risks, challenges, and risk mitigation

An *A. gambiae* pgSIT mass rearing facility of this scale has not been developed yet. While many mass rearing issues can be addressed by increasing the facility size, there are several notable concerns and solutions that warrant further discussion.

The proposed mass sex separation technologies have been extensively discussed with their manufacturer and technical teams, but these technologies have not been directly used in a mass rearing facility. Ideally, initial tests of these technologies will begin during the pgSIT field trials, which will provide opportunities to troubleshoot the scaling of this technology. During the smaller scale trials, the production needs will be low enough that manual sex sorting can be used, while automated sex sorting requires expensive devices that will not be necessary at the lower release rate, but will provide the opportunity to test the capabilities of the sorting options. While the sorting rates indicated by the supplier, Union Biometrica, should allow for the number of machines described in the COPAS section, these machines have not been used to sort mosquitoes for a mass rearing facility. In particular, the daily and weekly run-time capabilities of these machines has not been fully explored. However, further testing can be done by Union Biometrica to confirm these capabilities, and the field studies will allow us to test whether these sex sorting technologies are capable of running constantly under Active Phase conditions. Fortunately, the daily production requirement is well below the expected capability of the COPAS FP 500 and we have already budgeted two sorting machines in case of failure with the primary one.By evaluating these sex sorting technologies during the early testing stages, we can better estimate the machines required for this large scale facility.

Another potential solution to address the problem of sex sorting is to pursue collaboration with companies that have proprietary sex sorting technologies, such as Verily Life Sciences (South San Francisco, CA). While they have not made this technology commercially available, this may be a future possibility. This technology would act similarly to the Senecio Robotics adult stage sex sorting technology. This solution likely assumes a future collaboration with Verily Life Sciences.

Public support and regulatory approval is another important but high risk aspect of the project. With other genetic technologies being released in Brazil, Burkina Faso, Cayman Islands, Panama, India, Malaysia, and the United States, a path to regulatory approval is more certain; however, The Gambia has not had a previous genetically engineered insect trial to date(Lacroix et al. 2012; Carvalho et al. 2015; Halliday 2017; Gayler-Thompson 2023; Halliday 2018; Pare Toe et al. 2021; Waltz 2021). Public engagement is important to begin early and often to ensure community concerns can be discussed and addressed. Public communication and support are crucial for this case because this project relies on communities to manage pgSIT mosquito rearing on site. With this in mind, we have included a public outreach budget similar to other campaigns utilized in The Gambia for other malaria interventions. This outreach will rely on the expertise of our colleagues at the MRC Unit, who have managed previous large scale malaria prevention engagement activities in the region. The current cost of public outreach and training in the URR is approximately 9,000 USD annually, but due to the novelty of pgSIT, we plan up to two-fold more for these activities.

##### 2.4.1.7 Project timeline and development plan

In addition to project costs, it is important to determine the technological development timeline. The technology is likely 8 to 9 years from large-scale deployment, assuming funding and regulatory approvals adhere to predicted schedules (**Fig. 18**). If the facility is built concurrent to the field trials, the timeline should decrease to 6 or 7 years, but if funding is contingent upon the completion of the previous stage, this may not be realistic. The first stage of this project is currently underway, whereby we are developing the pgSIT 2.0 strain to support fluorescent marker-based sex sorting of the Cas9 and gRNA lines. This stage has minimal risk and relies on engineering the lines and confirming their phenotypes. This development and testing will take 1 to 2 years with the expectation that the non-fluorescent sex sorter line, pgSIT 1.0, can be tested in a cage trial in The Gambia, while pgSIT 2.0 testing is completed. These 1 to 2 years will also provide time to obtain approvals for field release. After the outdoor cage trials in The Gambia, the next step is a small-scale field release to determine the safety of this system (**Fig. 4**). These initial releases will be primarily geared towards safety, so the pgSIT sterile males will be released at smaller quantities and frequencies than would be needed for complete or large scale suppression. Depending on the results of the cage studies and subsequent updates to the predictive suppression models, these trials may be over just a single rainy season but will aim to inform larger scale trials the following year, assuming no unexpected regulatory or technical issues arise. In the larger scale field trials, safety testing and monitoring will continue, but the entomological impact, or ability to reduce the local *A. gambiae* population, and possibly the epidemiological impact, or ability to reduce malaria case incidence, will be key measurements of efficacy in these trials. Completing large scale trials provides the necessary data to plan and obtain approvals for larger scale releases in 6 or 7 years. The facility construction will then begin with an estimated two year completion timeline. If construction is started prior to the completion of the trials, releases can occur on the earlier timeline. The first year of release from the facility will have a significant number of foreign specialists tasked with training local staff in the facility. The following years will be led by the local staff in The Gambia. Finally, once the large-scale field releases and monitoring are established for multiple years, these results can be used to optimize releases, and the facility can be utilized for off-season production of other genetic suppression technologies for vector control.

**Fig. 18:**
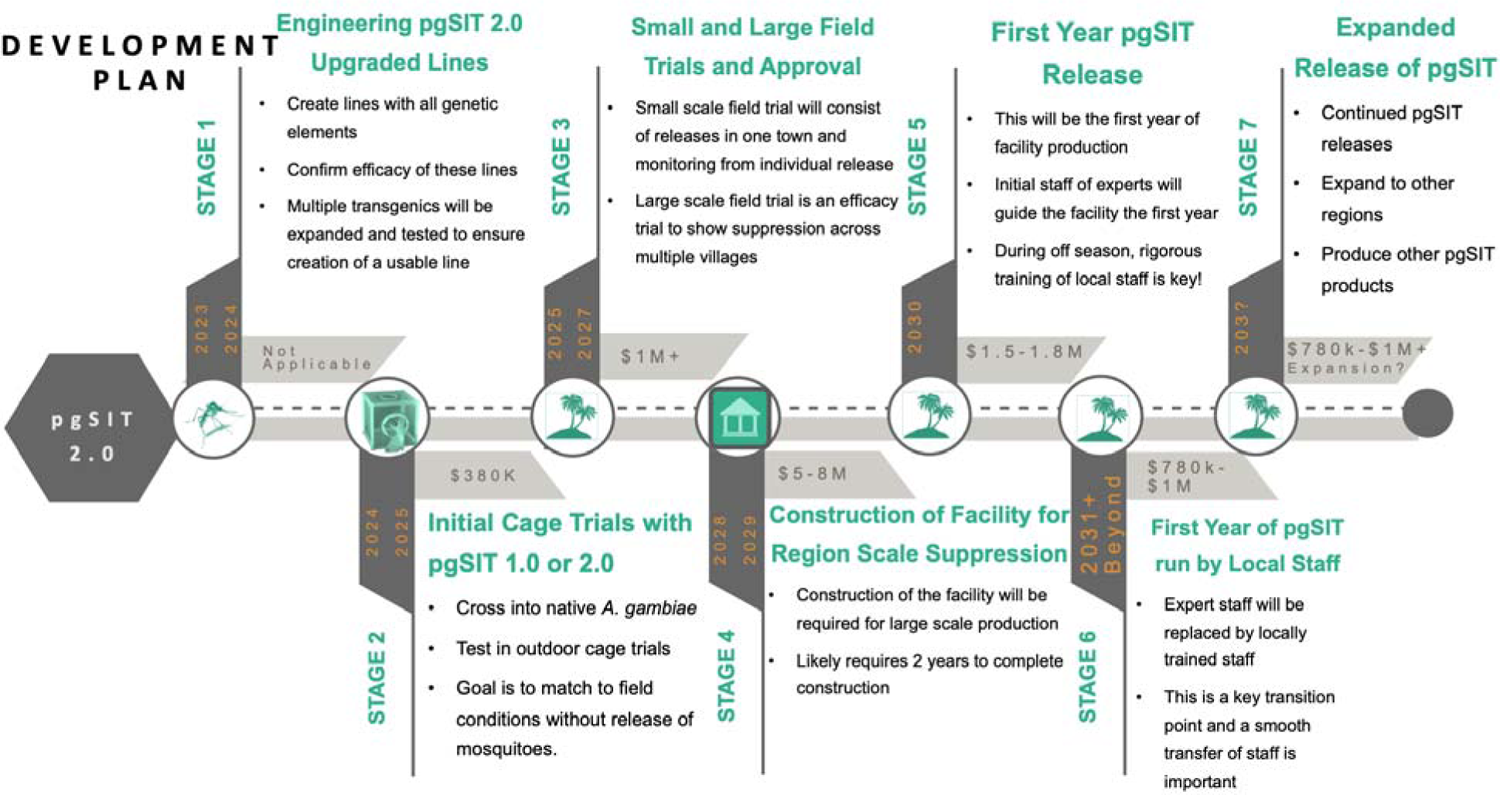
Development plan for pgSIT testing and large scale deployment.

Each stage of the development plan includes a brief description of activities, timeline, and cost estimates. This plan could be accelerated if construction of the facility begins earlier, or a modular design is used to simplify facility expansion, but the current plan minimizes investment risk by assuming the completion of each stage before continuing to the next.

## Methods

### Mathematical modeling

We used the MGDrivE 3 framework (Mondal, C, and Marshall 2023) to simulate releases of pgSIT *A. gambiae* mosquitoes to suppress malaria in the Upper River region of The Gambia. MGDrivE 3 is a modular framework for simulating releases of genetic control systems in spatially-structured mosquito populations which includes modules for: i) inheritance (i.e., the dynamics of the pgSIT system), ii) life history (i.e., the development of mosquitoes from egg to larva to pupa to adult, including species-specific bionomic parameters), iii) spatial structure (i.e., the distribution of mosquitoes in a metapopulation network), and iv) epidemiology (i.e., the reciprocal transmission of malaria parasites between mosquitoes and humans). We simulated the inheritance pattern of the pgSIT system within the inheritance module of MGDrivE (Sánchez C et al. 2020), the life history of *A. gambiae* using standard bionomic parameters and seasonality in larval carrying capacity driven by rainfall data for The Gambia, ignored spatial structure, and calibrated malaria transmission to incidence data from the Upper River region of The Gambia.

### Model of pgSIT inheritance dynamics

The inheritance pattern of the pgSIT system was modeled within the inheritance module of MGDrivE (Sánchez C et al. 2020). Based on laboratory data (Smidler, Apte, et al. 2023)(M. Li et al. 2021), we assumed the pgSIT system in *A. gambiae* would induce complete male sterility and female inviability, with inviability being manifest at hatching. We assumed that pgSIT eggs would be introduced into the environment in cups with sufficient water volume and larval resources such that larval mortality would be density-independent. Survival of eggs released in cups was therefore determined by expected juvenile life stage durations and their daily mortality rates (**Table S1**) - an egg stage of 3 days with a daily survival probability of 95%, a larval stage of 7 days with a daily survival probability of 85%, and a pupal stage of 1 day with a daily survival probability of 95%, leading to a viable emergence rate of 26% for male eggs. Offspring of pgSIT sterile males are unviable at the egg stage. The pgSIT cube also allows impacts of the construct on adult lifespan and male mating competitiveness to be incorporated into the model. Following the cage trial results, we assumed a 50% reduction in pgSIT male mating competitiveness (Smidler, Apte, et al. 2023). To be conservative, we also assumed a 25% reduction in pgSIT male lifespan compared to wild-type males, despite no reductions in lifespan being observed in this work (M. Li et al. 2021; Kandul et al. 2018).

### Model of A. gambiae life history

The MGDrivE 3 framework (Mondal, C, and Marshall 2023) models the development of mosquitoes from egg to larva to pupa to adult with overlapping generations, larval mortality increasing with larval density (White et al. 2011), and a mating structure in which females retain the genetic material of the adult male with whom they mate for the duration of their adult lifespan. Species-specific bionomic parameters for *A. gambiae* are listed in **Table S1**. The life history module of MGDrivE 2 permits life history parameters to change with time. Given the dependence of *A. gambiae* populations on recent rainfall to provide egg-laying and juvenile development sites, we utilize rainfall data from the URR of The Gambia (https://www.chc.ucsb.edu/data/chirps) to calibrate a seasonal time-series for larval carrying capacity, which is a major driver for their population dynamics. To smooth the seasonal profile of the raw rainfall data, we leverage a Fourier analysis-based approach that involves fitting a mixture of sinusoids to the raw data (https://github.com/mrc-ide/umbrella) to depict the general seasonal trends without the level of daily detail that is not replicated from year to year. The fitted seasonality profile for the Upper River is shown in **Fig. S1**.

**Fig. S1.**
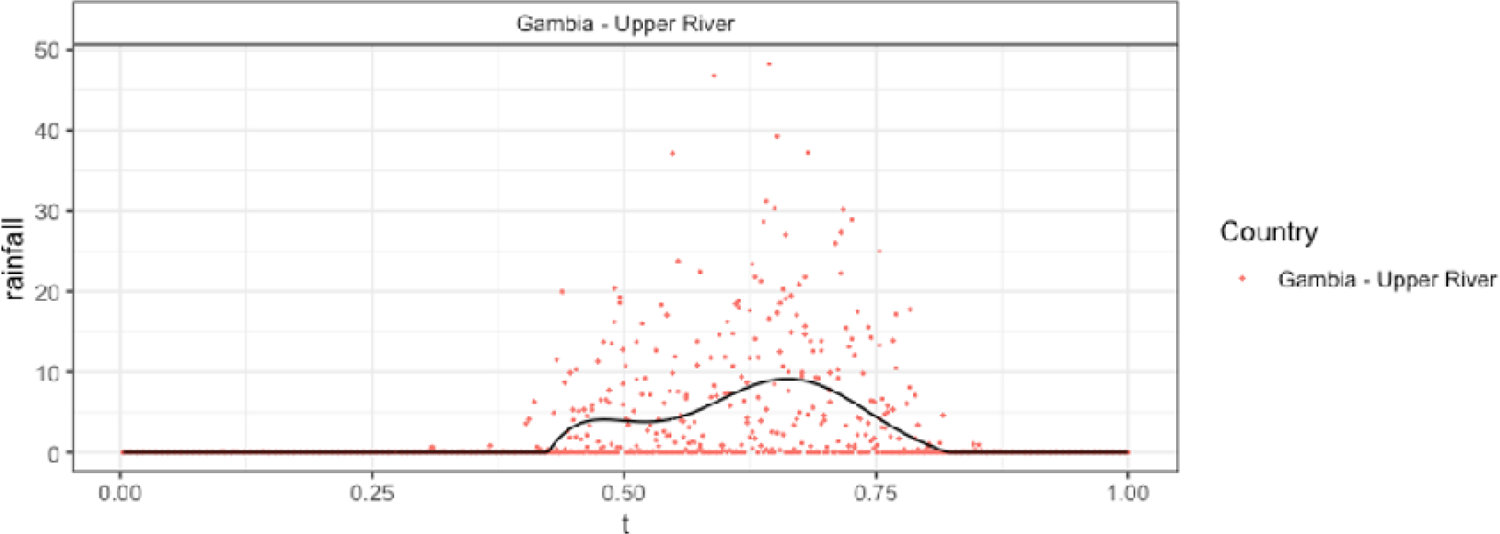
Seasonal rainfall profile for The Gambia. Actual daily rainfall in millimeters is depicted as red dots, and a mixture of sinusoids fitted using the Umbrella package in R (https://github.com/mrc-ide/umbrella) is depicted as a black line.

### Model of malaria transmission

To model epidemiological outcomes associated with pgSIT mosquito releases, we linked the MGDrivE 3 framework (Mondal, C, and Marshall 2023) to the ICL malaria model (Griffin et al. 2016, 2010). Linking the two models was achieved by allowing forces of infection (i.e., the probability of infection from mosquito-to-human and human-to-mosquito per individual per unit time) to be exchanged between the two models. We chose the ICL malaria model as it provides a validated and parsimonious framework to describe transmission in the human populations, including important features such as acquired and maternal immunity, symptomatic and asymptomatic infection, superinfection, age structure, biting heterogeneity, and antimalarial drug therapy and prophylaxis. The model describes humans moving through various infection states (susceptible, asymptomatic, treated, clinical disease, untreated disease, and protected) and includes various forms of immunity (maternal, acquired, and pre-erythrocytic), which reduce the probability of severe and clinical disease (**Fig. 2**). The model is age-structured, allowing outcomes to be estimated for chosen age groups. This is important, as pediatric cases of malaria tend to be the most severe.

The ICL malaria model was calibrated using malaria prevalence data from a randomized-control trial of mass drug intervention in the URR (Dabira et al. 2022). The study found a baseline malaria prevalence in the URR of ∼18% at the beginning of the rainy season, which we aligned with the simulation output. Furthermore, entomological data from the URR (Soumare et al. 2022) suggested vector breeding sites in this region are substantially more abundant in the rainy season than in the dry season, so we assumed that larval carrying capacity in the dry season was 10% that of the peak rainy season. The malaria model calibration took place in the context of model parameterization with existing levels of coverage of ACTs, LLINs, and IRS with insecticides, which modify the transition rates between human infectious states and life history parameters of the mosquito vector population, respectively. For the URR, data from the Malaria Atlas Project (Pfeffer et al. 2018) was used to parameterize an LLIN coverage of 55%, an IRS coverage of 52%, and 50% of symptomatic malaria cases treated by antimalarial drugs. The remaining parameters of the malaria transmission model were taken from the ICL implementation (Griffin et al. 2016, 2010), including a mortality rate of 21.5%, which represents the proportion of severe malaria cases that result in death and was parameterized according to hospital case data collected in Tanzania (Griffin et al. 2016).

### Intervention model

Weekly releases of pgSIT *A. gambiae* eggs were simulated from the beginning of the rainy season (June 1st), for a variable number of weeks and release sizes. Previous pgSIT modeling studies (M. Li et al. 2021; Kandul et al. 2018) suggested that local mosquito populations could potentially be eliminated by ∼10-24 consecutive releases of ∼40-400 pgSIT eggs per wild adult. We explored release schemes within and slightly below this range, focusing on schemes that required smaller weekly release sizes, as this was determined to be more cost-effective. Model outputs of interest to the cost-effectiveness analysis included clinical incidence over time, and mortality over time.

#### Anopheles gambiae mass rearing facility design

We used the IAEA *Anopheles* mass rearing protocol traditional SIT methods (FAO/IAEA 2017) as the basis for a mass rearing facility in this report. Additional features such as sex sorting and associated equipment, facility design, drone delivery technology, and similar were added as these were not included in the IAEA report. Some modifications were made to rearing racks and trays, larval feed, and other materials to find more cost effective and local alternatives to the ones suggested in the IAEA report. Facility size and scaling was estimated according to the planned work practices and the facility space and safety requirements.

#### Anopheles gambiae mass rearing facility cost assessment

The cost assessment was done by exploring numerous options for the facility design. When available, we obtained local cost estimates, and when necessary international prices were adjusted to account for currency conversion, importation, shipping, and other associated costs with outside sourcing.

**Table S1.** Model parameters describing pgSIT construct, *Anopheles gambiae* bionomics and malaria epidemiology for simulated releases in Upper River region, The Gambia.

| Parameter | Value | Reference |
| --- | --- | --- |
| **pgSIT construct:** |  |  |
| pgSIT male fertility | 0.0 | (Smidler, Apte, et al. 2023) |
| pgSIT female viability | 0.0 | (Smidler, Apte, et al. 2023) |
| pgSIT male mating competitiveness (relative to wild-type) | 0.5 | (Smidler, Apte, et al. 2023) |
| pgSIT lifespan (relative to wild-type) | 0.75 | (M. Li et al. 2021) |
| ***Anopheles gambiae* bionomics:** |  |  |
| Egg production per adult female (day<sup>-1</sup>) | 21 | (White et al. 2011) |
| Mean duration of egg stage (days) | 3 | (White et al. 2011) |
| Mean duration of larval stage (days) | 7 | (White et al. 2011) |
| Mean duration of pupa stage (days) | 1 | (White et al. 2011) |
| Egg stage mortality rate (day<sup>-1</sup>) | 0.05 | (Wu et al. 2021) |
| Larval stage mortality rate (day<sup>-1</sup>) | 0.15 | (Wu et al. 2021) |
| Pupal stage mortality rate (day<sup>-1</sup>) | 0.05 | (Wu et al. 2021) |
| Carrying capacity of environment for larvae | (time-varying) | (White et al. 2011; Bayoh and Lindsay 2003) |
| Mortality rate of adult mosquitoes (day<sup>-1</sup>) | 0.132 | (Wu et al. 2021) |
| **Malaria transmission (Upper River region, The Gambia):** |  |  |
| Malaria prevalence at peak of rainy season | 18% | (Dabira et al. 2022) |
| Proportion of larval breeding sites available in dry season | 10% | (Soumare et al. 2022) |
| LLIN coverage | 55% | (Pfeffer et al. 2018) |
| IRS coverage | 52% | (Pfeffer et al. 2018) |
| ACT drug coverage | 50% | (Pfeffer et al. 2018) |

**Table S2.** Blood feeding costs associated with alternative blood sourcing.

|  | **Total**<br>**Liters of**<br>**Blood Per**<br>**Year** | **Cost Per**<br>**Liter** | **Shipping**<br>**Cost** | **Storage**<br>**Cost**<br>**(Energy**<br>**and repair)** | **Up Front**<br>**Cost** |
| --- | --- | --- | --- | --- | --- |
| **Fresh**<br>**Blood: 14**<br>**Week High**<br>**Fecundity** | 1453 | <\$2.24 | N/A | N/A | N/A |
| **Fresh**<br>**Blood: 14**<br>**Week Low**<br>**Fecundity** | 2075 | <2.24 | N/A | N/A | N/A |
| **Frozen**<br>**Blood: 14**<br>**Week High**<br>**Fecundity** | 1453 | 2.24 | 6,000 | 1,800 | 18,000. |
| **Frozen**<br>**Blood: 14**<br>**Week Low**<br>**Fecundity** | 2075 | 2.24 | 6,000 | 1,800 | 18,000 |
| **Frozen**<br>**Blood: 14**<br>**Week High**<br>**Fecundity** | 1453 | 2.24 | 3,000 | 6,400 | 64,000 |
| **Frozen**<br>**Blood: 14**<br>**Week Low**<br>**Fecundity** | 2075 | 2.24 | 3,000 | 6,400 | 64,000 |

**Table S3.**
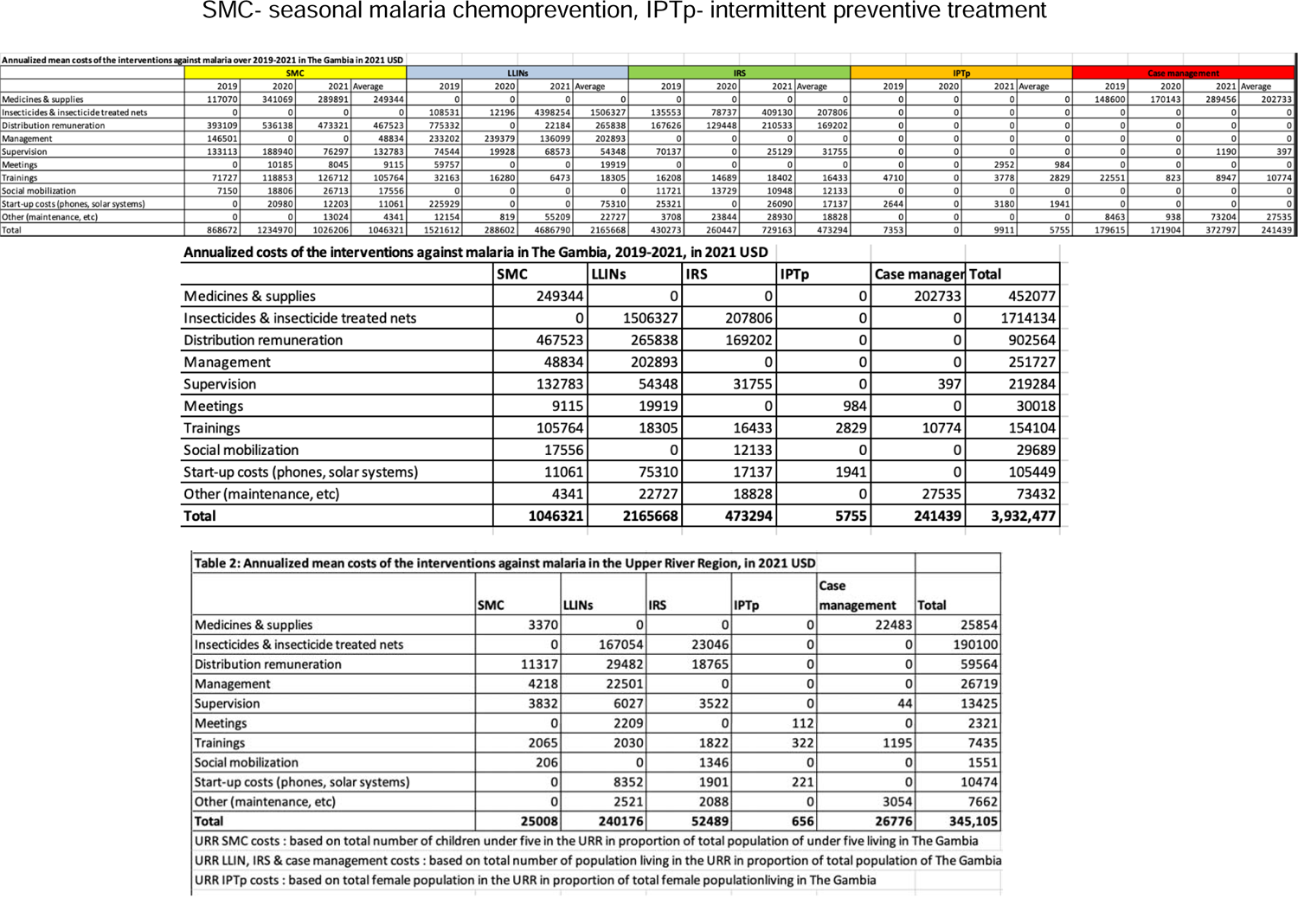
Current annualized investment cost for anti-malaria intervention in The Gambia, mean cost and URR specific costs. SMC-seasonal malaria chemoprevention, IPTp-intermittent preventive treatment.

**Table S4.**
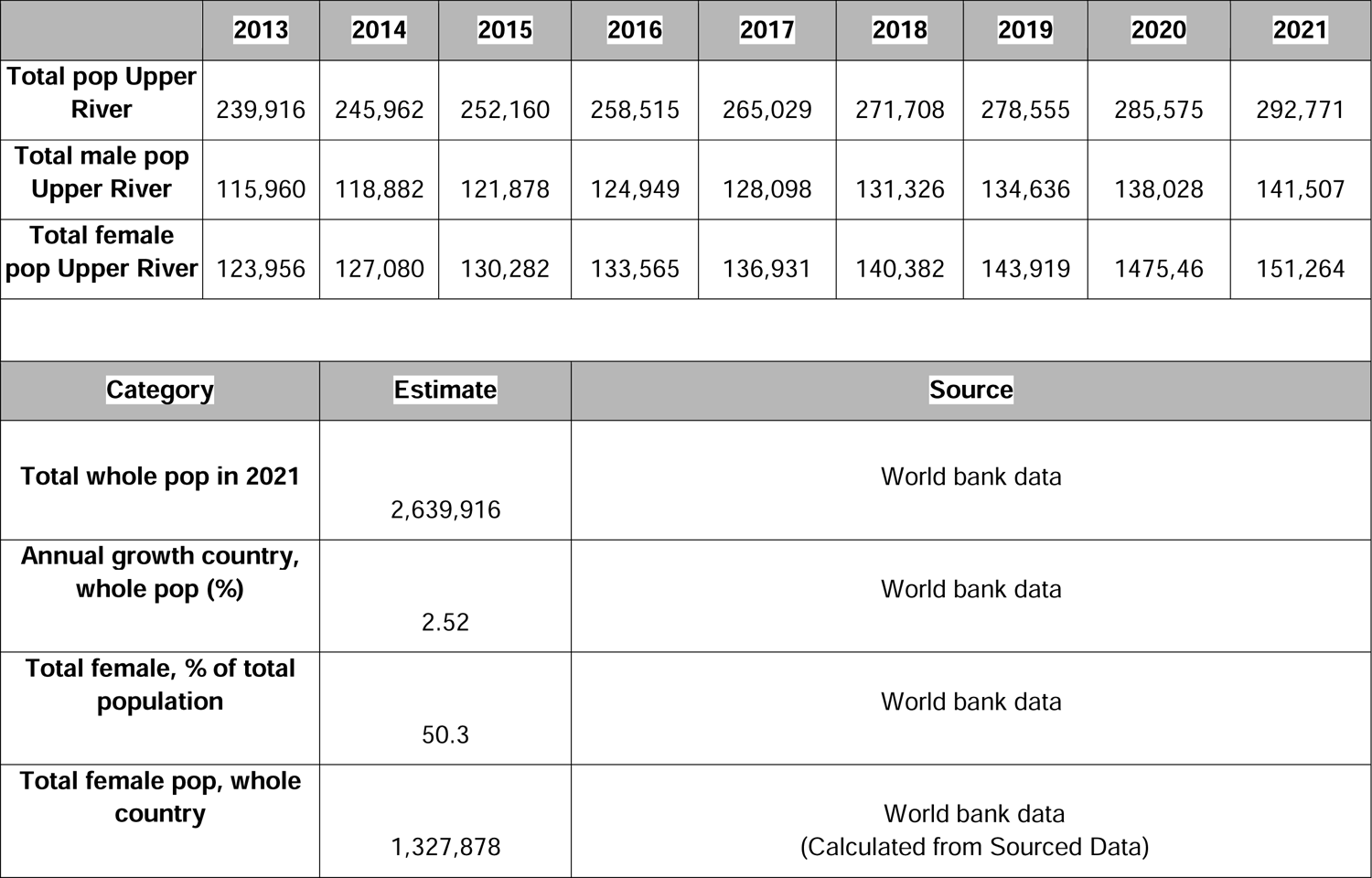
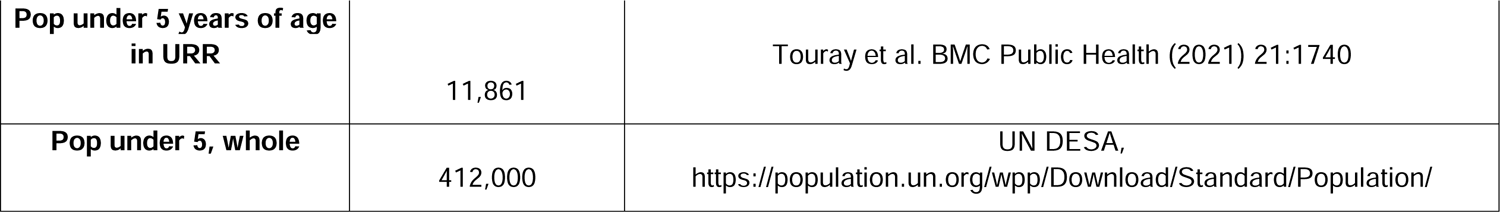
Population estimates for The Gambia and the Upper River Region.

|  | 2013 | 2014 | 2015 | 2016 | 2017 | 2018 | 2019 | 2020 | 2021 |
| --- | --- | --- | --- | --- | --- | --- | --- | --- | --- |
| **Total pop Upper River** | 239,916 | 245,962 | 252,160 | 258,515 | 265,029 | 271,708 | 278,555 | 285,575 | 292,771 |
| **Total male pop Upper River** | 115,960 | 118,882 | 121,878 | 124,949 | 128,098 | 131,326 | 134,636 | 138,028 | 141,507 |
| **Total female pop Upper River** | 123,956 | 127,080 | 130,282 | 133,565 | 136,931 | 140,382 | 143,919 | 147,546 | 151,264 |
| Category | Estimate |  | Source |  |  |  |  |  |  |
| **Total whole pop in 2021** | 2,639,916 |  | World bank data |  |  |  |  |  |  |
| **Annual growth country, whole pop (%)** | 2.52 |  | World bank data |  |  |  |  |  |  |
| **Total female, % of total population** | 50.3 |  | World bank data |  |  |  |  |  |  |
| **Total female pop, whole country** | 1,327,878 |  | World bank data<br>(Calculated from Sourced Data) |  |  |  |  |  |  |

|  |  |  |
| --- | --- | --- |
| **Pop under 5 years of age**<br>**in URR** | 11,861 | Touray et al. BMC Public Health (2021) 21:1740 |
| **Pop under 5, whole** | 412,000 | UN DESA,<br>https://population.un.org/wpp/Download/Standard/Population/ |

**Table S5.**
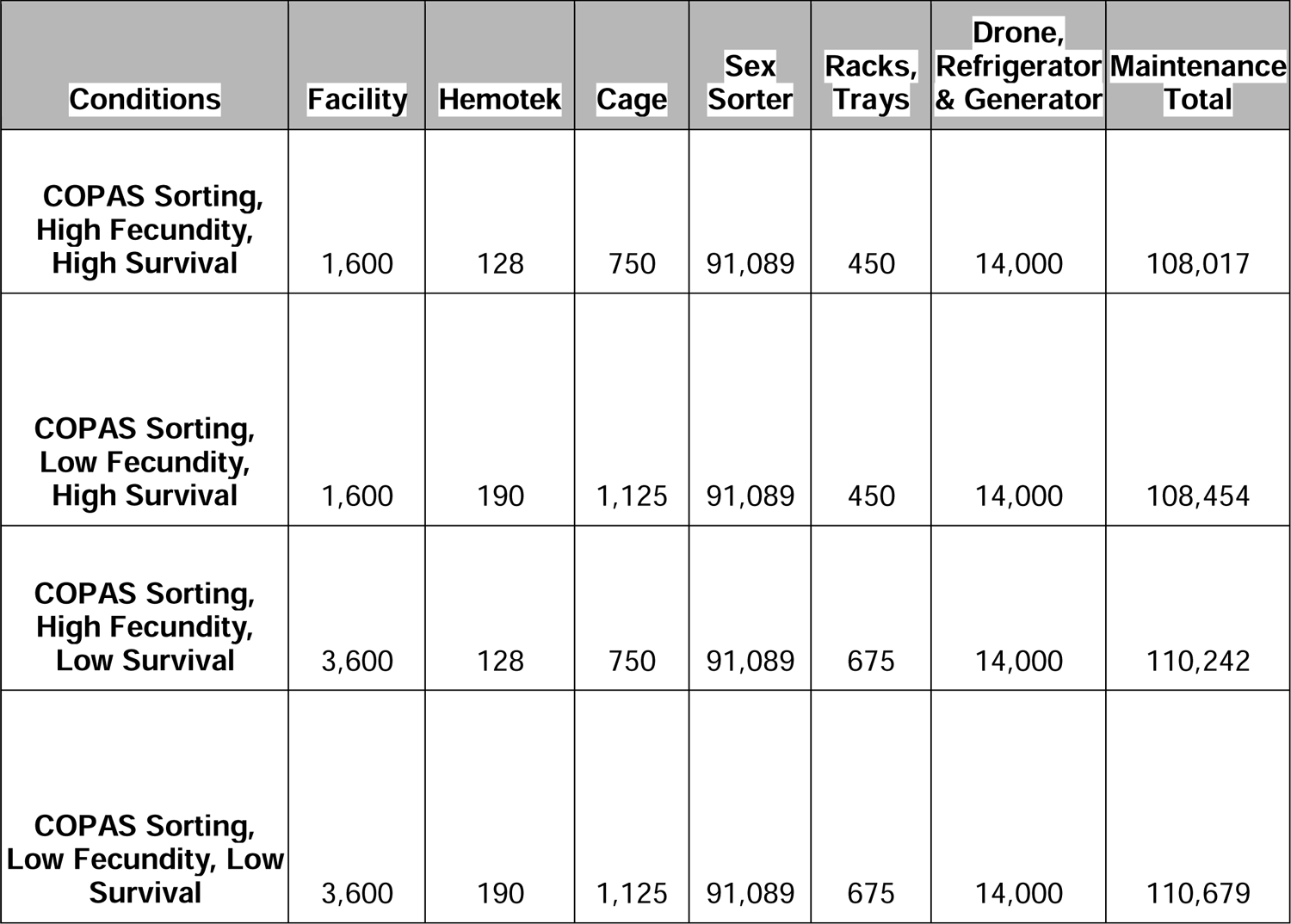
Annual maintenance fees.

| **Conditions** | **Facility** | **Hemotek** | **Cage** | **Sex**<br>**Sorter** | **Racks,**<br>**Trays** | **Drone,**<br>**Refrigerator**<br>**& Generator** | **Maintenance**<br>**Total** |
| --- | --- | --- | --- | --- | --- | --- | --- |
| **COPAS Sorting,**<br>**High Fecundity,**<br>**High Survival** | 1,600 | 128 | 750 | 91,089 | 450 | 14,000 | 108,017 |
| **COPAS Sorting,**<br>**Low Fecundity,**<br>**High Survival** | 1,600 | 190 | 1,125 | 91,089 | 450 | 14,000 | 108,454 |
| **COPAS Sorting,**<br>**High Fecundity,**<br>**Low Survival** | 3,600 | 128 | 750 | 91,089 | 675 | 14,000 | 110,242 |
| **COPAS Sorting,**<br>**Low Fecundity, Low**<br>**Survival** | 3,600 | 190 | 1,125 | 91,089 | 675 | 14,000 | 110,679 |

## Acknowledgments

This work was supported by funding from an Open Philanthropy award (309937-0001). The views, opinions, and/or findings expressed are those of the authors and should not be interpreted as representing the official views or policies of the U.S. government. Some figures were created using www.BioRender.com.

## Disclosures

O.S.A is a founder of Agragene, Inc. and Synvect, Inc. with equity interest. The terms of this arrangement have been reviewed and approved by the University of California, San Diego in accordance with its conflict of interest policies. All other authors declare no competing interests.

## Acronyms and Abbreviations

ABS: acrylonitrile butadiene styrene

ACT: artemisinin-based combination treatment

AI: artificial intelligence

β*2-tubulin*: beta-2 tubulin

Cas9: CRISPR associated protein 9

CHIRPS: Climate Hazards Group InfraRed Precipitation with Station COPAS Complex Object Parametric Analyzer and Sorter

CRISPR: clustered regularly interspaced short palindromic repeats

DNA: deoxyribonucleic acid

dsx: doublesex

GBP: British pound

GDP: gross domestic product

GE: genetically engineered

GMD: Gambian dalasi

HDR: homology directed repair

*fle femaleless*: a female essential gene

gRNA: guide RNA

IAEA: International Atomic Energy Agency

Ifegenia: inherited female elimination by genetically encoded nucleases to interrupt alleles

ICL: Imperial College London

IPTp: intermittent preventive treatment

IRR: internal rate of return

IRS: indoor residual spraying

L1: first instar larvae

LLIN: long last insecticidal nets

LSHTM: London School of Hygiene and Tropical Medicine MGDrivE Mosquito Gene Drive Explorer

MRC: Medical Research Council

NPV: net present value

pgSIT: precision guided sterile insect technique

pgSIT 2.0: pgSIT incorporating SEPARATOR technology to support fluorescent sex specific sorting of the pgSIT lines

PPP: purchasing power parity

RIDL: Release of Insects carrying a Dominant Lethal

RNA: ribonucleic acid

ROI: return on investment

SEPARATOR: Sexing Element Produced by Alternative RNA-splicing of A Transgenic Observable Reporter

SIT: sterile insect technique

SMC: seasonal malaria chemoprevention

QALY: quality adjusted life years

DALY: disability adjusted life years

URR: Upper River region

USD: United States dollar

USTD: United States Transportation Department

VSL: value of statistical life

VTOL: vertical take-off and landing

WHO: World Health Organization

WTP: willingness to pay

zpg: zero population growth

## Notes

### Summary of Updates

Due to a miscommunication, the prior cost-effectiveness assessment targeted mosquito suppression at the peak of the total population rather than targeting the mosquito adult population at the beginning of the rainy season. This resulted in over a 15-fold increase in the number of mosquitoes needed to be suppressed. This update has corrected this and has shown that this intervention is even more cost effective than previously assessed

## References

AAAWater. 2019. “Senegal.” AAAWater. April 4, 2019. https://aaawater.net/nproject/senegal/.

Agyapong, Jeffrey, Joseph Chabi, Aikins Ablorde, Worlasi D. Kartey, Joseph H. N. Osei, Dziedzom K. de Souza, Samuel Dadzie, et al. 2014. “Ovipositional Behavior of Anopheles Gambiae Mosquitoes.” Tropical Medicine and Health 42 (4): 187–90.

Akbari, Omar, Ming Li, Nikolay Kandul, Ruichen Sun, Ting Yang, Elena Dalla Benetta, Daniel Brogan, et al. 2023. “Targeting Sex Determination to Suppress Mosquito Populations.” *Research Square*, April. 10.21203/rs.3.rs-2834069/v1.

Arda. n.d. “Launching Soon.” Arda. Accessed June 7, 2023. https://www.ardaimpact.com/.

Awata, Hiroko, Takahito Watanabe, Yoshitaka Hamanaka, Taro Mito, Sumihare Noji, and Makoto Mizunami. 2015. “Knockout Crickets for the Study of Learning and Memory: Dopamine Receptor Dop1 Mediates Aversive but Not Appetitive Reinforcement in Crickets.” Scientific Reports 5 (November): 15885.

Aweis, Ahmed, Abdinur A. Salad, Fathi A. Araye, Abdifatah M. Ahmed, Osman A. Wehlie, Ali Abdirahman Osman, and Isaiah Gumbe Akuku. 2023. “Long-Lasting Insecticidal Nets (LLINs) Use among Household Members for Protection against Mosquito Bite in Mogadishu Districts.” PLOS Global Public Health 3 (3): e0000724.

Baber, Ibrahima, Moussa Keita, Nafomon Sogoba, Mamadou Konate, M ’bouye Diallo, Seydou Doumbia, Sékou F. Traoré, José M. C. Ribeiro, and Nicholas C. Manoukis. 2010. “Population Size and Migration of Anopheles Gambiae in the Bancoumana Region of Mali and Their Significance for Efficient Vector Control.” PloS One 5 (4): e10270.

Balestrino, Fabrizio, Jérémie R. L. Gilles, Sharon M. Soliban, Anton Nirschl, Quentin E. Benedict, and Mark Q. Benedict. 2011. “Mosquito Mass Rearing Technology: A Cold-Water Vortex Device for Continuous Unattended Separation of Anopheles Arabiensis Pupae from Larvae.” Journal of the American Mosquito Control Association 27 (3): 227–35.

Balestrino, F., M. Q. Benedict, and J. R. L. Gilles. 2012. “A New Larval Tray and Rack System for Improved Mosquito Mass Rearing.” Journal of Medical Entomology 49(3): 595–605.

Bassett, Andrew R., Charlotte Tibbit, Chris P. Ponting, and Ji-Long Liu. 2013. “Highly Efficient Targeted Mutagenesis of Drosophila with the CRISPR/Cas9 System.” Cell Reports 4 (1): 220–28.

Baughman, Ted, Chelsea Peterson, Corrie Ortega, Sarah R. Preston, Christopher Paton, Jessica Williams, Amy Guy, et al. 2017. “A Highly Stable Blood Meal Alternative for Rearing Aedes and Anopheles Mosquitoes.” PLoS Neglected Tropical Diseases 11 (12): e0006142.

Bayoh, M. N., and S. W. Lindsay. 2003. “Effect of Temperature on the Development of the Aquatic Stages of Anopheles Gambiae Sensu Stricto (Diptera: Culicidae).” Bulletin of Entomological Research 93 (5): 375–81.

Bimbilé Somda, Nanwintoum Séverin, Kounbobr Roch Dabiré, Hamidou Maiga, Hanano Yamada, Wadaka Mamai, Olivier Gnankiné, Abdoulaye Diabaté, Antoine Sanon, Jeremy Bouyer, and Jeremie Lionel Gilles. 2017. “Cost-Effective Larval Diet Mixtures for Mass Rearing of Anopheles Arabiensis Patton (Diptera: Culicidae).” Parasites & Vectors 10 (1): 619.

B&R Food Services. n.d. “Beef Blood Frozen 6 Gallon Case American.” Accessed June 8, 2023. https://www.brfood.us/product/beef-blood-frozen-6-gallon-case-american.html.

Broekhuizen, Henk, Alexandra Fehr, Claudia Nieto-Sanchez, Joan Muela, Koen Peeters-Grietens, Tom Smekens, Momodou Kalleh, Esmé Rijndertse, Jane Achan, and Umberto D’Alessandro. 2021. “Costs and Barriers Faced by Households Seeking Malaria Treatment in the Upper River Region, The Gambia.” Malaria Journal 20 (1): 368.

Caraballo, Hector, and Kevin King. 2014. “Emergency Department Management of Mosquito-Borne Illness: Malaria, Dengue, and West Nile Virus.” Emergency Medicine Practice 16 (5): 1–23; quiz 23–24.

Carvalho, Danilo O., Andrew R. McKemey, Luiza Garziera, Renaud Lacroix, Christl A. Donnelly, Luke Alphey, Aldo Malavasi, and Margareth L. Capurro. 2015. “Suppression of a Field Population of Aedes Aegypti in Brazil by Sustained Release of Transgenic Male Mosquitoes.” PLoS Neglected Tropical Diseases 9 (7): e0003864.

Chatzispyrou, Iliana A., Ntsiki M. Held, Laurent Mouchiroud, Johan Auwerx, and Riekelt H. Houtkooper. 2015. “Tetracycline Antibiotics Impair Mitochondrial Function and Its Experimental Use Confounds Research.” Cancer Research 75 (21): 4446–49.

Chikwendu, J. I., A. Onekutu, and I. O. Ogbonna. 2019. “Effects of Host Blood on Fecundity and Longevity of Female Anopheles Mosquitoes.” *International Journal of Pathogen Research*, November, 1–7.

Christiansen-Jucht, Céline D., Paul E. Parham, Adam Saddler, Jacob C. Koella, and María-Gloria Basáñez. 2015. “Larval and Adult Environmental Temperatures Influence the Adult Reproductive Traits of Anopheles Gambiae S.s.” Parasites & Vectors 8 (1): 456.

Christiansen-Jucht, Céline, Paul E. Parham, Adam Saddler, Jacob C. Koella, and María-Gloria Basáñez. 2014. “Temperature during Larval Development and Adult Maintenance Influences the Survival of Anopheles Gambiae S.s.” Parasites & Vectors 7 (November): 489.

Clements, A. N. 1992. The Biology of Mosquitoes. CABI.

Dabira, Edgard D., Harouna M. Soumare, Bakary Conteh, Fatima Ceesay, Mamadou O. Ndiath, John Bradley, Nuredin Mohammed, et al. 2022. “Mass Drug Administration of Ivermectin and Dihydroartemisinin-Piperaquine against Malaria in Settings with High Coverage of Standard Control Interventions: A Cluster-Randomised Controlled Trial in The Gambia.” The Lancet Infectious Diseases.

Damiens, D., S. M. Soliban, F. Balestrino, R. Alsir, M. J. B. Vreysen, and J. R. L. Gilles. 2013. “Different Blood and Sugar Feeding Regimes Affect the Productivity of Anopheles Arabiensis Colonies (Diptera: Culicidae).” Journal of Medical Entomology 50 (2): 336–43.

“Departmental Guidance on Valuation of a Statistical Life in Economic Analysis.” n.d. Accessed January 12, 2024. https://www.transportation.gov/office-policy/transportation-policy/revised-departmental-guidance-on-valuation-of-a-statistical-life-in-economic-analysis.

Dieng, Idrissa, Marie Henriette Dior Ndione, Cheikh Fall, Moussa Moïse Diagne, Mamadou Diop, Aboubacry Gaye, Mamadou Aliou Barry, et al. 2021. “Multifoci and Multiserotypes Circulation of Dengue Virus in Senegal between 2017 and 2018.” BMC Infectious Diseases 21 (1): 867.

Diepeveen, Stephanie, Tom Ling, Marc Suhrcke, Martin Roland, and Theresa M. Marteau. 2013. “Public Acceptability of Government Intervention to Change Health-Related Behaviours: A Systematic Review and Narrative Synthesis.” BMC Public Health 13 (August): 756.

Faiman, Roy, Alpha S. Yaro, Moussa Diallo, Adama Dao, Samake Djibril, Zana L. Sanogo, Margery Sullivan, Asha Krishna, Benjamin J. Krajacich, and Tovi Lehmann. 2020. “Quantifying Flight Aptitude Variation in Wild Anopheles Gambiae in Order to Identify Long-Distance Migrants.” Malaria Journal 19 (1): 263.

FAO/IAEA. 2017. “Guidelines for Standardised Mass Rearing of Anopheles Mosquitoes—Version 1.0.” FAO Rome, Italy. https://www.iaea.org/resources/manual/guidelines-for-standardised-mass-rearing-of-anopheles-mosquitoes-version-10.

Framework, Privacy Shield. n.d. “Gambia Country Commercial Guide: Gambia - Distribution & Sales Channels.” Accessed June 29, 2023. https://www.privacyshield.gov/article?id=Gambia-Distribution-Sales-Channels.

Fru, Paulette Ngum, Frederick Nchang Cho, Andrew N. Tassang, Celestina Neh Fru, Peter Nde Fon, and Albert Same Ekobo. 2021. “Ownership and Utilisation of Long-Lasting Insecticidal Nets in Tiko Health District, Southwest Region, Cameroon: A Cross-Sectional Study.” *Journal of Parasitology Research* 2021 (February): 8848091.

Gallup, J. L., and J. D. Sachs. 2001. “The Economic Burden of Malaria.” The American Journal of Tropical Medicine and Hygiene 64 (1-2 Suppl): 85–96.

Gambia Bureau of Statistics. 2015. “Population and Housing Census of Gambia, 2013.” Gambia Data Portal. June 1, 2015. https://gambia.opendataforafrica.org/mmfoqkd/population-and-housing-census-of-gambia-2013.

Gantz, Valentino M., Nijole Jasinskiene, Olga Tatarenkova, Aniko Fazekas, Vanessa M. Macias, Ethan Bier, and Anthony A. James. 2015. “Highly Efficient Cas9-Mediated Gene Drive for Population Modification of the Malaria Vector Mosquito Anopheles Stephensi.” Proceedings of the National Academy of Sciences of the United States of America 112 (49): E6736–43.

Garziera, Luiza, Michelle Cristine Pedrosa, Fabrício Almeida de Souza, Maylen Gómez, Márcia Bento Moreira, Jair Fernandes Virginio, Margareth Lara Capurro, and Danilo Oliveira Carvalho. 2017. “Effect of Interruption of OverlJflooding Releases of Transgenic Mosquitoes over Wild Population of *Aedes Aegypti*: Two Case Studies in Brazil.” Entomologia Experimentalis et Applicata 164 (3): 327–39.

Gayler-Thompson, Georgie. 2023. “Oxitec Announces Expansion into Central America to Launch New Program in Panama to Fight Malaria-Transmitting Mosquitoes Threatening Meso-America.” Oxitec. April 27, 2023. https://www.oxitec.com/en/news/oxitec-announces-expansion-into-panama.

Gerhard Brümmer, Lucy McLane, Elisa Campos, Thelma Hlatshwayo. 2022. 2022/23 Property & Construction Africa Cost Guide Handbook. AECOM.com.

Gratz, Scott J., Alexander M. Cummings, Jennifer N. Nguyen, Danielle C. Hamm, Laura K. Donohue, Melissa M. Harrison, Jill Wildonger, and Kate M. O’Connor-Giles. 2013. “Genome Engineering of Drosophila with the CRISPR RNA-Guided Cas9 Nuclease.” Genetics 194 (4): 1029–35.

Griffin, Jamie T., Samir Bhatt, Marianne E. Sinka, Peter W. Gething, Michael Lynch, Edith Patouillard, Erin Shutes, et al. 2016. “Potential for Reduction of Burden and Local Elimination of Malaria by Reducing Plasmodium Falciparum Malaria Transmission: A Mathematical Modelling Study.” The Lancet Infectious Diseases 16 (4): 465–72.

Griffin, Jamie T., T. Deirdre Hollingsworth, Lucy C. Okell, Thomas S. Churcher, Michael White, Wes Hinsley, Teun Bousema, et al. 2010. “Reducing Plasmodium Falciparum Malaria Transmission in Africa: A Model-Based Evaluation of Intervention Strategies.” PLoS Medicine 7 (8). 10.1371/journal.pmed.1000324.

Halliday Sarah. 2017. “Oxitec and GBIT Announce Launch of Friendly^TM^ Aedes Project in India.” Oxitec. January 23, 2017. https://www.oxitec.com/en/news/oxitec-and-gbit-announce-launch-of-friendly-aedes-project-in-india.

Halliday Sarah. 2018. “The Cayman Government and Oxitec Launch Innovative Pilot to Suppress Aedes Aegypti.” Oxitec. May 22, 2018. https://www.oxitec.com/en/news/the-cayman-government-and-oxitec-launch-innovative-pilot-to-suppress-aedes-aegypti.

Hamel, Mary J., Peter Otieno, Nabie Bayoh, Simon Kariuki, Vincent Were, Doris Marwanga, Kayla F. Laserson, John Williamson, Laurence Slutsker, and John Gimnig. 2011. “The Combination of Indoor Residual Spraying and Insecticide-Treated Nets Provides Added Protection against Malaria Compared with Insecticide-Treated Nets Alone.” The American Journal of Tropical Medicine and Hygiene 85 (6): 1080–86.

Hammond, Andrew, Roberto Galizi, Kyros Kyrou, Alekos Simoni, Carla Siniscalchi, Dimitris Katsanos, Matthew Gribble, et al. 2015. “A CRISPR-Cas9 Gene Drive System Targeting Female Reproduction in the Malaria Mosquito Vector Anopheles Gambiae.” Nature Biotechnology 34 (December): 78.

Health Organization, World. n.d. “Guidance Framework for Testing Genetically Modified Mosquitoes.” Accessed June 16, 2023. https://apps.who.int/iris/bitstream/handle/10665/341370/9789240025233-eng.pdf?sequence=1.

Hemingway, Janet. 2014. “The Role of Vector Control in Stopping the Transmission of Malaria: Threats and Opportunities.” Philosophical Transactions of the Royal Society of London. Series B, Biological Sciences 369 (1645): 20130431.

Horton, Susan. 2017. “Cost-Effectiveness Analysis in Disease Control Priorities.” Disease Control Priorities: Improving Health and Reducing Poverty. 3rd Edition. https://www.ncbi.nlm.nih.gov/books/NBK525287/.

Huang, Yuping, Yazhou Chen, Baosheng Zeng, Yajun Wang, Anthony A. James, Geoff M. Gurr, Guang Yang, Xijian Lin, Yongping Huang, and Minsheng You. 2016. “CRISPR/Cas9 Mediated Knockout of the Abdominal-A Homeotic Gene in the Global Pest, Diamondback Moth (Plutella Xylostella).” Insect Biochemistry and Molecular Biology 75 (August): 98–106.

Huestis, Diana L., Adama Dao, Moussa Diallo, Zana L. Sanogo, Djibril Samake, Alpha S. Yaro, Yossi Ousman, et al. 2019. “Windborne Long-Distance Migration of Malaria Mosquitoes in the Sahel.” Nature 574 (7778): 404–8.

Info, Worlddata. 2021. “Telecommunication in the Gambia.” Worlddata.info. 2021. https://www.worlddata.info/africa/gambia/telecommunication.php.

Kalghatgi, Sameer, Catherine S. Spina, James C. Costello, Marc Liesa, J. Ruben Morones-Ramirez, Shimyn Slomovic, Anthony Molina, Orian S. Shirihai, and James J. Collins. 2013. “Bactericidal Antibiotics Induce Mitochondrial Dysfunction and Oxidative Damage in Mammalian Cells.” Science Translational Medicine 5 (192): 192ra85.

Kandul, Nikolay P., Junru Liu, and Omar S. Akbari. 2021. “Temperature-Inducible Precision-Guided Sterile Insect Technique.” The CRISPR Journal 4 (6): 822–35.

Kandul, Nikolay P., Junru Liu, Anna Buchman, Isaiah C. Shriner, Rodrigo M. Corder, Natalie Warsinger-Pepe, Ting Yang, et al. 2022. “Precision Guided Sterile Males Suppress Populations of an Invasive Crop Pest.” GEN Biotechnology 1 (4): 372– 85.

Kandul, Nikolay P., Junru Liu, Hector M., Sanchez C, Sean L. Wu, John M. Marshall, and Omar S. Akbari. 2019. “Transforming Insect Population Control with Precision Guided Sterile Males with Demonstration in Flies.” Nature Communications 10 (1): 84.

Kandul, Nikolay P., Junru Liu, Hector M. Sanchez, Sean L. Wu, John M. Marshall, and Omar S. Akbari. 2018. “Transforming Insect Population Control with Precision Guided Sterile Males.” bioRxiv. 10.1101/377721.

Kayedi, M. H., J. D. Lines, A. A. Haghdoost, and S. Najafi. 2007. “A Randomized and Controlled Comparison of the Wash-Resistances and Insecticidal Efficacies of Four Types of Deltamethrin-Treated Nets, over a 6-Month Period of Domestic Use with Washing Every 2 Weeks, in a Rural Area of Iran.” Annals of Tropical Medicine and Parasitology 101 (6): 519–28.

Kip Viscusi, W., and Clayton J. Masterman. 2017. “Income Elasticities and Global Values of a Statistical Life.” Journal of Benefit-Cost Analysis 8 (2): 226–50.

Kistler, Kathryn E., Leslie B. Vosshall, and Benjamin J. Matthews. 2015. “Genome Engineering with CRISPR-Cas9 in the Mosquito Aedes Aegypti.” Cell Reports 11 (1): 51–60.

Lacroix, Renaud, Andrew R. McKemey, Norzahira Raduan, Lim Kwee Wee, Wong Hong Ming, Teoh Guat Ney, Siti Rahidah A A, et al. 2012. “Open Field Release of Genetically Engineered Sterile Male Aedes Aegypti in Malaysia.” PloS One 7 (8): e42771.

Li, Ming, Ting Yang, Michelle Bui, Stephanie Gamez, Tyler Wise, Nikolay P. Kandul, Junru Liu, et al. 2021. “Suppressing Mosquito Populations with Precision Guided Sterile Males.” Nature Communications 12 (1): 5374.

Li, Xueyan, Dingding Fan, Wei Zhang, Guichun Liu, Lu Zhang, Li Zhao, Xiaodong Fang, et al. 2015. “Outbred Genome Sequencing and CRISPR/Cas9 Gene Editing in Butterflies.” Nature Communications 6 (September): 8212.

Li, Yan, Jie Zhang, Dafeng Chen, Pengcheng Yang, Feng Jiang, Xianhui Wang, and Le Kang. 2016. “CRISPR/Cas9 in Locusts: Successful Establishment of an Olfactory Deficiency Line by Targeting the Mutagenesis of an Odorant Receptor Co-Receptor (Orco).” Insect Biochemistry and Molecular Biology 79 (December): 27–35.

Long, Kanya C., Luke Alphey, George J. Annas, Cinnamon S. Bloss, Karl J. Campbell, Jackson Champer, Chun-Hong Chen, et al. 2020. “Core Commitments for Field Trials of Gene Drive Organisms.” Science 370 (6523): 1417–19.

Lukwa, Akim Tafadzwa, Richard Mawoyo, Karen Nelwin Zablon, Aggrey Siya, and Olufunke Alaba. 2019. “Effect of Malaria on Productivity in a Workplace: The Case of a Banana Plantation in Zimbabwe.” Malaria Journal 18 (1): 390.

Lyimo, I. N., S. P. Keegan, L. C. Ranford-Cartwright, and H. M. Ferguson. 2012. “The Impact of Uniform and Mixed Species Blood Meals on the Fitness of the Mosquito Vector Anopheles Gambiae S.s: Does a Specialist Pay for Diversifying Its Host Species Diet?” Journal of Evolutionary Biology 25 (3): 452–60.

Maïga, Hamidou, Wadaka Mamai, Nanwintoum Séverin Bimbilé Somda, Anna Konczal, Thomas Wallner, Gustavo Salvador Herranz, Rafael Argiles Herrero, Hanano Yamada, and Jeremy Bouyer. 2019. “Reducing the Cost and Assessing the Performance of a Novel Adult Mass-Rearing Cage for the Dengue, Chikungunya, Yellow Fever and Zika Vector, Aedes Aegypti (Linnaeus).” PLoS Neglected Tropical Diseases 13 (9): e0007775.

Marois, Eric, Christina Scali, Julien Soichot, Christine Kappler, Elena A. Levashina, and Flaminia Catteruccia. 2012. “High-Throughput Sorting of Mosquito Larvae for Laboratory Studies and for Future Vector Control Interventions.” Malaria Journal. 10.1186/1475-2875-11-302.

Ma, Sanyuan, Jiasong Chang, Xiaogang Wang, Yuanyuan Liu, Jianduo Zhang, Wei Lu, Jie Gao, Run Shi, Ping Zhao, and Qingyou Xia. 2014. “CRISPR/Cas9 Mediated Multiplex Genome Editing and Heritable Mutagenesis of BmKu70 in Bombyx Mori.” Scientific Reports 4 (March): 4489.

Masum, Hassan, Ronak Shah, Karl Schroeder, Abdallah S. Daar, and Peter A. Singer. 2010. “Africa’s Largest Long-Lasting Insecticide-Treated Net Producer: Lessons from A to Z Textiles.” BMC International Health and Human Rights 10 Suppl 1 (Suppl 1): S6.

Mazigo, Ernest, Winifrida Kidima, Joseph Myamba, and Eliningaya J. Kweka. 2019. “The Impact of Anopheles Gambiae Egg Storage for Mass Rearing and Production Success.” Malaria Journal 18 (1): 52.

Mondal, Agastya, Héctor M., Sánchez C, and John M. Marshall. 2023. “MGDrivE 3: A Decoupled Vector-Human Framework for Epidemiological Simulation of Mosquito Genetic Control Tools and Their Surveillance.” *bioRxiv: The Preprint Server for Biology*, September. 10.1101/2023.09.09.556958.

“More Malaria Cases and Deaths in 2020 Linked to COVID-19 Disruptions.” n.d. Accessed May 31, 2023. https://www.who.int/news/item/06-12-2021-more-malaria-cases-and-deaths-in-2020-linked-to-covid-19-disruptions.

Moullan, Norman, Laurent Mouchiroud, Xu Wang, Dongryeol Ryu, Evan G. Williams, Adrienne Mottis, Virginija Jovaisaite, et al. 2015. “Tetracyclines Disturb Mitochondrial Function across Eukaryotic Models: A Call for Caution in Biomedical Research.” Cell Reports, March. 10.1016/j.celrep.2015.02.034.

Muriu, Simon M., Tim Coulson, Charles M. Mbogo, and H. Charles, J. Godfray. 2013. “Larval Density Dependence in Anopheles Gambiae S.s., the Major African Vector of Malaria.” The Journal of Animal Ecology 82 (1): 166–74.

Musoke, David, Edwinah Atusingwize, Carol Namata, Rawlance Ndejjo, Rhoda K. Wanyenze, and Moses R. Kamya. 2023. “Integrated Malaria Prevention in Low- and Middle-Income Countries: A Systematic Review.” Malaria Journal 22 (1): 79.

Njie, Hassan, Knut Reidar Wangen, Lumbwe Chola, Unni Gopinathan, Ibrahimu Mdala, Johanne S. Sundby, and Patrick G. C. Ilboudo. 2023. “Willingness to Pay for a National Health Insurance Scheme in The Gambia: A Contingent Valuation Study.” Health Policy and Planning 38 (1): 61–73.

Papathanos, Philippos A., Hervé C. Bossin, Mark Q. Benedict, Flaminia Catteruccia, Colin A. Malcolm, Luke Alphey, and Andrea Crisanti. 2009. “Sex Separation Strategies: Past Experience and New Approaches.” Malaria Journal 8 Suppl 2 (November): S5.

Pare Toe, Lea, Nourou Barry, Anselme D. Ky, Souleymane Kekele, Wilfrid Meda, Korotimi Bayala, Mouhamed Drabo, Delphine Thizy, and Abdoulaye Diabate. 2021. “Small-Scale Release of Non-Gene Drive Mosquitoes in Burkina Faso: From Engagement Implementation to Assessment, a Learning Journey.” Malaria Journal 20 (1): 395.

Pfeffer, Daniel A., Timothy C. D. Lucas, Daniel May, Joseph Harris, Jennifer Rozier, Katherine A. Twohig, Ursula Dalrymple, et al. 2018. “malariaAtlas: An R Interface to Global Malariometric Data Hosted by the Malaria Atlas Project.” Malaria Journal 17(1): 352.

Pryce, Joseph, Nancy Medley, and Leslie Choi. 2022. “Indoor Residual Spraying for Preventing Malaria in Communities Using Insecticide-Treated Nets.” Cochrane Database of Systematic Reviews 1 (1): CD012688.

Rashid, Iliyas, Melina Campos, Travis Collier, Marc Crepeau, Allison Weakley, Hans Gripkey, Yoosook Lee, Hanno Schmidt, and Gregory C. Lanzaro. 2022. “Spontaneous Mutation Rate Estimates for the Principal Malaria Vectors Anopheles Coluzzii and Anopheles Stephensi.” Scientific Reports 12 (1): 226.

Reid, William, and David A. O’Brochta. 2016. “Applications of Genome Editing in Insects.” Current Opinion in Insect Science 13 (February): 43–54.

Sachs, Jeffrey, and Pia Malaney. 2002. “The Economic and Social Burden of Malaria.” Nature 415 (6872): 680–85.

Sánchez C, Héctor M., Sean L. Wu, Jared B. Bennett, and John M. Marshall. 2020. “MGDrivE: A Modular Simulation Framework for the Spread of Gene Drives through Spatially Explicit Mosquito Populations.” Methods in Ecology and Evolution / British Ecological Society 11 (2): 229–39.

Sanogo, Zana L., Alpha S. Yaro, Adama Dao, Moussa Diallo, Ousman Yossi, Djibril Samaké, Benjamin J. Krajacich, Roy Faiman, and Tovi Lehmann. 2021. “The Effects of High-Altitude Windborne Migration on Survival, Oviposition, and Blood-Feeding of the African Malaria Mosquito, Anopheles Gambiae S.l. (Diptera: Culicidae).” Journal of Medical Entomology 58 (1): 343–49.

Schairer, Cynthia E., James Najera, Anthony A. James, Omar S. Akbari, and Cinnamon S. Bloss. 2021. “Oxitec and MosquitoMate in the United States: Lessons for the Future of Gene Drive Mosquito Control.” Pathogens and Global Health 115 (6): 365–76.

Shretta, Rima, Jenny Liu, Chris Cotter, Justin Cohen, Charlotte Dolenz, Kudzai Makomva, Gretchen Newby, et al. 2017. Malaria Elimination and Eradication. The International Bank for Reconstruction and Development / The World Bank.

Smidler, Andrea L., Reema A. Apte, James J. Pai, Martha L. Chow, Sanle Chen, Agastya Mondal, Héctor M., Sánchez C, Igor Antoshechkin, John M. Marshall, and Omar S. Akbari. 2023. “Eliminating Malaria Vectors with Precision Guided Sterile Males.” *bioRxiv: The Preprint Server for Biology*, July. 10.1101/2023.07.20.549947.

Smidler, Andrea L., James J. Pai, Reema A. Apte, Héctor M., Sánchez C, Rodrigo M. Corder, Eileen Jeffrey Gutiérrez, Neha Thakre, Igor Antoshechkin, John M. Marshall, and Omar S. Akbari. 2023. “A Confinable Female-Lethal Population Suppression System in the Malaria Vector,.” Science Advances 9 (27): eade8903.

Soumare, Harouna M., Edgard Diniba Dabira, Muhammed M. Camara, Lamin Jadama, Pa Modou Gaye, Sainey Kanteh, Ebrima A. Jawara, et al. 2022. “Entomological Impact of Mass Administration of Ivermectin and Dihydroartemisinin-Piperaquine in The Gambia: A Cluster-Randomized Controlled Trial.” Parasites & Vectors 15 (1): 435.

de Swart, Marieke M., Carlijn Balvers, Niels O. Verhulst, and Constantianus J. M. Koenraadt. 2023. “Effects of Host Blood on Mosquito Reproduction.” *Trends in Parasitology*, May. 10.1016/j.pt.2023.04.003.

Takken, W., M. J. Klowden, and G. M. Chambers. 1998. “Effect of Body Size on Host Seeking and Blood Meal Utilization in Anopheles Gambiae Sensu Stricto (Diptera: Culicidae): The Disadvantage of Being Small.” Journal of Medical Entomology 35 (5): 639–45.

targetmalaria.org. 2020. “Anopheles Gambiae Sl: Morphology, Life-Cycle, Ecology.” August 2020. https://targetmalaria.org/wp-content/uploads/2020/11/Ecology_FS_EN_Anopheles-gambiae-s.l.-morphology_August20.pdf.

“The Cost of Building in Africa.” 2022. RLB | Americas. Rider Levett Bucknall. April 9, 2022. https://www.rlb.com/americas/insight/perspective-2022-vol-1/the-cost-of-building-in-africa/.

Thomas, D. D., C. A. Donnelly, R. J. Wood, and L. S. Alphey. 2000. “Insect Population Control Using a Dominant, Repressible, Lethal Genetic System.” Science 287 (5462): 2474–76.

Trapero-Bertran, M., H. Mistry, J. Shen, and J. Fox-Rushby. 2013. “A Systematic Review and Meta-Analysis of Willingness-to-Pay Values: The Case of Malaria Control Interventions.” Health Economics 22 (4): 428–50.

Unicef, and Others. 2020. “Long-Lasting Insecticidal Nets: Supply Update UNICEF Supply Division. United Nations International ChildrenlJs Emergency Fund, New York, USA.”

“U.S. Department of State.” n.d. Accessed June 6, 2023. https://aoprals.state.gov/web920/hardship.asp.

Vanness, David J., James Lomas, and Hannah Ahn. 2021. “A Health Opportunity Cost Threshold for Cost-Effectiveness Analysis in the United States.” Annals of Internal Medicine 174 (1): 25–32.

“Vector-Borne Diseases.” n.d. Accessed August 19, 2022. https://www.who.int/news-room/fact-sheets/detail/vector-borne-diseases.

Viscusi, W. Kip. 2020. “Pricing the Global Health Risks of the COVID-19 Pandemic.” Journal of Risk and Uncertainty 61 (2): 101–28.

Waltz, Emily. 2021. “First Genetically Modified Mosquitoes Released in the United States.” Nature 593 (7858): 175–76.

Weng, Shih-Che, Igor Antoshechkin, Eric Marois, and Omar S. Akbari. 2023. “Efficient Sex Separation by Exploiting Differential Alternative Splicing of a Dominant Marker in Aedes Aegypti.” bioRxiv. 10.1101/2023.06.16.545348.

White, Michael T., Jamie T. Griffin, Thomas S. Churcher, Neil M. Ferguson, María-Gloria Basáñez, and Azra C. Ghani. 2011. “Modelling the Impact of Vector Control Interventions on Anopheles Gambiae Population Dynamics.” Parasites & Vectors 4 (July): 153.

“WHO Recommends R21/Matrix-M Vaccine for Malaria Prevention in Updated Advice on Immunization.” n.d. Accessed December 12, 2023. https://www.who.int/news/item/02-10-2023-who-recommends-r21-matrix-m-vaccine-for-malaria-prevention-in-updated-advice-on-immunization.

Wolbaki. n.d. “Wolbaki-Mosquito Larvae Mass Rearing Equipment.” Accessed June 7, 2023. https://wolbaki.com/enservice/114.html.

Wubishet, Mesfin Kelkile, Gebretsadik Berhe, Alefech Adissu, and Mesfin Segni Tafa. 2021. “Effectiveness of Long-Lasting Insecticidal Nets in Prevention of Malaria among Individuals Visiting Health Centres in Ziway-Dugda District, Ethiopia: Matched Case-Control Study.” Malaria Journal 20 (1): 301.

Wu, Sean L., Jared B. Bennett, Héctor M., Sánchez C, Andrew J. Dolgert, Tomás M. León, and John M. Marshall. 2021. “MGDrivE 2: A Simulation Framework for Gene Drive Systems Incorporating Seasonality and Epidemiological Dynamics.” PLoS Computational Biology 17 (5): e1009030.

Yao, Franck Adama, Abdoul-Azize Millogo, Patric Stephane Epopa, Ace North, Florian Noulin, Koulmaga Dao, Mouhamed Drabo, et al. 2022. “Mark-Release-Recapture Experiment in Burkina Faso Demonstrates Reduced Fitness and Dispersal of Genetically-Modified Sterile Malaria Mosquitoes.” Nature Communications 13 (1): 796.

Yaro, Alpha S., Adama Dao, Abdoulaye Adamou, Jacob E. Crawford, José M. C. Ribeiro, Robert Gwadz, Sekou F. Traoré, and Tovi Lehmann. 2006. “The Distribution of Hatching Time in Anopheles Gambiae.” Malaria Journal 5 (March): 19.

Yaro, A. S., A. Dao, A. Adamou, J. E. Crawford, S. F. Traoré, A. M. Touré, R. Gwadz, and T. Lehmann. 2006. “Reproductive Output of Female Anopheles Gambiae (Diptera: Culicidae): Comparison of Molecular Forms.” Journal of Medical Entomology 43 (5): 833–39.

